# Ovarian hormones mediate the prophylactic efficacy of (*R,S*)-ketamine and (2*R*,6*R*)-hydroxynorketamine in female mice

**DOI:** 10.1101/712752

**Authors:** Briana K. Chen, Christina T. LaGamma, Xiaoming Xu, Shi-Xian Deng, Rebecca A. Brachman, Raymond F. Suckow, Thomas B. Cooper, Donald W. Landry, Christine A. Denny

## Abstract

**BACKGROUND:** Females are more likely than males to develop major depressive disorder (MDD) after exposure to stress. We previously reported that the administration of (*R,S*)-ketamine before stress can prevent stress-induced depressive-like behavior in male mice but have yet to assess efficacy in female mice or for other compounds, such as the metabolites of (*R,S*)-ketamine.

**METHODS:** We administered (*R,S*)-ketamine or its metabolites (2*R*,6*R*)-hydroxynorketamine ((2*R*,6*R*)-HNK) and (2*S*,6*S*)-HNK at various doses 1 week before one of a number of stressors, including contextual fear conditioning (CFC), learned helplessness (LH), and chronic immobilization stress (CIS), in male and female 129S6/SvEv mice. To examine the interaction between ovarian hormones and stress resilience, female mice also underwent ovariectomy surgery (OVX) and a hormone replacement protocol prior to drug administration.

**RESULTS:** (*R,S*)-ketamine and (2*S*,6*S*)-HNK, but not (2*R*,6*R*)-HNK, attenuated learned fear in male mice. (*R,S*)-ketamine and (2*R*,6*R*)-HNK, but not (2*S*,6*S*)-HNK, significantly reduced stress-induced depressive-like behavior in male and female mice. (*R,S*)-ketamine and (2*R*,6*R*)-HNK) were prophylactically effective at a lower dose (10 mg/kg and 0.025 mg/kg, respectively) in female mice than in male mice (30 mg/kg and 0.075 mg/kg, respectively). Moreover, ovarian-derived hormones were necessary and sufficient for prophylaxis in female mice.

**CONCLUSIONS:** Our results suggest that prophylactics against stress-induced depressive-like behavior can be developed in a sex-specific manner and that ovarian hormones mediate prophylactic efficacy in females. To our knowledge, this is the first demonstration of the prophylactic efficacy of the metabolites of (*R,S*)-ketamine in male and female mice.

## INTRODUCTION

Major depressive disorder (MDD) is the leading cause of disability worldwide, affecting over 300 million people, and results from social, psychological, and biological factors (1). In 80% of cases, a traumatic event triggers the first major depressive episode, after which symptoms persist throughout an individual’s lifetime (2). MDD has a high comorbidity rate with other psychiatric disorders, including post-traumatic stress disorder (PTSD); approximately half of patients suffering from PTSD are concurrently diagnosed with depression, the majority of which are women (3-5). Regardless of age or socioeconomic status, women are twice as likely as men to be diagnosed with depression and develop MDD earlier in life (1, 5). fMRI data suggest that women experience fear more strongly than men and process trauma through distinct brain circuits (6). Furthermore, certain antidepressants are less effective in women than in men, and changes in hormone levels can influence antidepressant efficacy in women (7, 8). Given these sex-specific differences, it is necessary to develop more specific and efficacious treatments for female populations.

MDD is treated using a combination of therapy and pharmacological antidepressants (9). However, these medications are slow to provide relief and fail to treat up to 30% of patients (9). These drawbacks have led to the use of (*R,S*)-ketamine, a commonly-used anesthetic, as a rapid-acting antidepressant for treatment-resistant MDD (TRD) (10, 11). When given at sub-anesthetic doses, (*R,S*)-ketamine has a rapid antidepressant onset of 2 hours in humans and 30 minutes in mice, can last up to 2 weeks, and acts in a sex-specific manner (12-15). Controlling for weight, females are more sensitive than males to (*R,S*)-ketamine, requiring a lower concentration to reverse depressive-like behaviors, and doses beneficial to males are depressogenic and anxiogenic in females (14). These findings underscore the need for further sex-specific investigation into the use of (*R,S*)-ketamine in MDD treatment.

Because (*R,S*)-ketamine has a wide range of biological targets, isolating the specific mechanisms underlying its antidepressant actions has remained elusive (16). (*R,S*)-ketamine is stereoselectively metabolized into a variety of compounds, including (*R,S*)-norketamine and (2*R*,6*R*;2*S*,6*S*)-hydroxynorketamine (HNK) (17). (2*R*,6*R*;2*S*,6*S*)-HNK is a major metabolite found in the brain and plasma following (*R,S*)-ketamine infusion, comprising 15% of the original (*R,S*)-ketamine dose (18). Although some studies suggest that the *R* enantiomer of HNK, (2*R*,6*R*)-HNK, may be necessary for the antidepressant effects of racemic (*R,S*)-ketamine, the data remain unclear (19-23). Thus, further investigation is needed to determine whether the (*S*)- or (*R*)-enantiomer contribute to (*R,S*)-ketamine’s antidepressant effects and whether (*R,S*)-ketamine metabolites are an effective treatment for TRD.

In addition to (*R,S*)-ketamine’s actions in treating MDD, recent research indicate that (*R,S*)-ketamine could be used to *prevent* stress-induced depression before it develops. Our lab and others have shown that a single injection of (*R,S*)-ketamine before stress can protect against the onset of stress-induced depressive-like behavior and social avoidance as well as attenuate learned fear, suggesting the possibility of developing resilience-enhancing pharmacotherapy (24-27). However, it is still unknown whether stereospecific (*R,S*)-ketamine metabolites can have the same prophylactic efficacy of their racemic precursor.

Here, we investigated which metabolites of (*R,S*)-ketamine were driving prophylactic efficacy and whether the invidual metabolites could have the same prophylactic efficacy as (*R,S*)-ketamine in male and female mice. (*R,S*)-ketamine and (2*S*,6*S*)-HNK, but not (2*R*,6*R*)-HNK, attenuated learned fear in male mice. (*R,S*)-ketamine and (2*R*,6*R*)-HNK, but not (2*S*,6*S*)-HNK, significantly reduced stress-induced depressive-like behavior in male and female mice. (*R,S*)-ketamine and (2*R*,6*R*)-HNK) were prophylactically effective at a lower dose in female mice than in male mice. Moreover, ovarian-derived hormones were necessary and sufficient for prophylaxis in female mice. These data demonstrate that prophylactics against depressive-like behavior can be developed for use in females and emphasize the need for sex-specific approaches to the prevention and treatment of psychiatric disorders in future studies.

## METHODS AND MATERIALS

### Mice

Male and female 129S6/SvEvTac mice were purchased from Taconic (Hudson, NY) at 7 weeks of age. Mice were housed 5 per cage in a 12-h (06:00-18:00) light-dark colony room at 22°C. Food and water were provided *ad libitum*. Behavioral testing was performed during the light phase. All experiments were approved by the Institutional Animal Care and Use Committee (IACUC) at the New York Psychiatric Institute (NYSPI).

## RESULTS

### (2*R*,6*R*)-HNK and (2*S*,6*S*)-HNK protect against distinct stress-induced behaviors in male 129S6/SvEv mice

Male 129S6/SvEv mice were administered a single injection of saline, (*R,S*)-ketamine, (2*R*,6*R*)-HNK, or (2*S*,6*S*)-HNK at a range of doses 1 week prior to 3-shock CFC. Five days later, mice were then re-exposed to the CFC context and administered the FST (**Figure 1A**).

**Figure 1.**
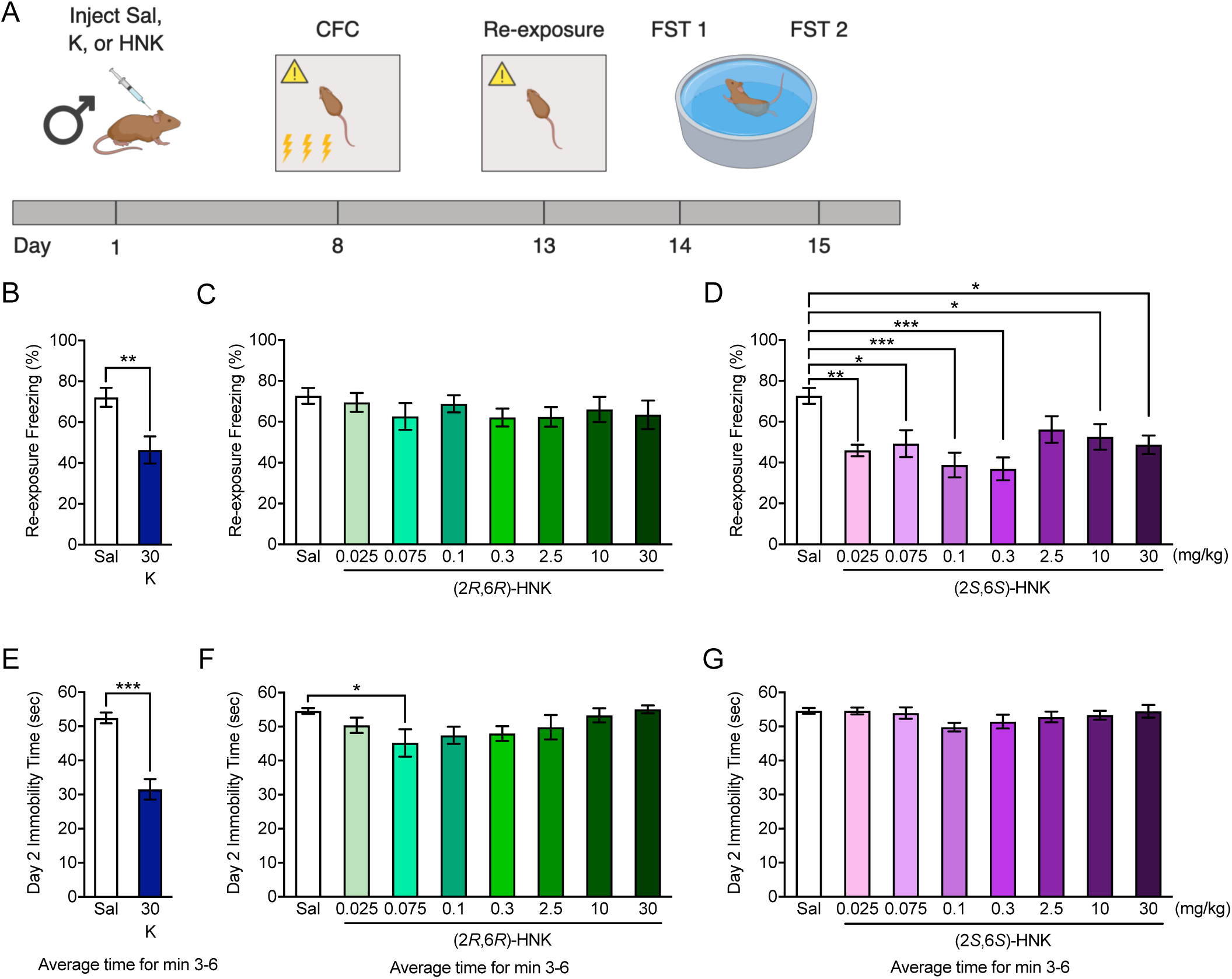
Prophylactic (*R,S*)-ketamine, (2*R*,6*R*)-HNK, and (2*S*,6*S*)-HNK differentially protect against stress in male 129S6/SvEv mice. **(A)** Experimental design. **(B), (C), (D)** (*R,S*)-ketamine (30 mg/kg) and (2*S*,6*S*)-HNK (0.025, 0.075, 0.1, 0.3, 10, and 30 mg/kg), but not (2*R*,6*R*)-HNK, attenuated learned in male mice. **(E)**, **(F)** (*R,S*)-ketamine (30 mg/kg) and (2*R*,6*R*)-HNK (0.075 mg/kg) decreased immobility time on day 2 of the FST. **(G)** (2*S*,6*S*)-HNK did not alter immobility in the FST. (n = 8-15 male mice per group). Error bars represent ± SEM. * p < 0.05. ** p < 0.01. *** p < 0.0001. Sal, saline; K, (*R,S*)-ketamine; HNK, hydroxynorketamine; CFC, contextual fear conditioning; FST, forced swim test.

We have recently shown that a single injection of (*R,S*)-ketamine given 1 week prior to CFC can attenuate learned fear in male mice (25). Here, during CFC training, neither (*R,S*)-ketamine nor (2*R*,6*R*)-HNK at any dose altered freezing (**Figure S01A-S01B**). However, multiple doses of (2*S*,6*S*)-HNK (0.025, 0.1, 0.3, 10, and 30 mg/kg) reduced freezing during training (**Figure S01C**). Consistent with our previous results (*R,S*)-ketamine (30 mg/kg) reduced freezing upon re-exposure to the aversive training context (**Figure 1B**). Although (2*R*,6*R*)-HNK did not affect fear behavior at any dose, (2*S*,6*S*)-HNK (0.025, 0.075, 0.1, 0.3, 10, and 30 mg/kg) significantly reduced freezing during re-exposure (**Figure 1C-1D**).

The FST is a behavioral assay that is broadly considered to have predictive validity when quantifying antidepressant efficacy (28). On day 1 of the FST, immobility time was comparable between saline and (*R,S*)-ketamine mice (30 mg/kg) (**Figure S01D**). Both (2*R*,6*R*)-HNK (10 mg/kg) and multiple doses of (2*S*,6*S*)-HNK (0.075, 0.1, 0.3, 10, and 30 mg/kg) reduced immobility time (**Figure S01E-S01F**). However, on day 2, mice administered (*R,S*)-ketamine (30 mg/kg) and (2*R*,6*R*)-HNK (0.075 mg/kg) exhibited reduced immobility times when compared to saline (**Figure 1E-1F**). (2*S*,6*S*)-HNK at any dose was not sufficient to alter immobility on day 2 of the FST (**Figure 1G**). In addition to replicating previously published work in male mice, our results indicate that (2*R*,6*R*)-HNK and (2*S*,6*S*)-HNK are effective at preventing divergent stress-induced behaviors in male mice; specifically, (2*S*,6*S*)-HNK attenuates fear, while (2*R*,6*R*)-HNK decreases depressive-like behavior.

### (*R,S*)-ketamine and (2*R*,6*R*)-HNK, but not (2*S*,6*S*)-HNK, are prophylactic against stress-induced depressive-like behavior in female 129S6/SvEv mice

We then sought to test whether (*R,S*)-ketamine, (2*R*,6*R*)-HNK, or (2*S*,6*S*)-HNK could be prophylactic in female mice. Female mice were administered a single injection of saline, (*R,S*)-ketamine, (2*R*,6*R*)-HNK, or (2*S*,6*S*)-HNK at varying doses 1 week before CFC (**Figure 2A**). Five days later, mice were placed back in the CFC context and then administered the FST, OF, and TI test.

**Figure 2.**
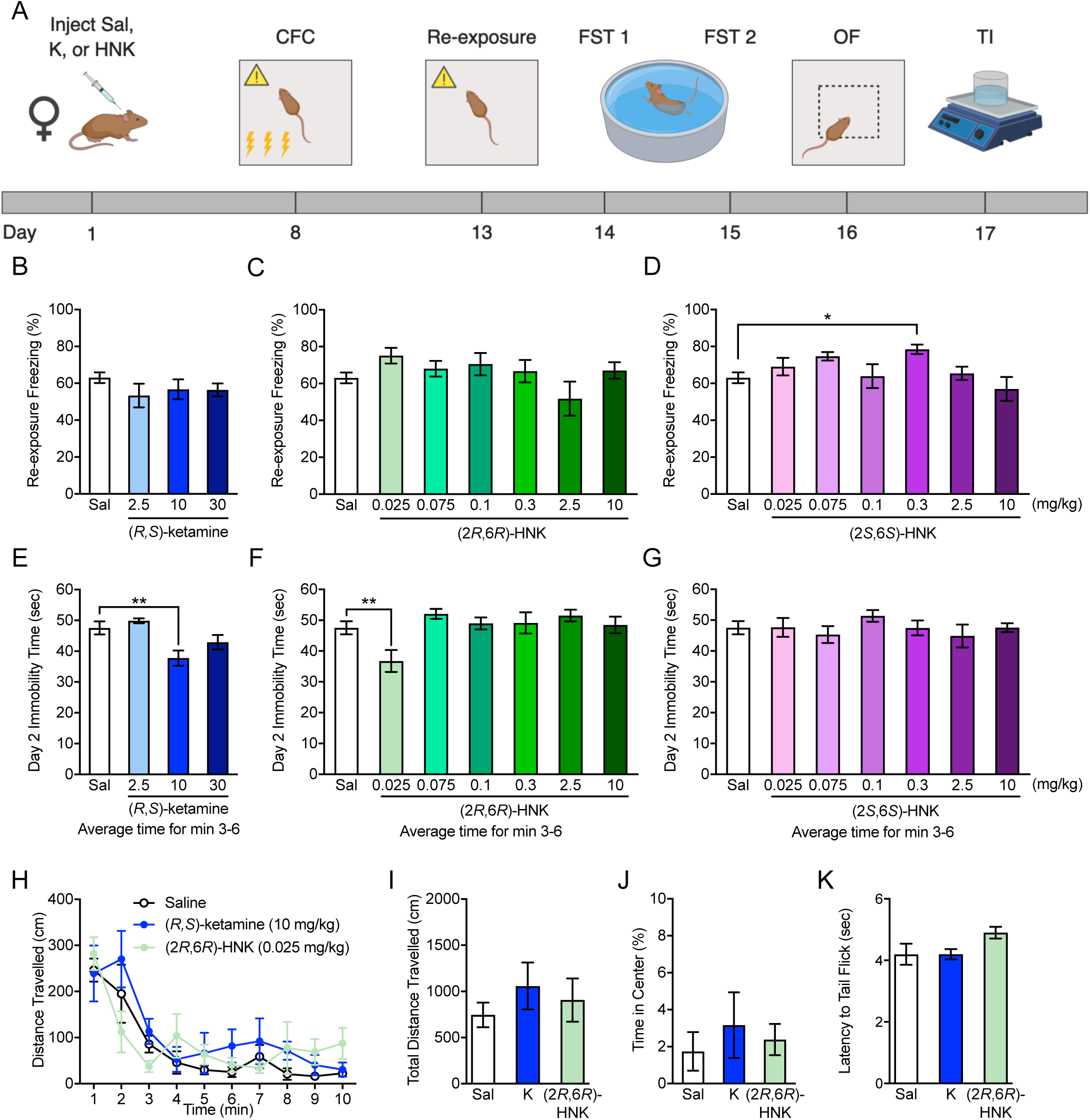
(*R,S*)-ketamine and (2*R*,6*R*)-HNK protect against stress-induced depressive-like behavior in female 129S6/SvEv mice. **(A)** Experimental design. **(B)**, **(C)**, **(D)** (*R,S*)-ketamine, (2*R*,6*R*)-HNK, and (2*S*,6*S*)-HNK did not alter freezing during re-exposure to the CFC context. **(E)**, **(F)** Administration of (*R,S*)-ketamine (10 mg/kg) and (2*R*,6*R*)-HNK (0.025 mg/kg) in female mice significantly reduced average immobility on day 2 of the FST. **(G)** (2*S*,6*S*)-HNK did not alter immobility in the FST. **(H)**, **(I)**, **(J)** (*R,S*)-ketamine (10 mg/kg) and (2*R*,6*R*)-HNK (0.025 mg/kg) did not alter distance travelled or time in the center of the OF when compared to saline. **(K)** Similarly, nociception was not significantly affected by (*R,S*)-ketamine or (2*R*,6*R*)-HNK. (n = 8-22 female mice per group). Error bars represent ± SEM. * p < 0.05. ** p < 0.01. Sal, Saline; K, Ketamine; HNK, hydroxynorketamine; CFC, contextual fear conditioning; FST, forced swim test; OF, open field; TI, tail immersion test.

Here, (*R,S*)-ketamine or (2*R*,6*R*)-HNK did not alter fear encoding (**Figure S02A-S02B**). However, (2*S*,6*S*)-HNK (0.3 mg/kg) increased freezing behavior during CFC training (**Figure S02C**). Upon re-exposure, (*R,S*)-ketamine or (2*R*,6*R*)-HNK did not alter fear expression (**Figure 2B-2C**). Again, (2*S*,6*S*)-HNK (0.3 mg/kg) increased freezing expression (**Figure 2D**). These data indicate that prophylactic (*R,S*)-ketamine and its metabolites do not attenuate learned fear in female mice as previously reported in male mice.

On day 1 of the FST, all groups exhibited comparable immobility time except mice administered (2*S*,6*S*)-HNK (0.3 mg/kg), which were more immobile when compared with saline mice (**S02D-S02F**). On day 2 of the FST, however, prophylactic (*R,S*)-ketamine (10 mg/kg) and (2*R*,6*R*)-HNK (0.025 mg/kg) significantly reduced immobility time when compared to saline (**Figure 2E-2G**). Immobility time in all groups administered (2*S*,6*S*)-HNK was not altered (**Figure 2G**).

Female mice were then assayed in the OF and TI test to determine if prophylaxis resulted in anxiolytic or nociceptive effects, respectively. There were no significant changes in total distance traveled in the OF, time spent in the center of the OF, or in tail withdrawal latency in the TI test across all groups (**Figure 2H-2K**). Therefore, the decreased immobility time in the FST is not confounded by nonspecific effects of either (*R,S*)-ketamine or (2*R*,6*R*)-HNK on locomotion or nociception. Overall, these data indicate that female mice require less than 1/3 of the dose needed by male mice to elicit the protective effects of both (*R,S*)-ketamine and (2*R*,6*R*)-HNK.

Previous data indicate that male and female rodents exhibit distinct pharmacokinetic profiles following ketamine administration (19). To examine how (*R,S*)-ketamine is metabolized in female 129S6/SvEv mice, we used liquid-chromatography mass spectrometry (LC-MS) to examine the levels of (*R,S*)-ketamine metabolites acutely after administration. Female mice were administered a single injection of saline, (*R,S*)-ketamine (10 mg/kg), (2*R*,6*R*)-HNK (0.025 mg/kg), or (2*S*,6*S*)-HNK (0.025 mg/kg) (**Figure S03A**). Ten minutes later, mice were sacrificed, and the brain and plasma were harvested. Following (*R,S*)-ketamine administration, a comparable level of (2*R*,6*R*)-HNK and (2*S*,6*S*)-HNK was found in both the brain and plasma (**Figure S03B-S03C**). However, in mice administered saline, (2*R*,6*R*)-HNK, or (2*S*,6*S*)-HNK, there were no detectable levels of either compound in the brain or plasma.

Next, we determined the effects of (*R,S*)-ketamine or (2*R*,6*R*)-HNK in non-stressed female mice. Mice were injected with saline, (*R,S*)-ketamine, or (2*R*,6*R*)-HNK 1 week before context exposure (**Figure S04A**). (*R,S*)-ketamine and (2*R*,6*R*)-HNK did not alter freezing behavior (**Figure S04B-S04D**) or depressive-like behavior (**Figure S04E-S04G**), indicating that the prophylactic effects of these compounds are specific to stress-induced behaviors.

### (*R,S*)-ketamine and (2*R*,6*R*)-HNK prevent LH-induced depressive-like behavior in female 129S6/SvEv mice

We have previously shown that prophylactic (*R,S*)-ketamine attenuates helplessness in male mice (24). To validate these findings in females, female mice were administered saline, (*R,S*)-ketamine, or (2*R*,6*R*)-HNK 1 week prior to LH training (**Figure 3A**). Two weeks later, mice were tested for escape latency and for depressive- and anxiety-like behaviors in the FST and elevated plus maze (EPM).

**Figure 3.**
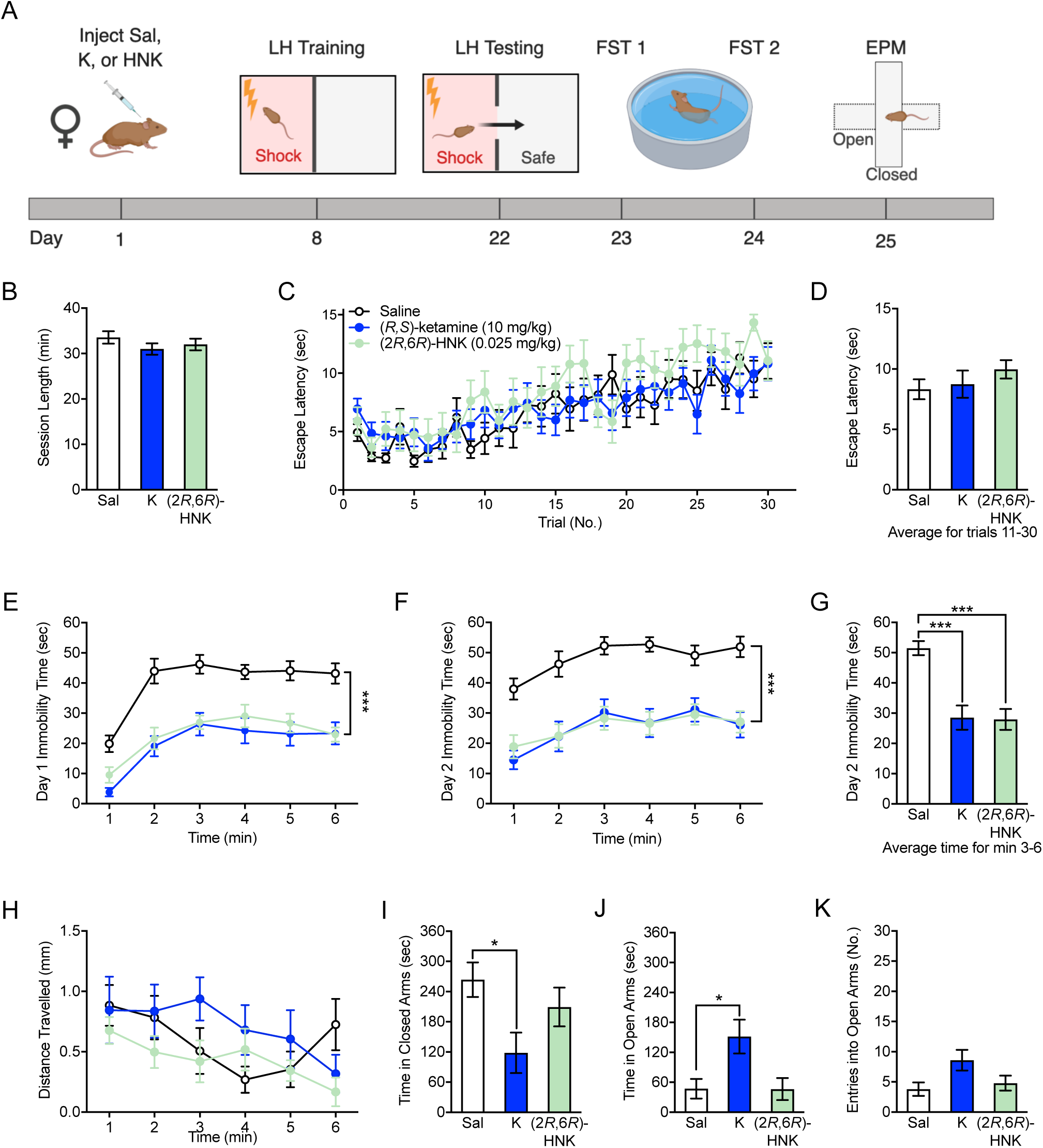
(*R,S*)-ketamine and (2*R*,6*R*)-HNK are prophylactic against LH stress in female 129S6/SvEv mice. **(A)** Experimental design. **(B)**, **(C),** (**D**) (*R,S*)-ketamine and (2*R*,6*R*)-HNK did not alter session length or escape latency when compared to saline in the LH assay. **(E), (F)**, **(G)** (*R,S*)-ketamine and (2*R*,6*R*)-HNK significantly reduced immobility on both days of the FST when compared to saline. **(H)** Mice in all groups travelled a comparable distance in the EPM. **(I)** (*R,S*)-ketamine mice spent significantly less time in the closed arms of the EPM when compared to saline mice. **(J)** Time spent in the open arms of the EPM was significantly increased in mice administered (*R,S*)-ketamine when compared with saline mice. **(K)** Entries into the open arms of the EPM was comparable in all groups. (n = 5-10 female mice per group). Error bars represent ± SEM. * p < 0.05. *** p < 0.0001. Sal, Saline; K, ketamine; HNK, hydroxynorketamine; LH, learned helplessness; FST, forced swim test; EPM, elevated plus maze.

There was no significant drug effect on session length or escape latency (**Figure 3B-3D**). However, in the FST, (*R,S*)-ketamine and (2*R*,6*R*)-HNK significantly decreased immobility time when compared to saline (**Figure 3E-4G**). In the EPM, mice in all groups travelled a comparable distance (**Figure 3H**). Mice administered (*R,S*)-ketamine, but not (2*R*,6*R*)-HNK, spent significantly less time in the closed arms and significantly more time in the open arms of the EPM when compared to saline mice (**Figures 3I-3J, S05A-S05B**). Entries into the open arms and time spent in the center of the EPM were comparable across all groups (**Figures 3K, S05C**). These data indicate that unlike in males, (*R,S*)-ketamine and (2*R*,6*R*)-HNK does not alter helpless behavior, but decreases stress-induced depressive-like behavior and may be anxiolytic in females exposed to LH stress.

### (*R,S*)-ketamine and (2*R*,6*R*)-HNK are prophylactic against CIS in female 129S6/SvEv mice

As previously tested in male mice (18), we validated our prophylactic findings in a model of chronic stress. Mice were administered saline, (*R,S*)-ketamine, or (2*R*,6*R*)-HNK (**Figure 4A**) 1 week before a 10-day CIS protocol. Subsequently, mice were tested in the FST and administered CFC. On day 1 of the FST, there was no difference in immobility between saline and (*R,S*)-ketamine mice **(Figure 4B**). (2*R*,6*R*)-HNK mice exhibited significantly increased immobility when compared with saline mice. However, on day 2 of the FST, both prophylactic drugs significantly lowered immobility when compared to saline (**Figure 4C-4D**). In both CFC training and re-exposure sessions, freezing behavior was comparable across all groups (**Figure 4E-4G**). These data indicate that prophylactic (*R,S*)-ketamine and (2*R*,6*R*)-HNK are sufficient to protect against the onset of depressive-like behavior following chronic stress.

**Figure 4.**
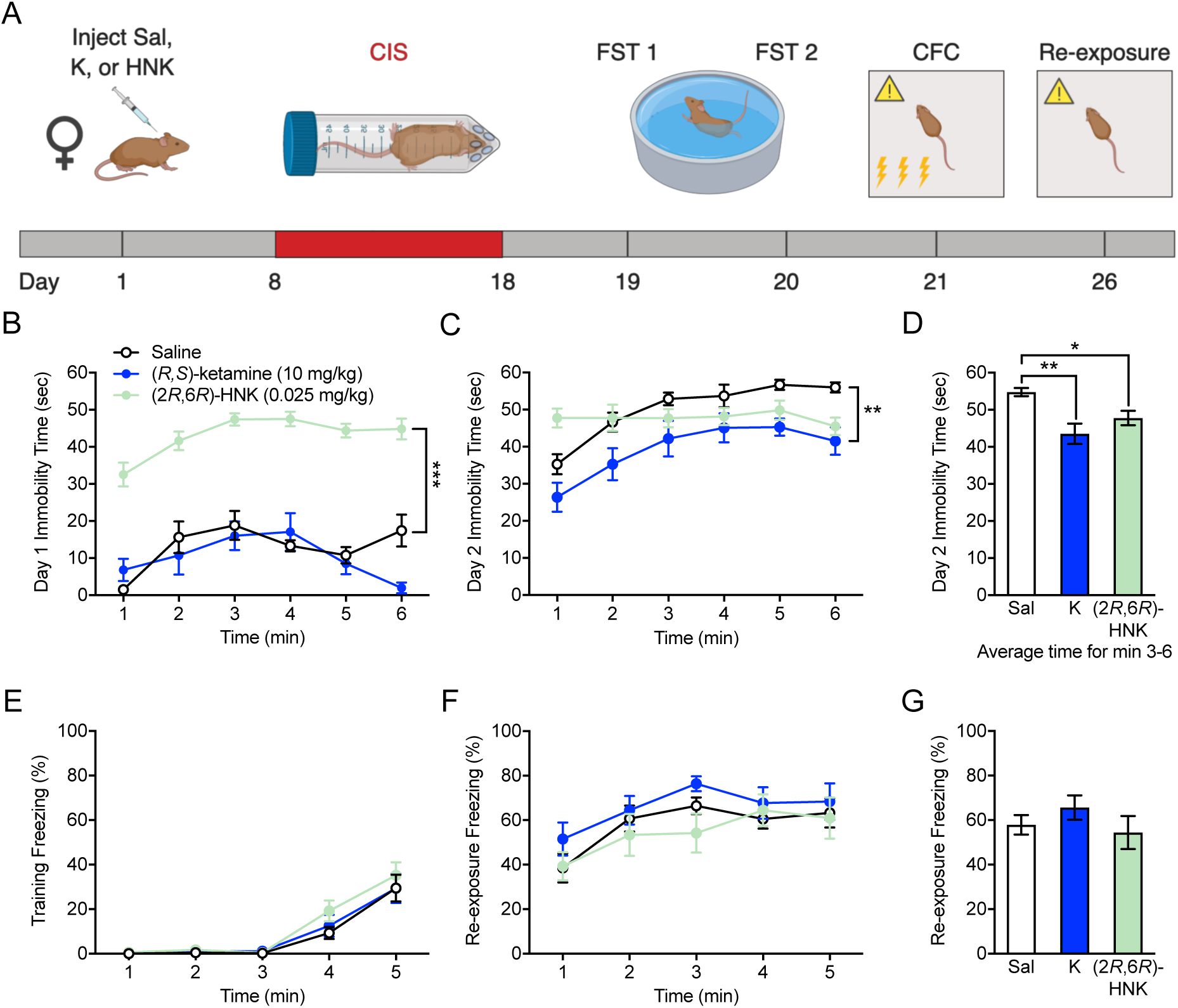
(*R,S*)-ketamine and (2*R*,6*R*)-HNK are prophylactic against CIS in female 129S6/SvEv mice. **(A)** Experimental design. **(B)** On day 1 of the FST, (2*R*,6*R*)-HNK mice were significantly more immobile when compared to saline mice. **(C)**, **(D)** On day 2 of the FST, (*R,S*)-ketamine and (2*R*,6*R*)-HNK reduced immobility when compared to saline. **(E)**, **(F)**, **(G)** Freezing during CFC training and re-exposure was comparable across all drug groups. (n = 9-10 mice per group). Error bars represent ± SEM. * p < 0.05. ** p < 0.01. *** p < 0.0001. Sal, Saline; K, ketamine; HNK, hydroxynorketamine; CIS, chronic immobilization stress; FST, forced swim test; CFC, contextual fear conditioning.

### (2*R*,6*R*)-HNK, but not (*R,S*)-ketamine, is prophylactic over a shorter time interval in female 129S6/SvEv mice

Previously, we determined that (*R,S*)-ketamine is prophylactic when administered one week, but not one day or one month, before exposure to stress in male mice (29). To determine the optimal time window of administration in female mice, we sought to determine if either compound could act prophylactically when given over a smaller time interval before stress (**Figure S06A**). We previously reported that (*R,S*)-ketamine attenuates learned fear when administered 1 week, but not 1 day or 1 month before stress in male mice (25). Here, saline, (*R,S*)-ketamine (2.5 mg/kg, 10 mg/kg, and 30 mg/kg), or (2*R*,6*R*)-HNK (0.025 mg/kg) was administered 3 days, instead of 1 week, before CFC in female mice.

During CFC training, (*R,S*)-ketamine (10 mg/kg) significantly increased freezing compared to saline (**Figure S06B**). During context re-exposure, all groups froze at comparable levels (**Figure S06C-S06D**). On both days of the FST, (2*R*,6*R*)-HNK mice reduced immobility time, indicating that (2*R*,6*R*)-HNK can be effective as a prophylactic at a shorter time interval (**Figure S06F-S06G**).

To test if (2*R*,6*R*)-HNK could be effective at an even smaller time interval, (2*R*,6*R*)-HNK was administered 24 hours before stress. However, (2*R*,6*R*)-HNK was not effective at attenuating learned fear or decreasing depressive-like behavior, indicating that both (*R,S*)-ketamine and (2*R*,6*R*)-HNK are only efficacious within a specific time window before stress (**Figure S07**).

### (*R,S*)-ketamine or (2*R*,6*R*)-HNK are not antidepressant in female 129S6/SvEv mice

Previous results have demonstrated that (*R,S*)-ketamine and (2*R*,6*R*)-HNK are rapid-acting antidepressants and effective in both sexes (14, 15, 19). However, this effect is strain- and stress-specific, as has been previously shown by our lab and others (30, 31). Here, we investigated if the prophylactically effective doses of (*R,S*)-ketamine and (2*R*,6*R*)-HNK were efficacious as antidepressants in nonstressed female mice. Mice were administered saline, (*R,S*)-ketamine, and (2*R*,6*R*)-HNK 1 hour prior to administration of the FST. (**Supplemental Figure S08A**). On both days of the FST, immobility was comparable across all drug groups (**Figure S08B-S08D**). These results demonstrate that the aforementioned prophylactic doses of (*R,S*)-ketamine and (2*R*,6*R*)-HNK are ineffective as antidepressants in nonstressed female mice.

We then sought to determine if (*R,S*)-ketamine or (2*R*,6*R*)-HNK could be effective when administered after a stressor. Mice were administered CFC and 1 hour later injected with saline, (*R,S*)-ketamine, or (2*R*,6*R*)-HNK, followed by the FST and re-exposure to the CFC context (**Figure S09A**). (*R,S*)-ketamine and (2*R*,6*R*)-HNK, administered after stress exposure, did not reduce immobility levels when compared to saline (**Figure S09B-S09D**). Additionally, we once again observed that neither compound affected fear behavior (**Figure S09E-S09F**).

### Ovarian hormones are necessary for the prophylactic efficacy of (*R,S*)-ketamine and (2*R*,6*R*)-HNK in female 129S6/SvEv mice

We hypothesized that the increased sensitivity to prophylactic (*R,S*)-ketamine exhibited by female mice was dependent on ovarian-derived hormones. To test the necessity of these hormones in facilitating (*R,S*)-ketamine’s effects, female mice were ovariectomized and allowed to recover for 10 days (**Figure 5A**). Mice were given a single injection of saline, (*R,S*)-ketamine, or (2*R*,6*R*)-HNK 1 week prior to administration of CFC.

**Figure 5.**
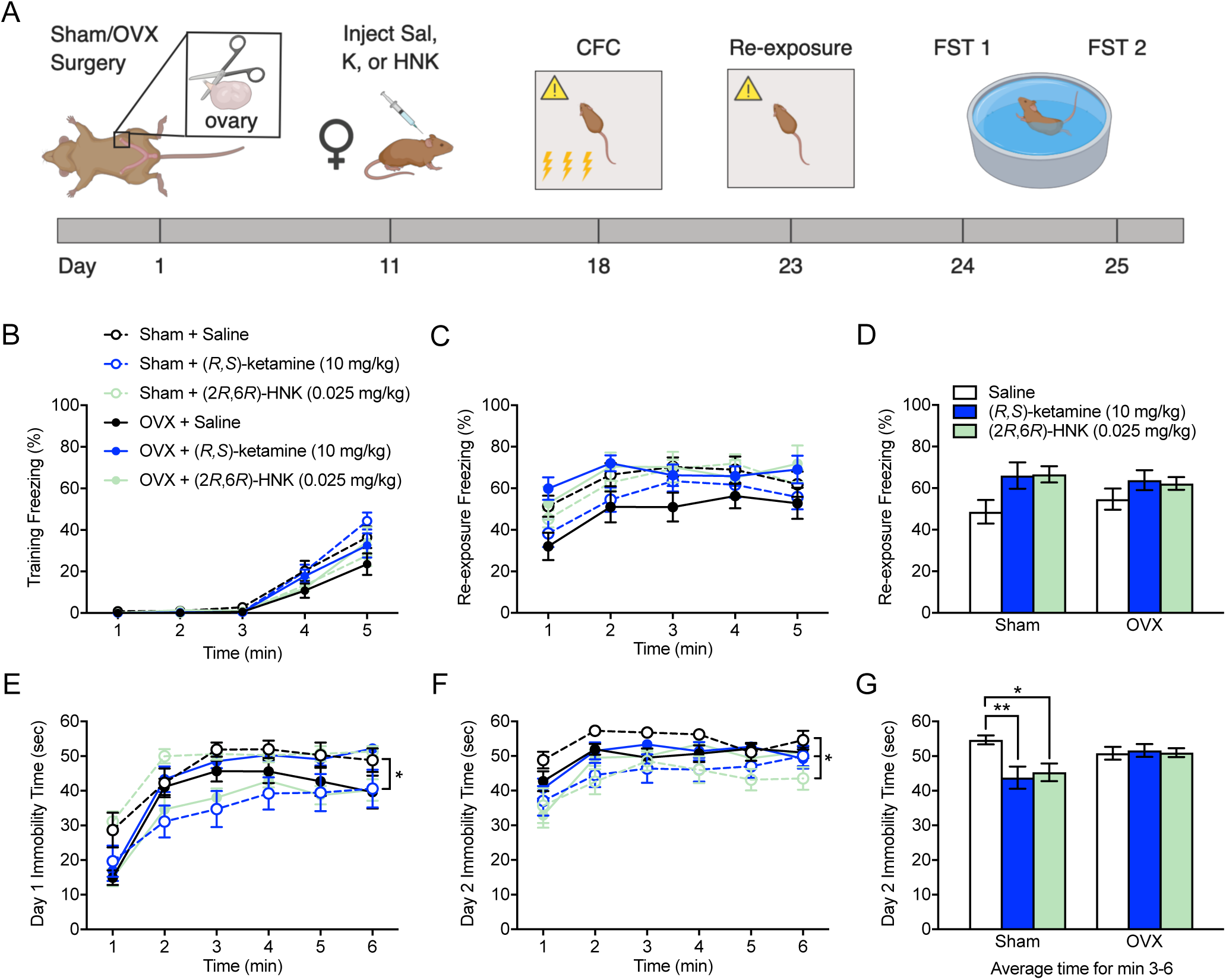
Ovarian hormones are necessary for the prophylactic efficacy of (*R,S*)-ketamine and (2*R*,6*R*)-HNK in female 129S6/SvEv mice. **(A)** Experimental design. **(B)**, **(C)**, **(D)** Neither surgery nor drug administration significantly altered freezing behavior during CFC training or re-exposure. **(E)** On day 1 of the FST, sham + (*R,S*)-ketamine mice had reduced immobility when compared to sham + saline mice. **(F)**, **(G)** On day 2 of the FST, within the sham group, (*R,S*)-ketamine and (2*R*,6*R*)-HNK reduced immobility when compared to saline. However, within the OVX group, there was no effect of drug on depressive-like behavior in the FST. (n = 13-15 mice per group). Error bars represent ± SEM. * p < 0.05. ** p < 0.01. OVX, ovariectomy; Sal, saline; K, (*R,S*)-ketamine; HNK, hydroxynorketamine; CFC, contextual fear conditioning; FST, forced swim test.

In both sham and OVX groups, mice administered saline, (*R,S*)-ketamine, or (2*R*,6*R*-HNK) froze at comparable levels during CFC training and re-exposure (**Figure 5B-5D**). On day 1 of the FST, in mice administered sham surgery, (*R,S*)-ketamine reduced immobility time when compared to saline **(Figure 5E**). On day 2 of the FST, in mice administered sham surgery, both (*R,S*)-ketamine and (2*R*,6*R*)-HNK decreased immobility time when compared with saline. However, in mice administered OVX surgery, (*R,S*)-ketamine and (2*R*,6*R*)-HNK were ineffective in decreasing immobility time when compared with saline (**Figure 6F-6G**). Therefore, these data suggest that ovarian-derived hormones are *necessary* for the prophylactic effects of (*R,S*)-ketamine and (2*R*,6*R*)-HNK in female mice.

**Figure 6.**
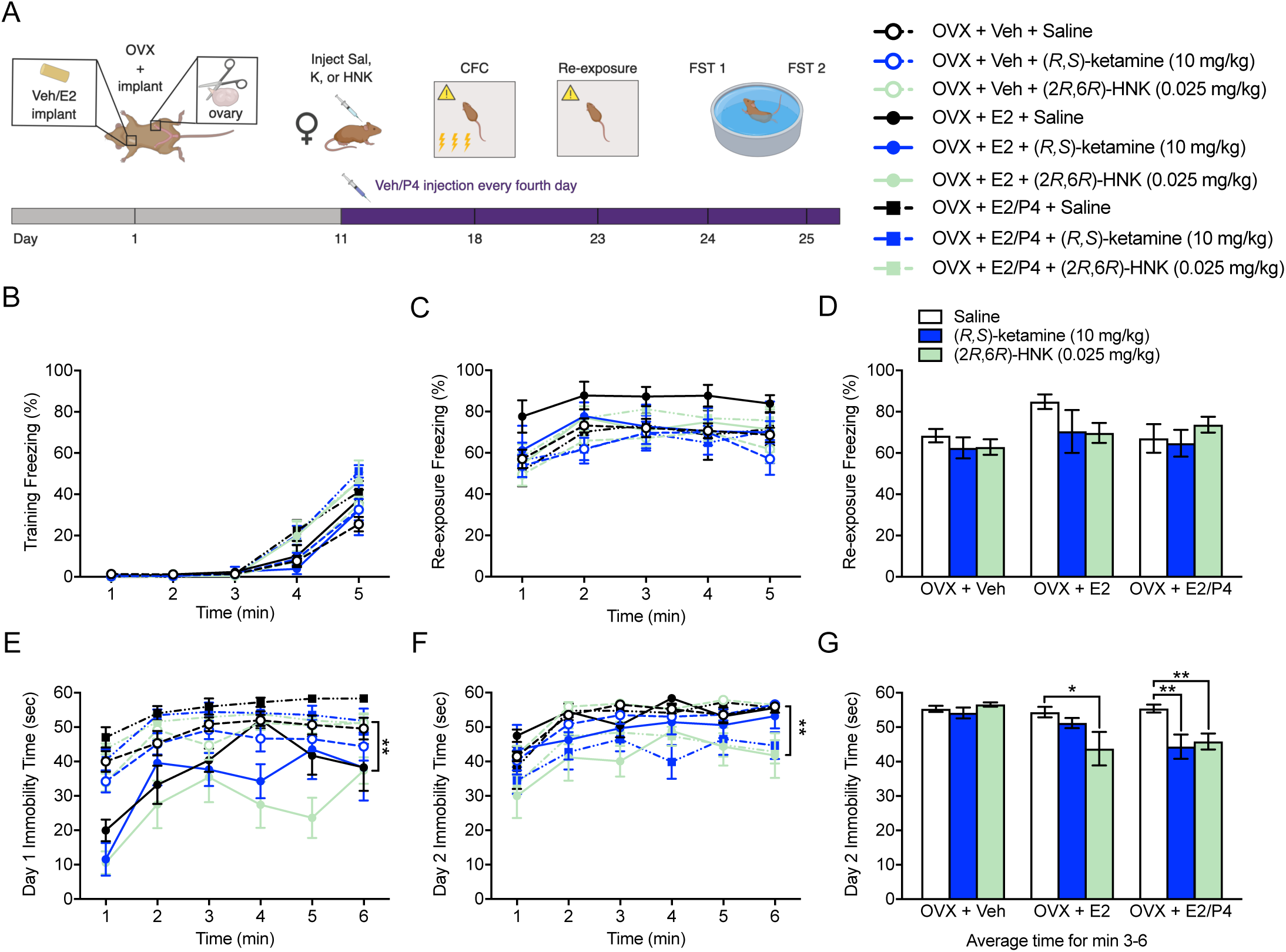
Ovarian hormones are sufficient to restore the prophylactic actions of (*R,S*)-ketamine and (2*R*,6*R*)-HNK in ovariectomized female 129S6/SvEv mice. **(A)** Experimental design and figure legend. **(B) (C)**, **(D)** Neither estrogen replacement, estrogen and cyclic progesterone replacement, nor prophylactic drug administration altered freezing during CFC training and re-exposure in OVX female mice. **(E)** On day 1 of the FST, E2 + (2*R*,6*R*)-HNK mice exhibited significantly reduced immobility when compared to Veh + (2*R*,6*R*)-HNK mice. **(F)**, **(G)** On day 2 of the FST, immobility was comparable between all drug groups in the OVX + Veh group. In the OVX + E2 group, (2*R*,6*R*)-HNK mice were significantly less immobile when compared to saline mice. In the OVX + E2/cyclic P4 group, both (*R,S*)-ketamine and (2*R*,6*R*)-HNK reduced immobility in the FST when compared to saline mice. (n = 5-18 mice per group). Error bars represent ± SEM. * p < 0.05. ** p < 0.01. Veh, vehicle; E2, estrogen; OVX, ovariectomy; P4, progesterone; Sal, saline; K, (*R,S*)-ketamine; HNK, hydroxynorketamine.

### Ovarian hormones are sufficient for the prophylactic efficacy of (*R,S*)-ketamine and (2*R*,6*R*)-HNK in female 129S6/SvEv mice

Subsequently, we aimed to investigate whether hormone replacement following OVX could restore the prophylactic effects of (*R,S*)-ketamine and (2*R*,6*R*)-HNK. Female mice were ovariectomized and separated into 1 of 3 groups: (1) following OVX, mice were implanted with a subcutaneous vehicle (Veh) implant and administered Veh injections every fourth day (OVX + Veh), (2) mice were implanted with a subcutaneous E2 implant and administered no injections (OVX + E2), or (3) mice were implanted with a subcutaneous E2 implant and administered P4 injections every fourth day (OVX + E2/P4) (**Figure 6A**). This hormone replacement protocol has previously been used to mimic the estrous cycle in rodents (32). Ten days after surgery, mice Veh/P4 injections were started and mice were given a single injection of saline, (*R,S*)-ketamine, or (2*R*,6*R*)-HNK and administered the same behavioral protocol as in **Figure 1A**.

During CFC training and re-exposure, all groups exhibited comparable levels of freezing (**Figure 6B-6D**). On day 1 of the FST, mice in the E2 + (2*R*,6*R*)-HNK group exhibited significantly reduced immobility when compared with mice in the Veh + (2*R*,6*R*)-HNK group (**Figure 6E**). There were no other differences in immobility time across all other groups. On day 2 of the FST, OVX + Veh control mice exhibited comparable immobility across all drug groups (**Figure 6F-6G**). In OVX + E2 mice, (2*R*,6*R*)-HNK significantly reduced immobility when compared with Sal. In OVX + E2/P4 mice, (*R,S*)-ketamine and (2*R*,6*R*)-HNK reduced immobility when compared to saline. E2 or E2/P4 replacement along did not significantly alter depressive-like behavior in the FST. These data show that E2 alone is sufficient to restore the prophylactic effects of (2*R*,6*R*)-HNK. However, E2/P4 replacement restores the protective effects of both (*R,S*)-ketamine and (2*R*,6*R*)-HNK.

## DISCUSSION

This series of experiments yielded three main findings: 1) (2*S*,6*S*)-HNK and (2*R*,6*R*)-HNK affect different stress-induced phenotypes in male mice; 2) (*R,S*)-ketamine and its metabolite (2*R*,6*R*)-HNK are prophylactic against stress-induced depressive-like behavior, but do not alter learned fear, in female mice, and 3) prophylactic efficacy of (*R,S*)-ketamine and (2*R*,6*R*)-HNK in female mice is modulated by ovarian hormones. In female mice, both (*R,S*)-ketamine and (2*R*,6*R*)-HNK prevented the onset of depressive-like behavior in a wide range of stress models when administered one week before stress exposure. Moreover, ovarian hormones were both necessary and sufficient for the protective properties of (*R,S*)-ketamine and (2*R*,6*R*)-HNK, suggesting that prophylactic compounds act on neural circuits mediated by gonadal hormones to enhance stress resilience in female mice.

To our knowledge, our study is the first to demonstrate that stereospecific metabolites of (*R,S*)-ketamine drive distinct protective effects in male mice. While (2*S*,6*S*)-HNK attenuated learned fear in male mice, (2*R*,6*R*)-HNK prevented stress-induced depressive-like behavior in both male and female mice. A number of previous studies have shown differing effects of stereospecific (*R,S*)-ketamine and (*R,S*)-ketamine metabolites on depressive-like behavior (19, 21-23, 33-35). (*S*)-ketamine and its metabolites possess a significantly greater affinity for the *N*-methyl-D-aspartate receptor (NMDAR) than their (*R*)-ketamine enantiomers (18). On the other hand, (2*R*,6*R*)-HNK is hypothesized to act preferentially on α-amino-3-hydroxy-5-methyl-4-isoxazolepropionic acid receptors (AMPARs), although some evidence suggests that this action may be dose-specific and may not have a direct effect on depressive-like behavior (19, 36-38). Our data suggest that the NMDAR may be a more critical target for modulating fear behavior in males, while alternative neural mechanisms may play a larger role in regulating stress-induced depressive-like behavior in both sexes.

Intriguingly, our data indicate that extremely low concentrations of (2*R*,6*R*)-HNK (0.025 mg/kg in female mice and 0.075 mg/kg in male mice) can still have a significant impact on behavior despite resulting in non-detectable levels in the brain and plasma following administration. Due to technical limitations, our LC-MS method may not be sensitive enough to detect brain or plasma concentrations of (2*R*,6*R*)-HNK within the picogram range, and future studies may have more success using accelerated mass spectrometry (39, 40). Although, to our knowledge, these microdoses of (*R,S*)-ketamine and its metabolites have not previously been studied, there has been a recent resurgence in examining the potential benefit of microdosing, in which small quantities of psychedelic substances are repeatedly ingested (39-47). This unique method of administration is reported to exert a myriad range of benefits, including improved mood, heightened creativity, and increased focus, without inducing nonspecific psychotropic or addictive side effects (43, 46, 47). Moreover, evidence suggests that studying microdosing at the preclinical stage may lead to more reliable predictions of the pharmacokinetics in human populations (41, 42, 44, 45). Further study is necessary to determine whether repeated small doses of (2*R*,6*R*)-HNK can extend the same protection against stress-induced depressive-like behavior as a single administration of the drug.

Historically, females have not been used to study pharmacological therapies for psychiatric disorders due to concerns over behavioral variability. However, previous studies indicate that male and female subjects respond differently to (*R,S*)-ketamine (14, 15). Although specific reasons for this difference are unknown, sex differences in pharmacokinetics or pharmacodynamics, such as differences in free drug concentrations or in the binding efficiency of certain brain receptors, could lead to greater transduction of (*R,S*)-ketamine’s effects in females (48-50). Another potential factor is the estrous cycle. Because our pharmacological and behavioral manipulations occurred over a long period of time, we did not track the estrous cycle in our study in order to minimize handling stress. However, ovarian hormones can significantly impact neuronal morphology, rodent behavior, and (*R,S*)-ketamine efficacy (51-53). Our results support the hypothesis that ovarian hormones induce changes at the neuronal level to causally mediate (*R,S*)-ketamine and (2*R*,6*R*)-HNK’s effects, whether antidepressant or prophylactic, and thus, impact behavior in females. Therefore, studying the interaction between ovarian-derived hormones and (*R,S*)-ketamine and/or its metabolites may facilitate the discovery of underlying mechanisms by which prophylactics enhance stress resilience.

In addition to differences in drug dosage, we observed dissimilarities in prophylactic (*R,S*)-ketamine’s effect on fear behavior between the sexes. Similar to our results, a recent study reported that prophylactic (*R,S*)-ketamine administered to female rats before inescapable shock stress prevented reductions in social exploration but did not report changes in LH behavior (24, 27). Indeed, across species, a number of studies have demonstrated divergent symptoms of mood disorders between sexes. While women diagnosed with MDD are more likely to experience a comorbid anxiety disorder and greater suicidal ideation, men are at greater risk of comorbid substance abuse (54). Moreover, in rodent models of depression, females do not acquire a LH phenotype, do not express anhedonia as strongly as males, and are more susceptible to swimming stress (55, 56). Thus, males and females likely process stress using separate neural strategies that result in distinct behavioral responses. Consequently, many paradigms developed to model pathological behavior in male animals may be inappropriate for use in female animals.

Here, we did not find that (*R,S*)-ketamine and (2*R*,6*R*)-HNK were antidepressant in female mice. However, we do not believe these results contradict previous findings (24-25,42). Instead, differences in mouse strain and drug dosing likely contributed to the discrepancies that we have demonstrated. Although we utilized 129S6/SvEvTac mice, ours and other previous studies have shown efficacy in BALB/cJ or C57BL/6J mice (19, 20, 33). Moreover, while antidepressant (2*R*,6*R*)-HNK is typically administered at 10 mg/kg doses, here, we administered this drug at 0.025 mg/kg. Thus, (*R,S*)-ketamine and (2*R*,6*R*)-HNK could be efficacious as antidepressants in females of different mouse strains or when administered at higher doses.

For future translation into human populations, the mechanisms of prophylactic (*R,S*)-ketamine and (2R,6*R*)-HNK must be further elucidated. In male mice, our lab has shown that prophylactic (*R,S*)-ketamine acts on the transcription factor ΔFosB to alter neural ensembles in the ventral hippocampus and can influence the balance of excitatory and inhibitory neurotransmitters in the brain after exposure to stress (57, 58). Other studies indicate that (*R,S*)-ketamine may sensitize prelimbic cortical inhibitory neurons projecting to the dorsal raphe nucleus (DRN) in order to exert its antidepressant effects in male and female rats (26, 27). Further study examining these and other resilience-enhancing mechanisms will be crucial for the development of next-generation prophylactic agents.

In summary, these experiments demonstrate that (*R,S*)-ketamine and (2*R*,6*R*)-HNK are effective prophylactics against a variety of stressors in females and are causally mediated by ovarian-derived hormones. Ultimately, this study offers insight into the prevention of stress-induced depressive-like behavior in a particularly susceptible population. Moreover, examining the mechanisms by which prophylactic compounds enhance stress resilience will elucidate the underlying neuropathology and sex differences driving MDD and contribute to advancements in targeted therapies for mood-based disorders.

## ACKNOWLEDGEMENTS

BKC was supported by Neurobiology & Behavior Research Training Grant T32 HD007430-19 and the Coulter Biomedical Accelerator (BioMedX) program. CTL was supported by an NIH DP5 OD017908. RAB was supported by the Coulter Biomedical Accelerator program. CAD was supported by an NIH DP5 OD017908, a New York Stem Cell Science NYSTEM C-021957, and a gift from For the Love of Travis, Inc. We thank M. Grunebaum, C. Anacker, S. Ramirez, B. McEwen, and members of the laboratory for insightful comments on this project and manuscript. Additionally, we thank Dr. Moshe Shalev and Lisa Moyano for instruction and assistance in performing ovariectomies.

## CONFLICT OF INTEREST

CTL, XX, SD, RFS, TBC, and DWL reported no biomedical financial interests or potential conflicts of interests. BKC, RAB, and CAD are named on provisional and non-provisional patent applications for the prophylactic use of (*R*,S)-ketamine and related compounds against stress-related psychiatric disorders.

