## Supplemental Information for "Ovarian hormones mediate the prophylactic efficacy of (*R,S*)-ketamine and (2*R*,6*R*)-hydroxynorketamine in female mice"

### SUPPLEMENTAL MATERIALS AND METHODS

#### Drugs

A single injection of saline (0.9% NaCl), (*R,S*)-ketamine (Fort Dodge Animal Health, Fort Dodge, IA), (2*R*,6*R*)-HNK (synthesized by the Organic Chemistry Collaborative Center (OCCC) at Columbia University), or (2*S*,6*S*)-HNK (synthesized by the OCCC) was administered once during the course of each experiment at approximately 8 weeks of age. All drugs were prepared in physiological saline and administered intraperitoneally (i.p.) in volumes of 0.1 cc per 10 mg body weight.

#### Contextual Fear Conditioning (CFC)

A 3-shock CFC paradigm was administered as previously described [1,2]. Fear conditioning was conducted in chambers obtained from Med Associates (St. Albans, VT), with internal dimensions of approximately 20 cm wide x 16 cm deep x 20.5 cm high. The chambers had metal walls on each side, clear plastic front and back walls and ceilings, and stainless steel bars on the floor. A house light (CM1820 bulb, 28v, 100mA) mounted directly above the chamber provided illumination. Each chamber was located inside a larger, insulated, plastic cabinet that provided protection from outside light and noise. Each cabinet contained a ventilation fan that was operated during the sessions. A paper towel dabbed with lemon solution was placed underneath the chamber floor. Mice were held outside the experimental room in their home cages prior to testing and transported to the conditioning apparatus individually in standard mouse cages. Chambers were cleaned with 70% EtOH between each set of mice. Training sessions were conducted using a 3-shock protocol. Mice were placed into the conditioning chamber and received shocks at 180 s, 240 s, and 300 s (2 s duration, 0.75 mA). Fifteen seconds after the last shock, mice were removed from the chamber. Overall, the training session lasted 317 s. During re-exposure, mice were placed in the conditioning chamber for 5 minutes and did not receive any shocks. All sessions were scored for freezing using FreezeFrame4.

**Learned Helplessness (LH)**

In this paradigm, mice are exposed to unpredictable and uncontrollable stress (shocks) and then develop coping deficits to deal with the inscape shocks. We modified a previously published paradigm [3,4]. Briefly, the procedure was conducted in a two-chamber shuttle box (model ENV 010MD; Med Associates, St. Albans, VT) located within a sound-attenuated cubicle. The grid floor was made of stainless steel and connected to a shock generator. The scrambled shock generator (model ENV 414S, Med Associates) created varying electrical potential differences between bars preventing an animal from avoiding shock.

*Inescapable shock (training)*: At approximately 8 weeks of age, mice were trained in the LH paradigm. For each shuttle box, 2 animals were administered the protocol at the same time; the central door was closed, with one animal in each side's chamber. After a 3 min habituation period, the shock deliveries began. The training protocol consisted of 70 shocks, each with a 3 s average duration, at 0.5 mA, and with an intertrial interval (ITI) of approximately 15 s.

*Shock escape (testing):* Mice were tested in the same shuttle box used in the inescapable shock training. The box consisted of two identical chambers (17 l x 20 w x 17 h), separated by an automated door that opened vertically. The shuttle box was equipped with 8 infrared beams (4 on each side) for detecting position and activity of the animal (Med Associates, St. Albans, VT). Each mouse was placed into the right chamber with the door raised and was allowed to freely explore both chambers for 3 minutes. Then the door then closed automatically.

At the beginning of each trial, the door was raised and 5 s later a foot shock (0.5 mA) was delivered. The subject's exit from the shocked side ended the trial. If the mouse did not exit after 15 s, the shock was turned off and the trial ended. The door was lowered at the end of the trial. A session consisted of 30 trials separated by a 30 s ITI. Escape latencies were computed as the time from shock onset to the end of trial. If the subject failed to make a transition the maximum 15 s was used for the escape latency score.

**Chronic Immobilization Stress (CIS)**

The CIS procedure induces a depressive-like phenotype in test subjects and has both predictive and face validity [5]. Here, we used an adapted protocol from Ramirez *et al*. [6]. Mice were restrained using Mouse DecapiCone disposable restrainers (Braintree Scientific, Braintree, MA). Restraint lasted 2 hours per day for 10 consecutive days.

**Forced Swim Test (FST)**

The FST is typically used in rodents to screen for potential human antidepressants [7,8]. In fact, many papers examining ketamine in mouse models only observe effects in the FST [9-11]. In the FST, time spent immobile, as opposed to swimming, is used as a measure of depressive behavior.

The FST was administered as previously described [12]. Briefly, mice were placed into clear plastic buckets 20 cm in diameter and 23 cm deep filled 2/3 of the way with 22°C water. Mice were videotaped from the side for 6 min and were exposed to the swim test on 2 consecutive days. Immobility time was scored by an experimenter blind to the experimental groups.

**Tail Immersion (TI) Test**

The TI test was administered as previously described [13]. Compared to other nociceptive tests, TI provides reliable results across and within subjects [13]. Prior to testing, mice were habituated to a restraint apparatus for 5 days, which consisted of a Falcon tube through which was drilled 10 air holes of 2 mm in diameter (Fisher Scientific, Pittsburgh, PA). During the test, 50 mL of water was heated to 52°C. Mice were immobilized in the tube with their tail hanging freely before dipping the last two-thirds of the tail into the hot water. Tail withdrawal latency was measured in seconds using a stopwatch. Mice were tested in 3 consecutive trials and the average across all 3 trials was used for analysis.

**Elevated Plus Maze (EPM)**

Testing was performed as previously described [14]. Briefly, the maze is a plus-cross-shaped apparatus consisting of four arms, two open and two enclosed by walls, linked by a central platform at a height of 50 cm from the floor. Mice were individually placed in the center of the maze facing an open arm and were allowed to explore the maze for 5 min. The time spent in and the number of entries into the open arms was used as an anxiety index. Videos were scored using ANY-maze behavior tracking software (Stoelting, Wood Dale, IL).

**Open Field (OF)**

The OF assay was administered as previously described [14]. Briefly, motor activity was quantified in four Plexiglas open field boxes 43×43 $cm^2$ (MED Associates, Georgia, VT). Two sets of 16 pulse-modulated infrared photobeams on opposite walls 2.5-cm apart recorded x–y ambulatory movements. Activity chambers were computer interfaced for data sampling at 100-ms resolution. The computer defined grid lines that dividing center and surround regions, with the center square consisting of four lines 11 cm from the wall.

**Plasma And Brain Analysis of Hydroxynorketamine (HNK) Enantiomers**

Mice were sacrificed 10 minutes following saline or drug injection via cervical dislocation. Immediately, mice were decapitated, and the brain was extracted from the skull and placed on dry ice for 30 seconds. Following freezing, the cerebellum was removed, and remaining brain tissue was sectioned into right and left hemispheres and placed into 1 mL Eppendorf tubes. Brain tissue was then immediately stored on dry ice.

For blood collection, trunk blood was collected immediately after decapitation using a 1000 µl pipette and placed into Eppendorf tubes coated with 0.5 mL of ethylenediaminetetraacetic acid (EDTA) (Invitrogen, Waltham, MA). Blood was then spun down for 10 minutes at 2000 x g and at 4ºC, and the supernatant (e.g. plasma) was collected for analysis.

Plasma and brain levels of (2*S*,6*S*)- and (2*R*,6*R*)-HNK enantiomers were quantified by a liquid-liquid extraction procedure, followed by a modified, previously described liquid chromatographic method with mass spectrometric detection [15]. The method was modified using a 3µ, 250 x 4.6mm Lux-Amylose-2 chiral column (Phenomenex®, Torrance, CA), with a mobile phase consisting of 64% 10 mM ammonium formate (pH=7.4), 32% acetonitrile, and 4% 2-propanol at a flow rate of 0.8ml/min. The mass spectrometer was set at the positive APCI mode for the detection of the molecular ions, which included the internal standard monoethylglycinexylidide (MEGX). An 8-point calibration curve in the range of 300 to 5 ng/mL of each enantiomer, in either plasma or brain homogenate, was included with each assay. The retention times for (2*S*,6*S*)-HNK and (2*R*,6*R*)-HNK and MEGX were 5.3, 6.6, and 5.7 min, respectively. Intra-assay variation (C.V.) based upon 8 replicates of each calibration concentration did not exceed 12%. Inter-assay variation (C.V.) based upon 3 concentrations of quality controls did not exceed 11% (n=8 consecutive days).

**Ovariectomy (OVX) Surgery**

Surgery was performed as previously described [16]. Mice were anesthetized with 1.5% isoflurane in an $O_2/N_2O$ (30%/70%) mixture and placed on a T/Pump heating pad

(Stryker, Kalamazoo, MI). A dorsal 3x3 cm area was shaved and disinfected with Betadine and alcohol before making a 2-cm midline incision. Ovarian and surrounding adipose tissue were bilaterally cut and cauterized. The fascia was closed with Coated VICRYL© Sutures (Johnson & Johnson, New Brunswick, NJ), and the skin was closed with 7 mm wound clips (Braintree Scientific, Braintree, MA). The closed incision was treated with a 5% topical lidocaine gel (ESBA Laboratories, Jupiter, FL). Prior to and for 3 days after surgery, mice were administered Carprofen (5 mg/kg) subcutaneously or perorally (MediGel CPF, Clear $H_2O$, Portland, ME) and allowed to recover for 10 days before behavioral testing.

**Hormone Replacement**

Silastic estrogen (E2) capsules were prepared and implanted as previously described [17]. Two cm lengths of silastic tubing (Dow Corning, Midland, MI) were cut and filled with a 2 mg/ml solution of 17β-estradiol (Sigma Aldrich, St. Louis, MO) dissolved in sesame oil (Sigma Aldrich, St. Louis, MO). As controls, vehicle implants were filled with sesame oil. Implants were sealed with silicone sealant (Dap Products, Baltimore, MD) and subcutaneously implanted. A 2x2 cm area along the dorsal aspect of the neck was shaved and disinfected with Betadine and alcohol before making a 2 mm incision in the skin. A small subcutaneous pocket was gently created, and the implant was placed vertically along the body. The incision was then closed with sutures (Johnson & Johnson, New Brunswick, NJ) and treated with 5% topical lidocaine gel. Prior to and for 3 days after surgery, mice were administered Carprofen (5 mg/kg) subcutaneously or perorally (MediGel CPF, Clear $H_2O$, Portland, ME) and allowed to recover for 10 days before behavioral testing. Cyclic progesterone administration began at the time of saline, (*R,S*)-ketamine, or (2*R*,6*R*)-HNK injection and continued for the remainder of the behavioral protocol. A 1.5 mg/ml progesterone solution of progesterone (Sigma Aldrich, St. Louis, MO) was prepared in sesame oil and injected subcutaneously every 4 days.

**Statistical Analysis**

All data were analyzed using StatView 5.0 (SAS Institute, Cary, NC) or Prism 7.0 (Graphpad Software, La Jolla, CA). Alpha was set to 0.05 for all analyses. Generally, the effect of Drug or Group was analyzed using an analysis of variance (ANOVA), using repeated measures where appropriate. Post-hoc Dunnett, Sidak, or Tukey tests were used where appropriate. All statistical tests and *p* values are listed in **Table S01**.

### SUPPLEMENTAL FIGURES AND LEGENDS

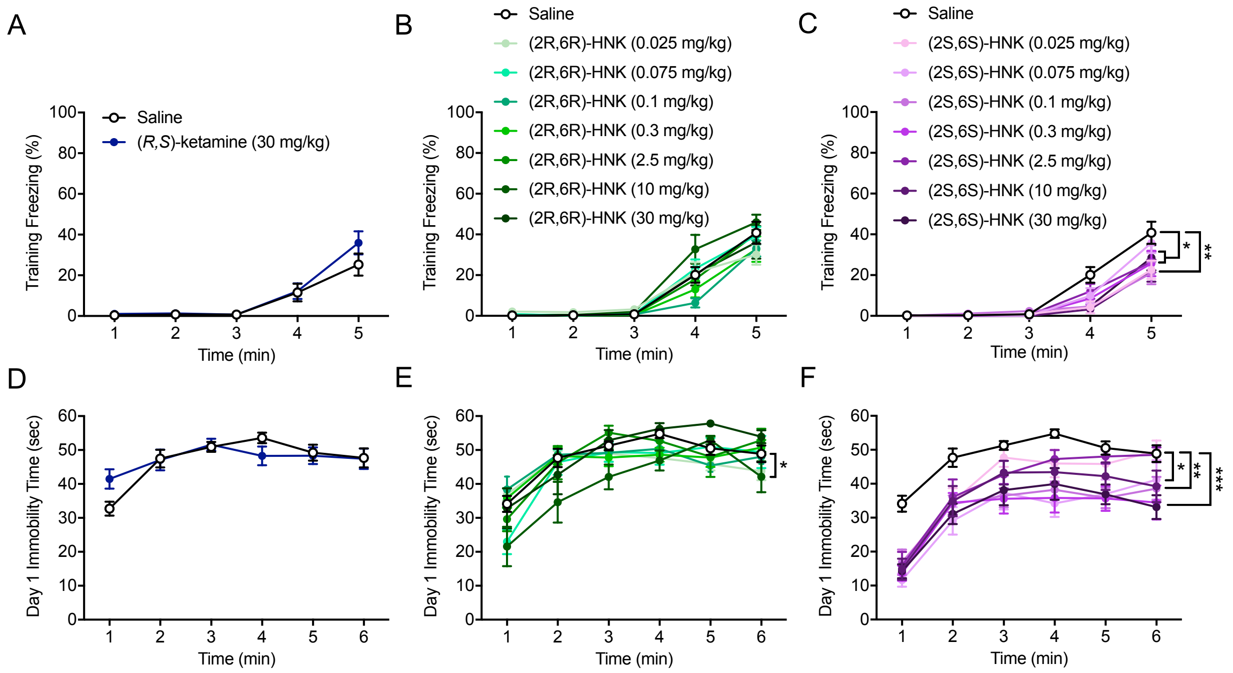

**Figure S01.** Prophylactic (*R,S*)-ketamine, (2*R*,6*R*)-HNK, and (2*S*,6*S*)-HNK differentially protect against stress in male mice. **(A)** (*R*,*S*)-ketamine did not alter freezing behavior during CFC training. **(B)** All doses of (2*R*,6*R*)-HNK did not alter freezing behavior during CFC training. **(C)** Numerous doses of (2*S*,6*S*)-HNK (0.025, 0.1, 0.3, 10, and 30 mg/kg) decreased freezing behavior during CFC training when compared with administration of saline. All other doses of (2*S*,6*S*)-HNK did not alter freezing behavior during CFC training. **(D)** (*R*,*S*)-ketamine did not impact immobility time during day 1 of the FST. **(E)** (2*R*,6*R*)-HNK (10 mg/kg) reduced immobility time during day 1 of the FST. All other doses of (2*R*,6*R*)-HNK did not impact immobility time during day 1 of the FST. **(I)** Administration of (2*S*,6*S*)-HNK (0.075, 0.1, 0.3, 10, and 30 mg/kg) decreased immobility time during day 1 of the FST when compared with administration of saline. All other doses of (2*S*,6*S*)-HNK do not alter immobility time during day 1 of the FST. (n = 8-15 male mice per group). Error bars represent ± SEM. * $p < 0.05$, ** $p < 0.01$. *** $p < 0.0001$. (2*R*,6*R*)-HNK, (2*R*,6*R*)-hydroxynorketamine.

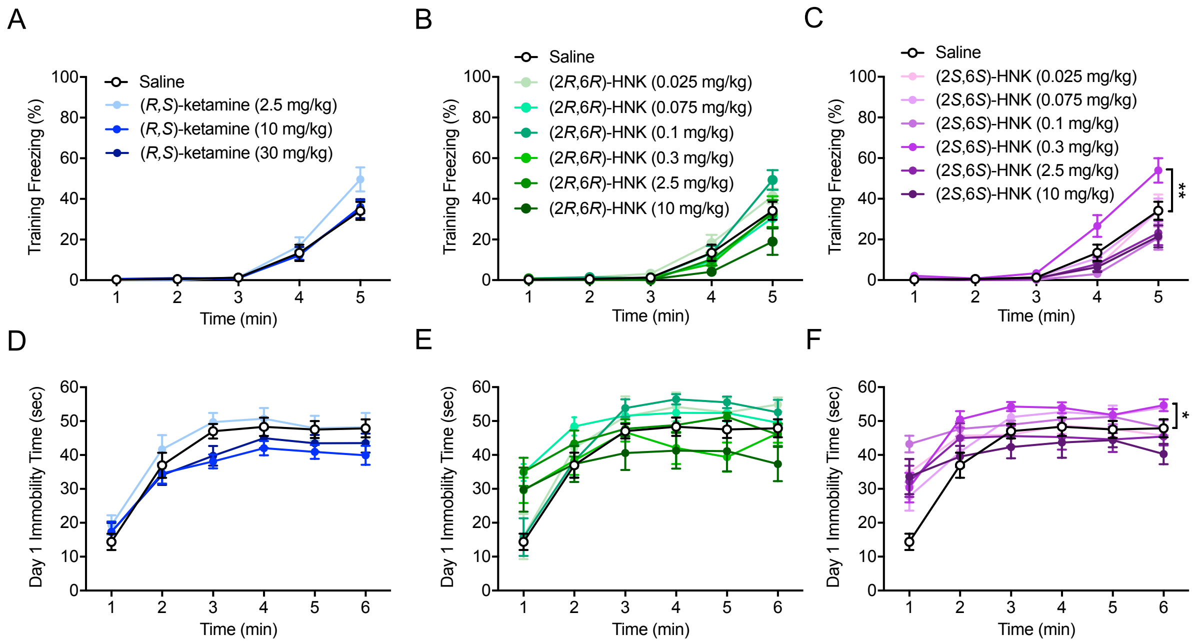

**Figure S02.** Prophylactic (*R,S*)-ketamine and (2*R*,6*R*)-HNK, but not (2*S*,6*S*)-HNK, decrease stress-induced depressive-like behavior in female mice. **(A)** All doses of (*R*,*S*)-ketamine did not alter freezing behavior during CFC training. **(B)** All doses of (2*R*,6*R*)-HNK did not alter freezing behavior during CFC training. **(C)** Administration of (2*S*,6*S*)-HNK (0.3 mg/kg) increased freezing behavior during CFC training when compared with administration of saline. All other doses of (2*S*,6*S*)-HNK did not alter freezing behavior during CFC training. **(D)** All doses of (*R*,*S*)-ketamine did not impact immobility time during day 1 of the FST. **(E)** All doses of (2*R*,6*R*)-HNK did not impact immobility time during day 1 of the FST. **(F)** Administration of (2*S*,6*S*)-HNK (0.3 mg/kg) increased immobility time during day 1 of the FST when compared with administration of saline. All other doses of (2*S*,6*S*)-HNK do not alter immobility time during day 1 of the FST. (n = 8-22 female mice per group). Error bars represent ± SEM. * $p < 0.05$, ** $p < 0.01$. (2*R*,6*R*)-HNK, (2*R*,6*R*)-hydroxynorketamine.

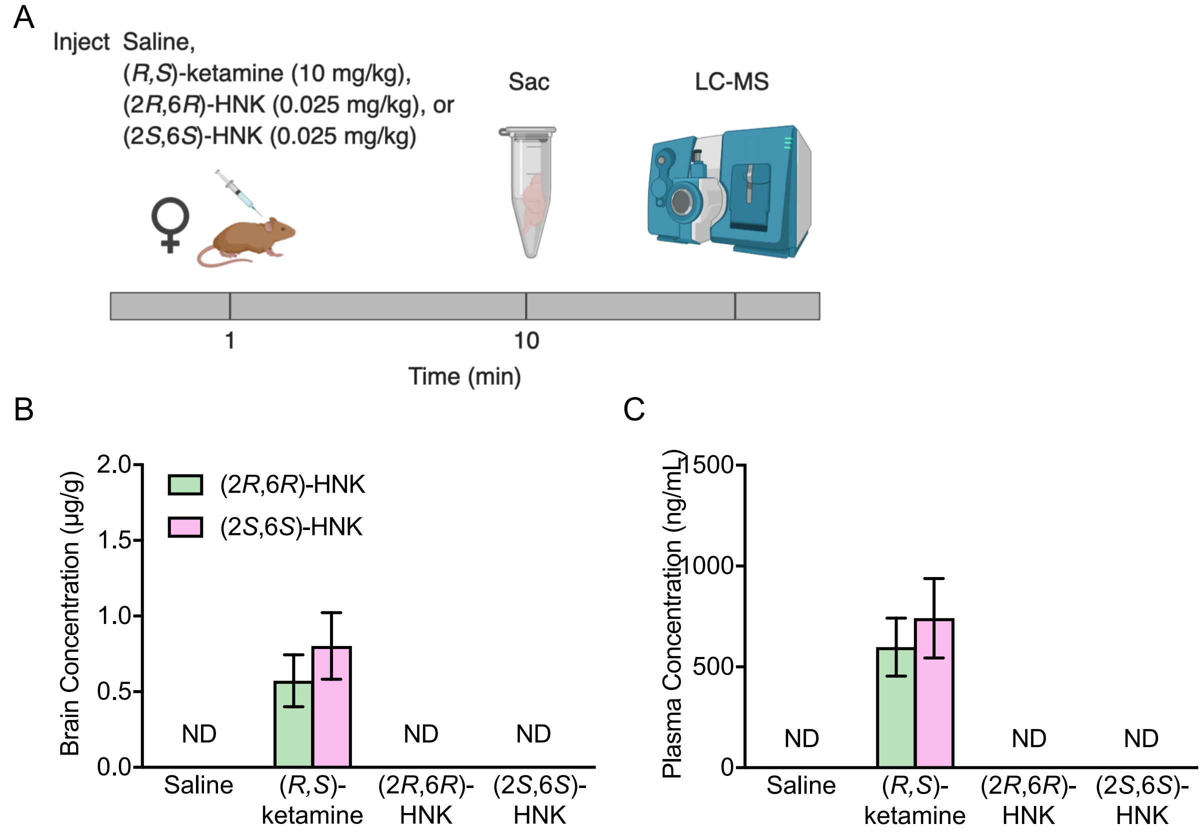

**Figure S03**. (*R*,*S*)-ketamine, but not (2*R*,6*R*)-HNK or (2*S*,6*S*)-HNK, results in detectable levels of (2*R*,6*R*)-HNK and (2*S*,6*S*)-HNK in the brain and plasma. (**A**) Experimental design. Female mice were injected with either saline, (*R*,*S*)-ketamine (10 mg/kg), (2*R*,6*R*)-HNK (0.025 mg/kg), or (2*S*,6*S*)-HNK (0.025 mg/kg). Ten minutes later mice were euthanized. Brain tissue and plasma were immediately harvested for LC-MS. (**B**) An (*R*,*S*)-ketamine injection resulted in detectable levels of both (2*R*,6*R*)-HNK and (2*S*,6*S*)-HNK in the brain. All other groups did not have detectable levels. (**C**) An (*R*,*S*)-ketamine injection resulted in detectable levels of both (2*R*,6*R*)-HNK and (2*S*,6*S*)-HNK in plasma. All other groups did not have detectable levels. (n = 5 female mice per group). Error bars represent ± SEM. (2*R*,6*R*)-HNK, (2*R*,6*R*)-hydroxynorketamine;

(2*S*,6*S*)-HNK, (2*S*,6*S*)-hydroxynorketamine; sac, sacrifice; LC-MS, liquid chromatography-mass spectrometry; ND, not detected.

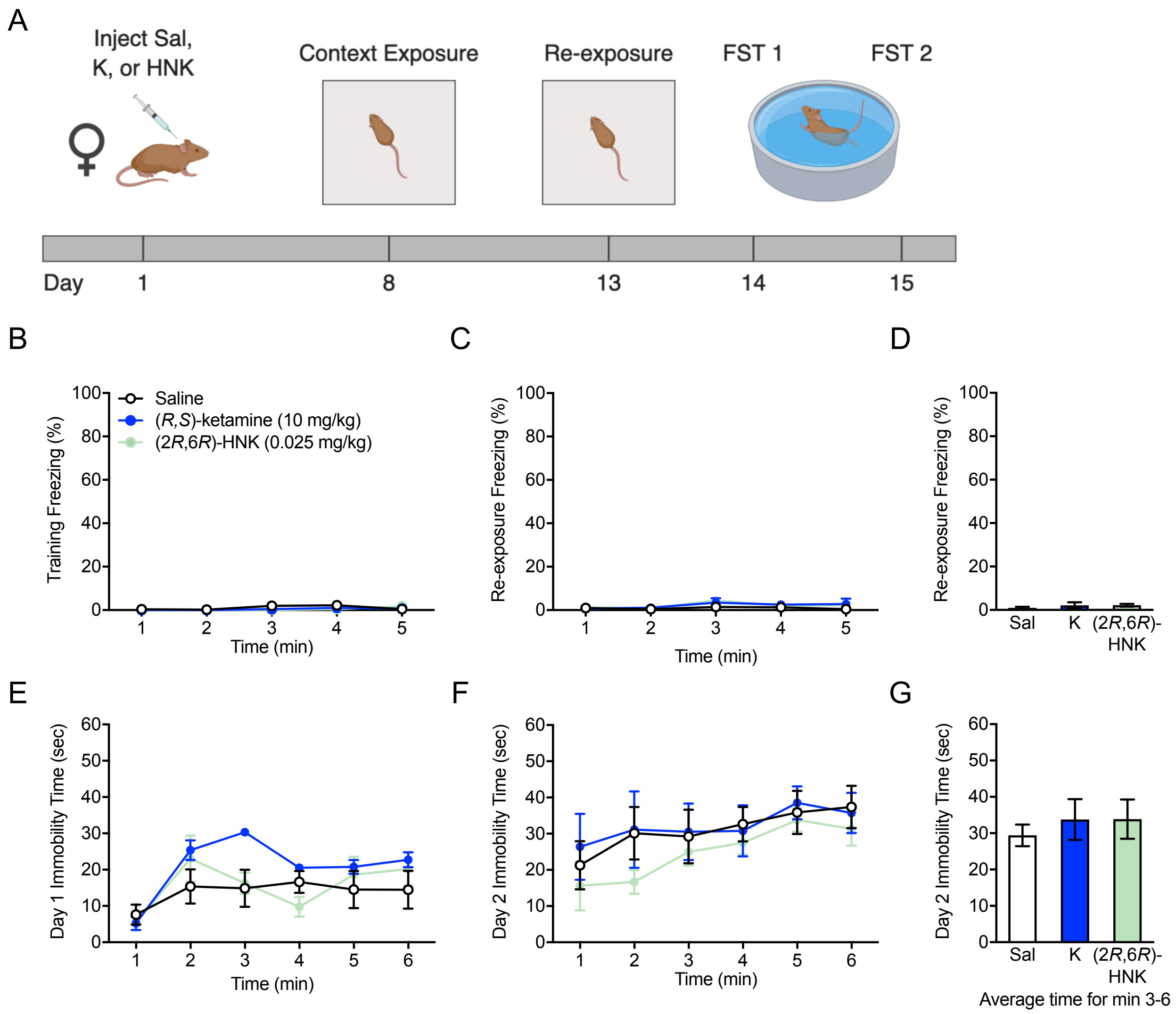

**Figure S04.** Administration of (*R*,*S*)-ketamine or (2*R*,6*R*)-HNK 1 week prior to context exposure does not alter depressive-like behavior in the FST in female mice. (**A**) Experimental design. Female mice were injected with saline, (*R,S*)-ketamine (10 mg/kg), or (2*R*,6*R*)-HNK one week prior to a no-shock context exposure. Five days later, mice were re-exposured to the same context. Mice were then tested in 2 days of the FST. (**B-D**) All groups of mice exhibited no freezing behavior during both context exposures. (**E-G**) All groups of mice exhibited comparable levels of immobility during both days of the FST. (n = 5 female mice per group). Error bars represent ± SEM. Sal, saline; K, (*R*,*S*)-ketamine; (2*R*,6*R*)-HNK, (2*R*,6*R*)-hydroxynorketamine; FST, forced swim test.

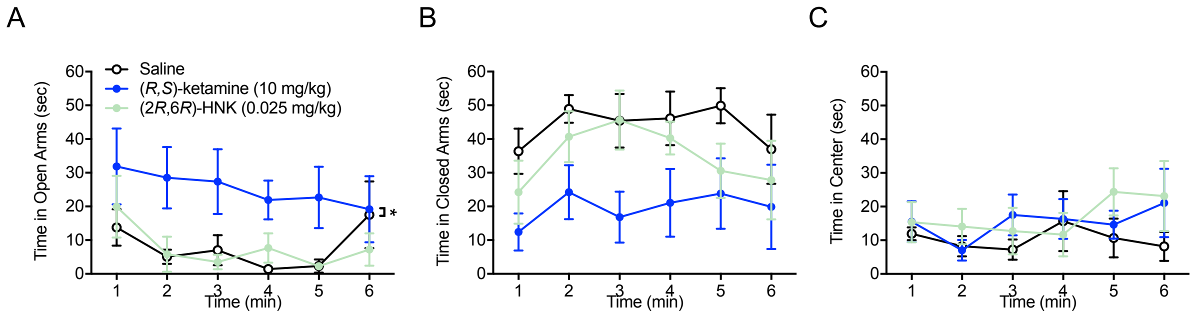

**Figure S05.** Administration of (*R*,*S*)-ketamine but not (2*R*,6*R*)-HNK 1 week prior to LH decreases stress-induced anxiety-like behavior in the EPM in female mice. (**A**) Mice administered (*R*,*S*)-ketamine (10 mg/kg) exhibited increased time in the open arms of the EPM relative to mice administered saline. (**B**) There was a trending, but not significant, effect of Drug on time spent in the closed arms of the EPM. (**C**) All groups of mice exhibited comparable time in the center of the EPM. (n = 5 female mice per group). Error bars represent ± SEM. (2*R*,6*R*)-HNK, (2*R*,6*R*)-hydroxynorketamine.

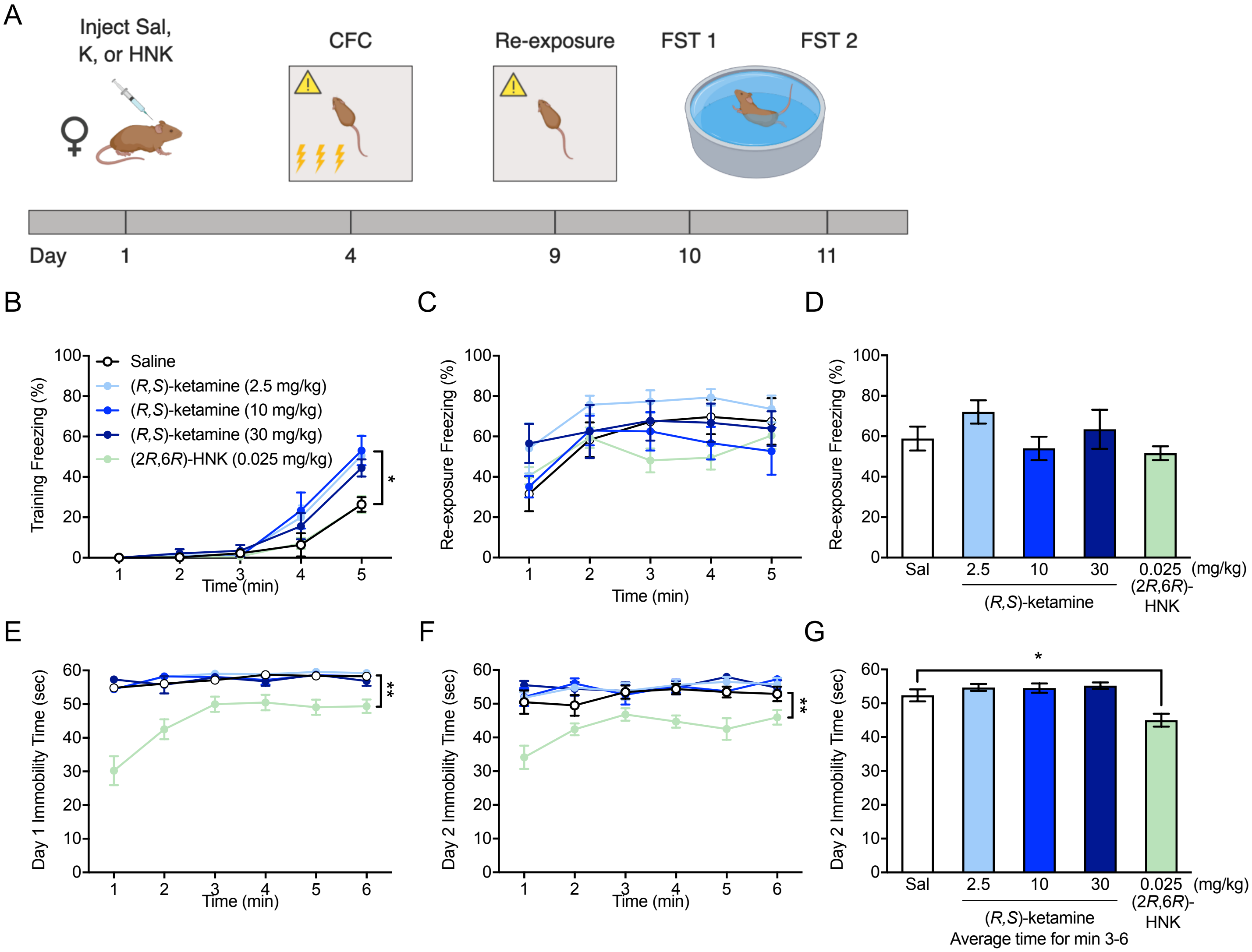

**Figure S06.** Administration of (2*R*,6*R*)-HNK, but not (*R*,*S*)-ketamine, 3 days prior to CFC decreases stress-induced depressive-like behavior in female mice. (**A**) Experimental design. Mice were administered a single injection of saline, (*R,S*)-ketamine (2.5, 10, or 30 mg/kg), or (2*R*,6*R*)-HNK (0.025 mg/kg) 3 days prior to 3-shock CFC. Five days later, mice were re-exposure to the training context. Mice were then administered 2 days of the FST. (**B**) Mice administered (*R*,*S*)-ketamine (10 mg/kg) exhibited increased freezing relative to mice administered saline during CFC training. All other groups of mice exhibited comparable freezing during CFC training. (**C-D**) All groups of mice froze comparably during CFC re-expsoure. (**E**) Administration of (2*R*,6*R*)-HNK, but not (*R*,*S*)-ketamine, 3 days before CFC decreased stress-induced depressive-like behavior on day 1 of the FST in female mice. (**F-G**) Administration of (2*R*,6*R*)-HNK, but not (*R*,*S*)-ketamine, 3 days before CFC decreased depressive-like behavior on day 2 of the FST in female mice. (n = 5-10 female mice per group). Error bars represent ± SEM. * $p < 0.05$, ** $p < 0.01$. CFC, contextual fear conditioning; sal, saline; K, (*R*,*S*)-ketamine; (2*R*,6*R*)-HNK, (2*R*,6*R*)-hydroxynorketamine; FST, forced swim test.

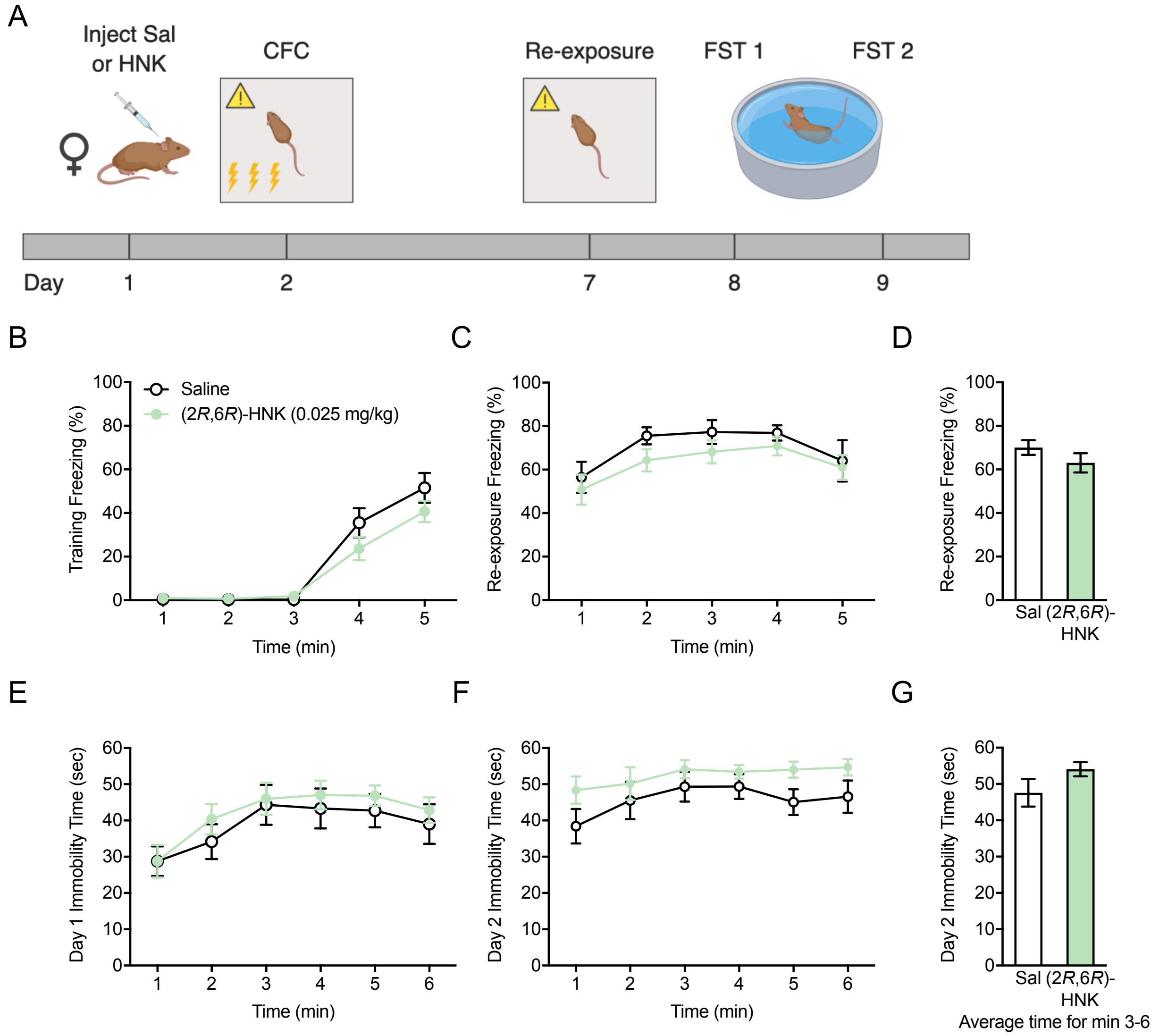

**Figure S07.** Administration of (2*R*,6*R*)-HNK 1 day prior to CFC does not attenuate learned fear or prevent stress-induced depressive-like behavior in female mice. (**A**) Experimental design. Female mice were given a single injection of saline or (2*R*,6*R*)-HNK (0.025 mg/kg) 24 hours prior to 3-shock CFC. Five days later, mice were re-exposed to the training context. Mice were then tested in 2 days of the FST. (**B**) Both groups of mice increased freezing behavior following shocks. (**C-D**) Both groups of mice froze comparably during CFC re-expsoure. (**E**) Both groups exhibited comparable immobility time during day 1 of the FST. (**F-G**) Both groups had comparable immobility time during day 2 of the FST. (n = 9-10 female mice per group). Error bars represent ± SEM. CFC, contextual fear conditioning; sal, saline; (2*R*,6*R*)-HNK, (2*R*,6*R*)-hydroxynorketamine; FST, forced swim test.

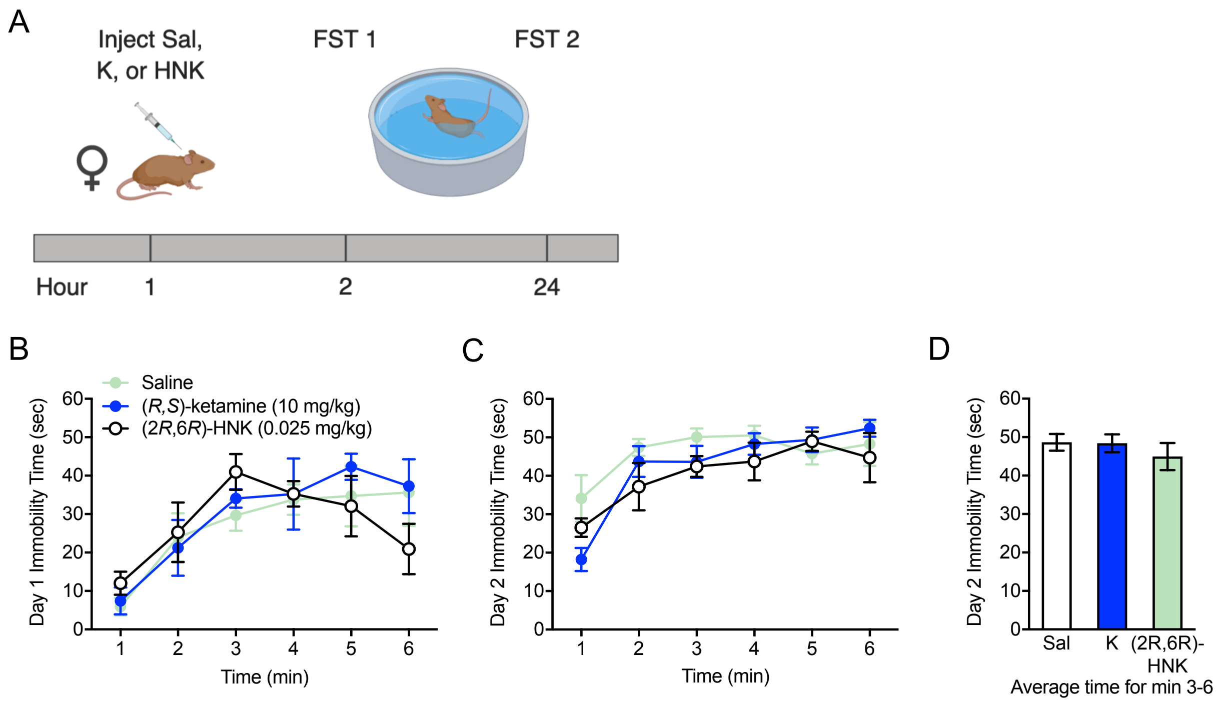

**Figure S08.** Administration of (*R*,*S*)-ketamine or (2*R*,6*R*)-HNK in naïve, non-stressed female mice does not produce an antidepressant-like effect in the FST. (**A**) Experimental design. Female mice were adminstered a single injection of saline, (*R,S*)-ketamine (10 mg/kg) or (2*R*,6*R*)-HNK (0.025 mg/kg) 1 hour prior to Day 1 of the FST. One day later, mice were then administered Day 2 of the FST. (**B**) All groups had comparable immobility time during day 1 of the FST. (**C-D**) All groups had comparable immobility time during day 2 of the FST. (n = 5 female mice per group). Error bars represent ± SEM. Sal, saline; K, (*R*,*S*)-ketamine; (2*R*,6*R*)-HNK, (2*R*,6*R*)-hydroxynorketamine; FST, forced swim test.

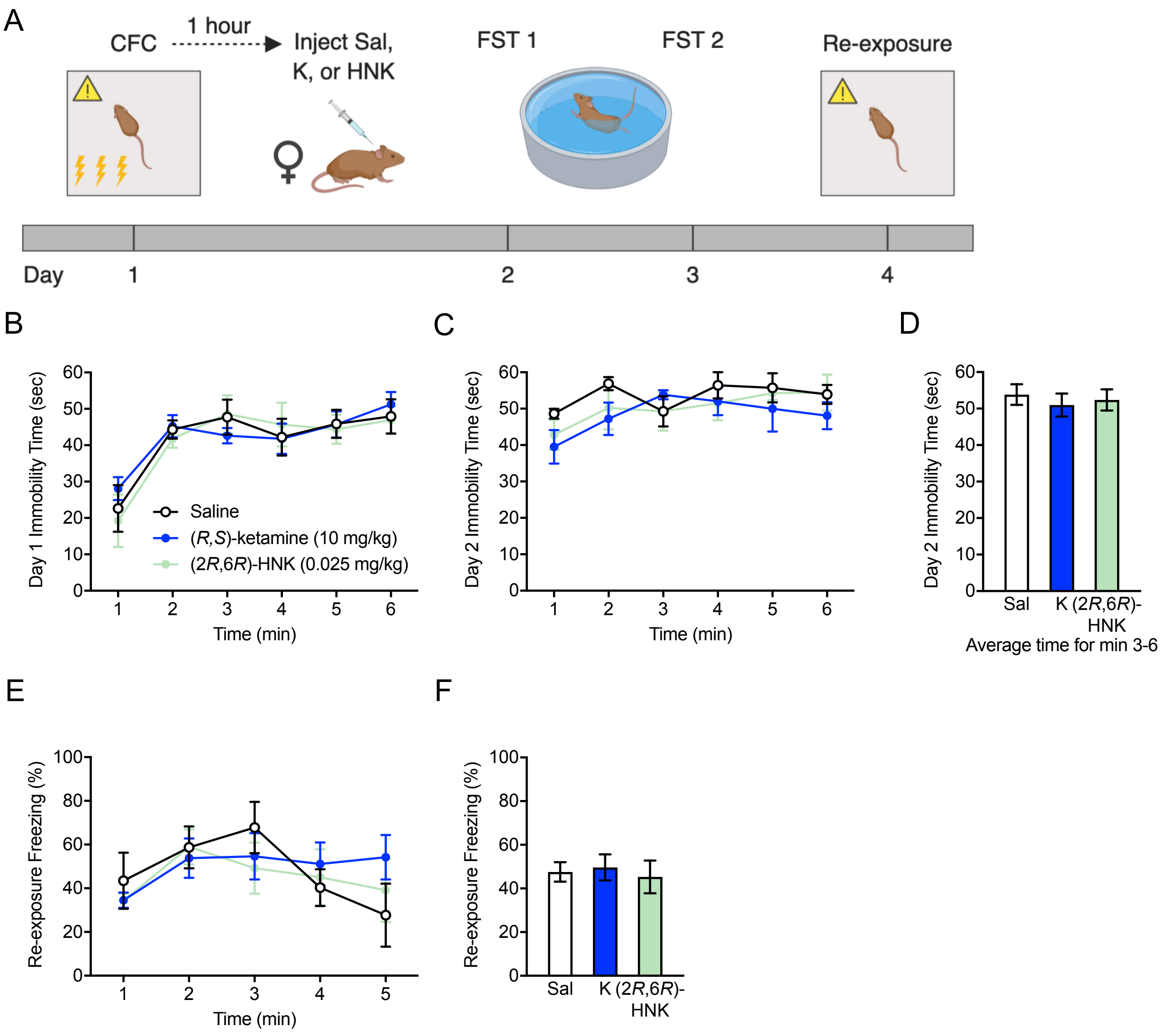

**Figure S09.** Administration of (*R*,*S*)-ketamine or (2*R*,6*R*)-HNK immediately following CFC does not attenuate learned fear or prevent stress-induced depressive-like behavior in female mice. (**A**) Experimental design. Female mice were given 3-shock CFC. One hour later, female mice were administered of a single injection of saline, (*R,S*)-ketamine (10 mg/kg), or (2*R*,6*R*)-HNK (0.025 mg/gk). Mice were then administered 2 days of the FST followed by re-exposure to the aversive CFC training context. (**B**) All groups of mice had comparable immobility time during day 1 of the FST. (**C-D**) All groups of mice had comparable immobility time during day 2 of the FST. (**E-F**) All groups of mice froze comparably during CFC re-exposure. (n = 4-5 female mice per group). Error bars represent ± SEM. CFC, contextual fear conditioning; sal, saline; K, (*R*,*S*)-ketamine; (2*R*,6*R*)-HNK, (2*R*,6*R*)-hydroxynorketamine; FST, forced swim test.

**Table S01.** Statistical analysis summary.

**Table S01: Statistical analysis**

| Cohort | Behavioral Paradigm | Abbrev | Measurement | Statistical Test | Comparison | F | ° of freedom | p | * | Fig. |
|---|---|---|---|---|---|---|---|---|---|---|
| 1 week prophylactic drug, male | Contextual Fear Conditioning Re-exposure | CFC Re-exposure | Freezing (Average) | *t*-test | Sal vs. K (30) | - | - | 0.0042 | ** | 1B |
| | | | | ANOVA | Drug | 0.703 | 7,72 | 0.6698 | - | 1C |
| | | | | ANOVA | Drug | 5.618 | 7,72 | <0.0001 | *** | 1D |
| | | | | Dunnett | Sal vs. (2S,6S)-HNK (0.025) | - | - | 0.0017 | ** | |
| | | | | | Sal vs. (2S,6S)-HNK (0.075) | - | - | 0.0148 | * | |
| | | | | | Sal vs. (2S,6S)-HNK (0.1) | - | - | <0.0001 | *** | |
| | | | | | Sal vs. (2S,6S)-HNK (0.3) | - | - | <0.0001 | *** | |
| | | | | | Sal vs. (2S,6S)-HNK (2.5) | - | - | 0.1484 | - | |
| | | | | | Sal vs. (2S,6S)-HNK (10) | - | - | 0.0490 | * | |
| | | | | | Sal vs. (2S,6S)-HNK (30) | - | - | 0.0122 | * | |
| | Forced Swim Test Day 2 | FST Day 2 | Immobility Time Average (min 3-6) | *t*-test | Sal vs. K (30) | - | - | <0.0001 | *** | 1E |
| | | | | ANOVA | Drug | 2.260 | 7,72 | 0.0388 | * | 1F |
| | | | | Dunnett | Sal vs. (2R,6R)-HNK (0.025) | - | - | 0.6113 | - | |
| | | | | | Sal vs. (2R,6R)-HNK (0.075) | - | - | 0.0248 | * | |
| | | | | | Sal vs. (2R,6R)-HNK (0.1) | - | - | 0.1149 | - | |
| | | | | | Sal vs. (2R,6R)-HNK (0.3) | - | - | 0.1654 | - | |
| | | | | | Sal vs. (2R,6R)-HNK (2.5) | - | - | 0.5103 | - | |
| | | | | | Sal vs. (2R,6R)-HNK (10) | - | - | 0.9978 | - | |
| | | | | | Sal vs. (2R,6R)-HNK (30) | - | - | 0.9998 | - | |
| | | | | ANOVA | Drug | 2.025 | 7,72 | 0.0656 | - | 1G |
| 1 week prophylactic | Contextual Fear Conditioning Re-exposure | CFC Re-exposure | Freezing (Average) | ANOVA | Drug | 0.767 | 3,69 | 0.5166 | - | 2B |
| | | | | ANOVA | Drug | 1.815 | 6,67 | 0.1093 | - | 2C |
| | | | | ANOVA | Drug | 2.939 | 6,65 | 0.0134 | * | 2D |
| | | | | Dunnett | Sal vs. (2S,6S)-HNK (0.025) | - | - | 0.7553 | - | |
| | | | | | Sal vs. (2S,6S)-HNK (0.075) | - | - | 0.1069 | - | |
| | | | | | Sal vs. (2S,6S)-HNK (0.1) | - | - | 0.9988 | - | |
| | | | | | Sal vs. (2S,6S)-HNK (0.3) | - | - | 0.0221 | * | |
| | | | | | Sal vs. (2S,6S)-HNK (2.5) | - | - | 0.9955 | - | |
| | | | | | Sal vs. (2S,6S)-HNK (10) | - | - | 0.7497 | - | |
| | Forced Swim Test Day 2 | FST Day 2 | Immobility Time Average (min 3-6) | ANOVA | Drug | 4.564 | 3,69 | 0.0056 | ** | 2E |
| | | | | Dunnett | Sal vs. K (2.5) | - | - | 0.8869 | - | |
| | | | | | Sal vs. K (10) | - | - | 0.0073 | ** | |
| | | | | | Sal vs. K (30) | - | - | 0.3190 | - | |
| | | | | ANOVA | Drug | 3.494 | 6,67 | 0.0046 | ** | 2F |
| | | | | Dunnett | Sal vs. (2R,6R)-HNK (0.025) | - | - | 0.0085 | ** | |
| | | | | | Sal vs. (2R,6R)-HNK (0.075) | - | - | 0.5884 | - | |
| | | | | | Sal vs. (2R,6R)-HNK (0.1) | - | - | 0.9949 | - | |
| | | | | | Sal vs. (2R,6R)-HNK (0.3) | - | - | 0.9946 | - | |
| | | | | | Sal vs. (2R,6R)-HNK (2.5) | - | - | 0.7349 | - | |
| | | | | | Sal vs. (2R,6R)-HNK (10) | - | - | 0.9996 | - | |
| | | | | ANOVA | Drug | 0.524 | 6,65 | 0.7881 | - | 2G |
| | | | Distance Travelled (min 1-10) | RMANOVA | Drug | 0.530 | 2,12 | 0.6017 | | |
| | | | | | Distance | 21.192 | 9,108 | <0.0001 | *** | |
| | | | | | Drug x Distance | 1.746 | 18,108 | 0.0419 | * | |
| | | | Distance Travelled (min 1) | Dunnett | Sal vs. K (10) | - | - | 0.9834 | - | |
| | | | | | Sal vs. (2R,6R)-HNK (0.025) | - | - | 0.6746 | - | |
| | | | Distance Travelled | Dunnett | Sal vs. K (10) | - | - | 0.2061 | - | |

| | | | | | | | | | | |
|---|---|---|---|---|---|---|---|---|---|---|
| drug, female | Open Field | OF | (min 2) | Dunnett | Sal vs. (2*R*,6*R*)-HNK (0.025) | - | - | 0.1509 | - | 2H |
| | | | Distance Travelled (min 3) | Dunnett | Sal vs. K (10) | - | - | 0.7863 | - | |
| | | | | | Sal vs. (2*R*,6*R*)-HNK (0.025) | - | - | 0.4882 | - | |
| | | | Distance Travelled (min 4) | Dunnett | Sal vs. K (10) | - | - | 0.9842 | - | |
| | | | | | Sal vs. (2*R*,6*R*)-HNK (0.025) | - | - | 0.3716 | - | |
| | | | Distance Travelled (min 5) | Dunnett | Sal vs. K (10) | - | - | 0.6660 | - | |
| | | | | | Sal vs. (2*R*,6*R*)-HNK (0.025) | - | - | 0.7061 | - | |
| | | | Distance Travelled (min 6) | Dunnett | Sal vs. K (10) | - | - | 0.3882 | - | |
| | | | | | Sal vs. (2*R*,6*R*)-HNK (0.025) | - | - | 0.9248 | - | |
| | | | Distance Travelled (min 7) | Dunnett | Sal vs. K (10) | - | - | 0.7061 | - | |
| | | | | | Sal vs. (2*R*,6*R*)-HNK (0.025) | - | - | 0.8253 | - | |
| | | | Distance Travelled (min 8) | Dunnett | Sal vs. K (10) | - | - | 0.4614 | - | |
| | | | | | Sal vs. (2*R*,6*R*)-HNK (0.025) | - | - | 0.3882 | - | |
| | | | Distance Travelled (min 9) | Dunnett | Sal vs. K (10) | - | - | 0.8304 | - | |
| | | | | | Sal vs. (2*R*,6*R*)-HNK (0.025) | - | - | 0.4381 | - | |
| | | | Distance Travelled (min 10) | Dunnett | Sal vs. K (10) | - | - | 0.9795 | - | |
| | | | | | Sal vs. (2*R*,6*R*)-HNK (0.025) | - | - | 0.2968 | - | |
| | | | Distance Travelled Average (min 1-10) | ANOVA | Drug | 0.530 | 2,12 | 0.6017 | - | 2I |
| | | | Time in Center/(Time in Center + Time in Periphery) | ANOVA | Drug | 0.295 | 2,12 | 0.7500 | - | 2J |
| | Tail Immersion Test | TI | Withdrawal Latency | ANOVA | Drug | 2.689 | 2,12 | 0.1084 | - | 2K |
| 1 week prophylactic drug, learned helplessness stress | Learned Helplessness | LH | Session Length | ANOVA | Drug | 1.021 | 2,35 | 0.3707 | - | 3B |
| | | | Escape Latency (trials 1-30) | RMANOVA | Drug | 1.009 | 2,35 | 0.3751 | - | 3C |
| | | | | | Trial | 9.820 | 291,015 | <0.0001 | *** | |
| | | | | | Drug x Trial | 0.949 | 581,015 | 0.5843 | - | |
| | | | Escape Latency Average (trials 11-30) | ANOVA | Drug | 0.657 | 2,35 | 0.5248 | - | 3D |
| | Forced Swim Test Day 1 | FST Day 1 | Immobility Time (min 1-6) | RMANOVA | Drug | 19.929 | 2,130 | <0.0001 | *** | 3E |
| | | | | | Time | 35,450 | 5,130 | <0.0001 | *** | |
| | | | | | Drug x Time | 0.909 | 10,130 | 0.5271 | - | |
| | | | | Dunnett | Sal vs. K (10) | - | - | <0.0001 | *** | |
| | | | | | Sal vs. (2*R*,6*R*)-HNK (0.025) | - | - | <0.0001 | *** | |
| | | | Immobility Time Average (min 3-6) | ANOVA | Drug | 15.04 | 2,26 | <0.0001 | *** | data not shown |
| | | | | Dunnett | Sal vs. K (10) | - | - | <0.0001 | *** | |
| | | | | | Sal vs. (2*R*,6*R*)-HNK (0.025) | - | - | 0.0003 | ** | |
| | Forced Swim Test Day 2 | FST Day 2 | Immobility Time (min 1-6) | RMANOVA | Drug | 16.452 | 2,130 | <0.0001 | *** | 3F |
| | | | | | Time | 14.876 | 5,130 | <0.0001 | *** | |
| | | | | | Drug x Time | 0.701 | 10,130 | 0.7218 | - | |
| | | | | Dunnett | Sal vs. K (10) | - | - | <0.0001 | *** | |
| | | | | | Sal vs. (2*R*,6*R*)-HNK (0.025) | - | - | <0.0001 | *** | |
| | | | Immobility Time Average (min 3-6) | ANOVA | Drug | 15.061 | 2,26 | <0.0001 | *** | 3G |
| | | | | Dunnett | Sal vs. K (10) | - | - | 0.0001 | *** | |
| | | | | | Sal vs. (2*R*,6*R*)-HNK (0.025) | - | - | <0.0001 | *** | |
| | | | Distance Travelled (min 1-6) | RMANOVA | Drug | 1.448 | 2,12 | 0.2733 | - | 3H |
| | | | | | Time | 3.277 | 5,60 | 0.0111 | * | |

| | | | | | | | | | | |
|---|---|---|---|---|---|---|---|---|---|---|
| | | | (min 1-6) | | Drug x Time | 1.404 | 10,60 | 0.2009 | - | |
| | Elevated Plus Maze | EPM | Time in Closed Arms Average (min 1-6) | ANOVA | Drug | 3.777 | 2,12 | 0.0534 | * | 3I |
| | | | | Dunnett | Sal vs. K (10) | - | - | 0.0338 | * | |
| | | | | | Sal vs. (2R,6R)-HNK (0.025) | - | - | 0.5099 | - | |
| | | | Time in Open Arms Average (min 1-6) | ANOVA | Drug | 5.438 | 2,12 | 0.0208 | * | 3J |
| | | | | Dunnett | Sal vs. K (10) | - | - | 0.0147 | * | |
| | | | | | Sal vs. (2R,6R)-HNK (0.025) | - | - | 0.9855 | - | |
| | | | Entries into Open Arms | ANOVA | Drug | 3.352 | 2,12 | 0.0697 | - | 3K |
| 1 week prophylactic drug, chronic immobilization stress | Forced Swim Test Day 1 | FST Day 1 | Immobility Time (min 1-6) | RMANOVA | Drug | 64.951 | 2,26 | <0.0001 | *** | 4B |
| | | | | | Time | 12.521 | 5,130 | <0.0001 | *** | |
| | | | | | Drug x Time | 2.932 | 10,130 | 0.0024 | ** | |
| | | | | Dunnett | Sal vs. K (10) | - | - | 0.6158 | - | |
| | | | | | Sal vs. (2R,6R)-HNK (0.025) | - | - | <0.0001 | *** | |
| | | | Immobility Time (min 1) | Dunnett | Sal vs. K (10) | - | - | 0.3936 | - | |
| | | | | | Sal vs. (2R,6R)-HNK (0.025) | - | - | <0.0001 | *** | |
| | | | Immobility Time (min 2) | Dunnett | Sal vs. K (10) | - | - | 0.4423 | - | |
| | | | | | Sal vs. (2R,6R)-HNK (0.025) | - | - | <0.0001 | *** | |
| | | | Immobility Time (min 3) | Dunnett | Sal vs. K (10) | - | - | 0.7582 | - | |
| | | | | | Sal vs. (2R,6R)-HNK (0.025) | - | - | <0.0001 | *** | |
| | | | Immobility Time (min 4) | Dunnett | Sal vs. K (10) | - | - | 0.6097 | - | |
| | | | | | Sal vs. (2R,6R)-HNK (0.025) | - | - | <0.0001 | *** | |
| | | | Immobility Time (min 5) | Dunnett | Sal vs. K (10) | - | - | 0.8327 | - | |
| | | | | | Sal vs. (2R,6R)-HNK (0.025) | - | - | <0.0001 | *** | |
| | | | Immobility Time (min 6) | Dunnett | Sal vs. K (10) | - | - | 0.0014 | ** | |
| | | | | | Sal vs. (2R,6R)-HNK (0.025) | - | - | <0.0001 | *** | |
| | | | Immobility Time Average (min 1-6) | ANOVA | Drug | 79.304 | 2,26 | <0.0001 | *** | |
| | | | | Dunnett | Sal vs. K (10) | - | - | 0.3118 | - | |
| | | | | | Sal vs. (2R,6R)-HNK (0.025) | - | - | <0.0001 | *** | |
| | Forced Swim Test Day 2 | FST Day 2 | Immobility Time (min 1-6) | RMANOVA | Drug | 8.249 | 2,130 | 0.0017 | ** | 4C |
| | | | | | Time | 13.472 | 5,130 | <0.0001 | *** | |
| | | | | | Drug x Time | 3.528 | 10,130 | 0.0004 | ** | |
| | | | | Dunnett | Sal vs. K (10) | - | - | 0.0012 | ** | |
| | | | | | Sal vs. (2R,6R)-HNK (0.025) | - | - | 0.5892 | - | |
| | | | Immobility Time (min 1) | Dunnett | Sal vs. K (10) | - | - | 0.0608 | - | |
| | | | | | Sal vs. (2R,6R)-HNK (0.025) | - | - | 0.0057 | ** | |
| | | | Immobility Time (min 2) | Dunnett | Sal vs. K (10) | - | - | 0.0156 | * | |
| | | | | | Sal vs. (2R,6R)-HNK (0.025) | - | - | 0.9408 | - | |
| | | | Immobility Time (min 3) | Dunnett | Sal vs. K (10) | - | - | 0.0236 | * | |
| | | | | | Sal vs. (2R,6R)-HNK (0.025) | - | - | 0.3496 | - | |
| | | | Immobility Time (min 4) | Dunnett | Sal vs. K (10) | - | - | 0.0792 | - | |
| | | | | | Sal vs. (2R,6R)-HNK (0.025) | - | - | 0.3042 | - | |
| | | | Immobility Time (min 5) | Dunnett | Sal vs. K (10) | - | - | 0.0152 | * | |
| | | | | | Sal vs. (2R,6R)-HNK (0.025) | - | - | 0.1760 | - | |
| | | | Immobility Time (min 6) | Dunnett | Sal vs. K (10) | - | - | 0.0016 | ** | |
| | | | | | Sal vs. (2R,6R)-HNK (0.025) | - | - | 0.0222 | * | |
| | | | Immobility Time Average (min 3-6) | ANOVA | Drug | 8.115 | 2,26 | 0.0018 | ** | 4D |
| | | | | Dunnett | Sal vs. K (10) | - | - | 0.0010 | ** | |
| | | | | | Sal vs. (2R,6R)-HNK (0.025) | - | - | 0.0328 | * | |
| | Contextual Fear Conditioning Training | CFC Training | Freezing (min 1-5) | RMANOVA | Drug | 1.065 | 2,104 | 0.3592 | - | 4E |
| | | | | | Time | 55.948 | 4,104 | <0.0001 | *** | |

| | | | | | | | | | | |
|---|---|---|---|---|---|---|---|---|---|---|
| | Conditioning Training | | | | Drug x Time | 0.529 | 8,104 | 0.8323 | - | |
| | Contextual Fear Conditioning Re-exposure | CFC Re-exposure | Freezing (min 1-5) | RMANOVA | Drug | 0.913 | 2,104 | 0.4140 | - | 4F |
| | | | | | Time | 15.611 | 4,104 | <0.0001 | *** | |
| | | | | | Drug x Time | 1.048 | 8,104 | 0.4056 | - | |
| | | | Freezing Average | ANOVA | Drug | 0.913 | 2,26 | 0.4140 | - | 4G |
| | Contextual Fear Conditioning Training | CFC Training | Freezing (min 1-5) | RMANOVA | Surgery | 2.482 | 1,77 | 0.1193 | - | 5B |
| | | | | | Drug | 1.164 | 2,77 | 0.3175 | - | |
| | | | | | Surgery x Drug | 1.765 | 2,77 | 0.1780 | - | |
| | | | | | Time | 172.273 | 4,308 | <0.0001 | *** | |
| | | | | | Surgery x Time | 1.581 | 4,308 | 0.1791 | - | |
| | | | | | Drug x Time | 1.522 | 8,308 | 0.1488 | - | |
| | | | | | Surgery x Drug x Time | 1.514 | 8,308 | 0.1515 | - | |
| | Contextual Fear Conditioning Re-exposure | CFC Re-exposure | Freezing (min 1-5) | RMANOVA | Surgery | 0.002 | 1,77 | 0.9667 | - | 5C |
| | | | | | Drug | 4.751 | 2,77 | 0.0113 | * | |
| | | | | | Surgery x Drug | 0.649 | 2,77 | 0.5252 | - | |
| | | | | | Time | 21.391 | 4,308 | <0.0001 | *** | |
| | | | | | Surgery x Time | 2.126 | 4,308 | 0.0775 | - | |
| | | | | | Drug x Time | 0.488 | 8,308 | 0.8647 | - | |
| | | | | | Surgery x Drug x Time | 0.640 | 8,308 | 0.7437 | - | |
| | | | | Tukey | Sham Sal vs. Sham K (10) | - | - | 0.1526 | - | |
| | | | | | Sham Sal vs. Sham (2*R*,6*R*)-HNK (0.025) | - | - | 0.1280 | - | |
| | | | | | OVX Sal vs. OVX K (10) | - | - | 0.7566 | - | |
| | | | | | OVX Sal vs. OVX (2*R*,6*R*)-HNK (0.025) | - | - | 0.8664 | - | |
| | | | | | Sham Sal vs. OVX Sal | - | - | 0.9480 | - | |
| | | | | | Sham K (10) vs. OVX K (10) | - | - | 0.9996 | - | |
| | | | | | Sham (2*R*,6*R*)-HNK (0.025) vs. OVX (2*R*,6*R*)-HNK | - | - | 0.9876 | - | |
| | | | Freezing Average (min 1-5) | ANOVA | Surgery | 0.00175 | 1,77 | 0.9667 | - | 5D |
| | | | | | Drug | 4.751 | 2,77 | 0.0113 | * | |
| | | | | | Surgery x Drug | 0.6494 | 2,77 | 0.5252 | - | |
| | | | | Sidak | Sham Sal vs. Sham K (10) | - | - | 0.2218 | - | |
| | | | | | Sham Sal vs. Sham (2*R*,6*R*)-HNK (0.025) | - | - | 0.1835 | - | |
| | | | | | OVX Sal vs. OVX K (10) | - | - | 0.9501 | - | |
| | | | | | OVX Sal vs. OVX (2*R*,6*R*)-HNK (0.025) | - | - | 0.9893 | - | |
| | | | | | Sham Sal vs. OVX Sal | - | - | 0.9992 | - | |
| | | | | | Sham K (10) vs. OVX K (10) | - | - | >0.9999 | - | |
| | | | | | Sham (2*R*,6*R*)-HNK (0.025) vs. OVX (2*R*,6*R*)-HNK | - | - | >0.9999 | - | |
| | | | Immobility Time (min 1-6) | RMANOVA | Surgery | 2.727 | 1,77 | 0.1028 | - | |
| | | | | | Drug | 0.795 | 2,77 | 0.4554 | - | |
| | | | | | Surgery x Drug | 9.314 | 2,77 | 0.0002 | ** | |
| | | | | | Time | 113.921 | 5,385 | <0.0001 | *** | |
| | | | | | Surgery x Time | 4.092 | 5,385 | 0.0012 | ** | |
| | | | | | Drug x Time | 1.330 | 10,385 | 0.2122 | - | |
| | | | | | Surgery x Drug x Time | 1.073 | 10,385 | 0.3823 | - | |
| | | | | Tukey | Sham Sal vs. Sham K (10) | - | - | 0.0384 | * | |
| | | | | | Sham Sal vs. Sham (2*R*,6*R*)-HNK (0.025) | - | - | 0.9978 | - | |
| | | | | | OVX Sal vs. OVX K (10) | - | - | 0.7164 | - | |
| | | | | | OVX Sal vs. OVX (2*R*,6*R*)-HNK (0.025) | - | - | 0.9309 | - | |
| | | | | | Sham Sal vs. OVX Sal | - | - | 0.3483 | - | |
| | | | | | Sham K (10) vs. OVX K (10) | - | - | 0.1455 | - | |
| | | | | | Sham (2*R*,6*R*)-HNK (0.025) vs. OVX (2*R*,6*R*)-HNK | - | - | 0.0139 | * | |
| | | | Immobility Time (min | | Sham Sal vs. Sham K (10) | - | - | 0.4345 | - | |
| | | | | | Sham Sal vs. Sham (2*R*,6*R*)-HNK (0.025) | - | - | 0.9960 | - | |
| | | | | | OVX Sal vs. OVX K (10) | - | - | 0.9982 | - | |

| | | | | | | | | | | |
|---|---|---|---|---|---|---|---|---|---|---|
| Ovariectomized, 1 week prophylactic drug | Forced Swim Test Day 1 | FST Day 1 | Immobility Time (min 1) | Tukey | OVX Sal vs. OVX (2*R*,6*R*)-HNK (0.025) | - | - | >0.9999 | - | 5E |
| | | | | | Sham Sal vs. OVX Sal | - | - | 0.0413 | * | |
| | | | | | Sham K (10) vs. OVX K (10) | - | - | 0.9918 | - | |
| | | | | | Sham (2*R*,6*R*)-HNK (0.025) vs. OVX (2*R*,6*R*)-HNK | - | - | 0.0078 | ** | |
| | | | Immobility Time (min 2) | Tukey | Sham Sal vs. Sham K (10) | - | - | 0.1975 | - | |
| | | | | | Sham Sal vs. Sham (2*R*,6*R*)-HNK (0.025) | - | - | 0.6026 | - | |
| | | | | | OVX Sal vs. OVX K (10) | - | - | 0.9967 | - | |
| | | | | | OVX Sal vs. OVX (2*R*,6*R*)-HNK (0.025) | - | - | 0.7070 | - | |
| | | | | | Sham Sal vs. OVX Sal | - | - | 0.2750 | - | |
| | | | | | Sham K (10) vs. OVX K (10) | - | - | 0.1090 | - | |
| | | | | | Sham (2*R*,6*R*)-HNK (0.025) vs. OVX (2*R*,6*R*)-HNK | - | - | 0.0141 | * | |
| | | | Immobility Time (min 3) | Tukey | Sham Sal vs. Sham K (10) | - | - | 0.0058 | ** | |
| | | | | | Sham Sal vs. Sham (2*R*,6*R*)-HNK (0.025) | - | - | 0.9998 | - | |
| | | | | | OVX Sal vs. OVX K (10) | - | - | 0.9898 | - | |
| | | | | | OVX Sal vs. OVX (2*R*,6*R*)-HNK (0.025) | - | - | 0.5322 | - | |
| | | | | | Sham Sal vs. OVX Sal | - | - | 0.7722 | - | |
| | | | | | Sham K (10) vs. OVX K (10) | - | - | 0.0452 | * | |
| | | | | | Sham (2*R*,6*R*)-HNK (0.025) vs. OVX (2*R*,6*R*)-HNK | - | - | 0.0795 | - | |
| | | | Immobility Time (min 4) | Tukey | Sham Sal vs. Sham K (10) | - | - | 0.0921 | - | |
| | | | | | Sham Sal vs. Sham (2*R*,6*R*)-HNK (0.025) | - | - | 0.9993 | - | |
| | | | | | OVX Sal vs. OVX K (10) | - | - | 0.9034 | - | |
| | | | | | OVX Sal vs. OVX (2*R*,6*R*)-HNK (0.025) | - | - | 0.9910 | - | |
| | | | | | Sham Sal vs. OVX Sal | - | - | 0.7443 | - | |
| | | | | | Sham K (10) vs. OVX K (10) | - | - | 0.1822 | - | |
| | | | | | Sham (2*R*,6*R*)-HNK (0.025) vs. OVX (2*R*,6*R*)-HNK | - | - | 0.6080 | - | |
| | | | Immobility Time (min 5) | Tukey | Sham Sal vs. Sham K (10) | - | - | 0.2246 | - | |
| | | | | | Sham Sal vs. Sham (2*R*,6*R*)-HNK (0.025) | - | - | >0.9999 | - | |
| | | | | | OVX Sal vs. OVX K (10) | - | - | 0.7393 | - | |
| | | | | | OVX Sal vs. OVX (2*R*,6*R*)-HNK (0.025) | - | - | 0.9495 | - | |
| | | | | | Sham Sal vs. OVX Sal | - | - | 0.5877 | - | |
| | | | | | Sham K (10) vs. OVX K (10) | - | - | 0.3357 | - | |
| | | | | | Sham (2*R*,6*R*)-HNK (0.025) vs. OVX (2*R*,6*R*)-HNK | - | - | 0.1123 | - | |
| | | | Immobility Time (min 6) | Tukey | Sham Sal vs. Sham K (10) | - | - | 0.5360 | - | |
| | | | | | Sham Sal vs. Sham (2*R*,6*R*)-HNK (0.025) | - | - | 0.9954 | - | |
| | | | | | OVX Sal vs. OVX K (10) | - | - | 0.0709 | - | |
| | | | | | OVX Sal vs. OVX (2*R*,6*R*)-HNK (0.025) | - | - | >0.9999 | - | |
| | | | | | Sham Sal vs. OVX Sal | - | - | 0.3641 | - | |
| | | | | | Sham K (10) vs. OVX K (10) | - | - | 0.1486 | - | |
| | | | | | Sham (2*R*,6*R*)-HNK (0.025) vs. OVX (2*R*,6*R*)-HNK | - | - | 0.1679 | - | |
| | | | Immobility Time Average (min 3-6) | ANOVA | Surgery | 0.853 | 1,77 | 0.3586 | - | data not shown |
| | | | | | Drug | 0.469 | 2,77 | 0.6277 | - | |
| | | | | | Surgery x Drug | 9.029 | 2,77 | 0.0003 | * | |
| | | | | Sidak | Sham Sal vs. Sham K (10) | | | 0.0447 | * | |
| | | | | | Sham Sal vs. Sham (2*R*,6*R*)-HNK (0.025) | | | >0.9999 | - | |
| | | | | | OVX Sal vs. OVX K (10) | | | 0.5082 | - | |
| | | | | | OVX Sal vs. OVX (2*R*,6*R*)-HNK (0.025) | | | 0.9462 | - | |
| | | | | | Sham Sal vs. OVX Sal | | | 0.4448 | - | |
| | | | | | Sham K (10) vs. OVX K (10) | | | 0.0546 | - | |
| | | | | | Sham (2*R*,6*R*)-HNK (0.025) vs. OVX (2*R*,6*R*)-HNK | | | 0.0867 | - | |
| | | | | RMANOVA | Surgery | 0.806 | 1,77 | 0.3721 | - | |
| | | | | | Drug | 4.979 | 2,77 | 0.0093 | ** | |
| | | | | | Surgery x Drug | 3.116 | 2,77 | 0.0500 | - | |
| | | | | | Time | 30.320 | 5,385 | <0.0001 | *** | |
| | | | | | Surgery x Time | 7867.000 | 5,385 | 0.0992 | - | |

| | | | | | | | | | | |
|---|---|---|---|---|---|---|---|---|---|---|
| | Forced Swim Test Day 2 | FST Day 2 | Immobility Time (min 1-6) | | Drug x Time | 1.614 | 10,385 | 0.1003 | - | 5F |
| | | | | | Surgery x Drug x Time | 1.498 | 10,385 | 0.1376 | - | |
| | | | | Tukey | Sham Sal vs. Sham K (10) | - | - | 0.0473 | * | |
| | | | | | Sham Sal vs. Sham (2*R*,6*R*)-HNK (0.025) | - | - | 0.0077 | ** | |
| | | | | | OVX Sal vs. OVX K (10) | - | - | >0.9999 | - | |
| | | | | | OVX Sal vs. OVX (2*R*,6*R*)-HNK (0.025) | - | - | 0.9831 | - | |
| | | | | | Sham Sal vs. OVX Sal | - | - | 0.6508 | - | |
| | | | | | Sham K (10) vs. OVX K (10) | - | - | 0.6391 | - | |
| | | | | | Sham (2*R*,6*R*)-HNK (0.025) vs. OVX (2*R*,6*R*)-HNK | - | - | 0.6526 | - | |
| | | | Immobility Time Average (min 3-6) | ANOVA | Surgery | 3.914 | 1,74 | 0.0516 | - | 5G |
| | | | | | Drug | 3.980 | 2,74 | 0.0228 | * | |
| | | | | | Surgery x Drug | 4.907 | 2,74 | 0.0100 | * | |
| | | | | Sidak | Sham Sal vs. Sham K (10) | - | - | 0.0092 | ** | |
| | | | | | Sham Sal vs. Sham (2*R*,6*R*)-HNK (0.025) | - | - | 0.0225 | * | |
| | | | | | OVX Sal vs. OVX K (10) | - | - | >0.9999 | - | |
| | | | | | OVX Sal vs. OVX (2*R*,6*R*)-HNK (0.025) | - | - | >0.9999 | - | |
| | | | | | Sham Sal vs. OVX Sal | - | - | 0.9335 | - | |
| | | | | | Sham K (10) vs. OVX K (10) | - | - | 0.1466 | - | |
| | | | | | Sham (2*R*,6*R*)-HNK (0.025) vs. OVX (2*R*,6*R*)-HNK | - | - | 0.4752 | - | |
| | | | Freezing (min 1-5) | RMANOVA | Hormone | 11.141 | 2,86 | <0.0001 | *** | |
| | | | | | Drug | 0.833 | 2,86 | 0.4382 | - | |
| | | | | | Hormone x Drug | 0.569 | 4,86 | 0.6861 | - | |
| | | | | | Time | 243.859 | 4,344 | <0.0001 | *** | |
| | | | | | Hormone x Time | 7.942 | 8,344 | <0.0001 | *** | |
| | | | | | Drug x Time | 1.499 | 8,344 | 0.1561 | - | |
| | | | | | Hormone x Drug x Time | 0.721 | 16,344 | 0.7723 | - | |
| | | | | Tukey | Vehicle Sal vs. Vehicle K (10) | - | - | 0.9996 | - | |
| | | | | | Vehicle Sal vs. Vehicle (2*R*,6*R*)-HNK (0.025) | - | - | 0.9986 | - | |
| | | | | | E2 Sal vs. E2 K (10) | - | - | 0.9994 | - | |
| | | | | | E2 Sal vs. E2 (2*R*,6*R*)-HNK (0.025) | - | - | 0.9911 | - | |
| | | | | | E2/P4 Sal vs. E2/P4 K (10) | - | - | 0.9993 | - | |
| | | | | | E2/P4 Sal vs. E2/P4 (2*R*,6*R*)-HNK (0.025) | - | - | >0.9999 | - | |
| | | | | | Vehicle Sal vs. E2 Sal | - | - | 0.9866 | - | |
| | | | | | Vehicle Sal vs. E2/P4 Sal | - | - | 0.1530 | - | |
| | | | | | Vehicle K (10) vs. E2 K (10) | - | - | >0.9999 | - | |
| | | | | | Vehicle K (10) vs. E2/P4 K (10) | - | - | 0.1272 | - | |
| | | | | | Vehicle (2*R*,6*R*)-HNK (0.025) vs. E2 (2*R*,6*R*)-HNK | - | - | 0.7961 | - | |
| | | | | | Vehicle (2*R*,6*R*)-HNK (0.025) vs. E2/P4 (2*R*,6*R*)- | - | - | 0.1960 | - | |
| | | | Freezing (min 1) | Tukey | Vehicle Sal vs. Vehicle K (10) | - | - | >0.9999 | - | |
| | | | | | Vehicle Sal vs. Vehicle (2*R*,6*R*)-HNK (0.025) | - | - | >0.9999 | - | |
| | | | | | E2 Sal vs. E2 K (10) | - | - | >0.9999 | - | |
| | | | | | E2 Sal vs. E2 (2*R*,6*R*)-HNK (0.025) | - | - | >0.9999 | - | |
| | | | | | E2/P4 Sal vs. E2/P4 K (10) | - | - | >0.9999 | - | |
| | | | | | E2/P4 Sal vs. E2/P4 (2*R*,6*R*)-HNK (0.025) | - | - | >0.9999 | - | |
| | | | | | Vehicle Sal vs. E2 Sal | - | - | >0.9999 | - | |
| | | | | | Vehicle Sal vs. E2/P4 Sal | - | - | >0.9999 | - | |
| | | | | | Vehicle K (10) vs. E2 K (10) | - | - | >0.9999 | - | |
| | | | | | Vehicle K (10) vs. E2/P4 K (10) | - | - | >0.9999 | - | |
| | | | | | Vehicle (2*R*,6*R*)-HNK (0.025) vs. E2 (2*R*,6*R*)-HNK | - | - | >0.9999 | - | |
| | | | | | Vehicle (2*R*,6*R*)-HNK (0.025) vs. E2/P4 (2*R*,6*R*)- | - | - | >0.9999 | - | |
| | | | | | Vehicle Sal vs. Vehicle K (10) | - | - | >0.9999 | - | |
| | | | | | Vehicle Sal vs. Vehicle (2*R*,6*R*)-HNK (0.025) | - | - | >0.9999 | - | |
| | | | | | E2 Sal vs. E2 K (10) | - | - | >0.9999 | - | |

| | | | | | | | | | | |
|---|---|---|---|---|---|---|---|---|---|---|
| | Contextual Fear Conditioning Training | CFC Training | Freezing (min 2) | Tukey | E2 Sal vs. E2 (2*R*,6*R*)-HNK (0.025) | - | - | >0.9999 | - | 6B |
| | | | | | E2/P4 Sal vs. E2/P4 K (10) | - | - | >0.9999 | - | |
| | | | | | E2/P4 Sal vs. E2/P4 (2*R*,6*R*)-HNK (0.025) | - | - | >0.9999 | - | |
| | | | | | Vehicle Sal vs. E2 Sal | - | - | >0.9999 | - | |
| | | | | | Vehicle Sal vs. E2/P4 Sal | - | - | >0.9999 | - | |
| | | | | | Vehicle K (10) vs. E2 K (10) | - | - | >0.9999 | - | |
| | | | | | Vehicle K (10) vs. E2/P4 K (10) | - | - | >0.9999 | - | |
| | | | | | Vehicle (2*R*,6*R*)-HNK (0.025) vs. E2 (2*R*,6*R*)-HNK | - | - | >0.9999 | - | |
| | | | | | Vehicle (2*R*,6*R*)-HNK (0.025) vs. E2/P4 (2*R*,6*R*)- | - | - | >0.9999 | - | |
| | | | Freezing (min 3) | Tukey | Vehicle Sal vs. Vehicle K (10) | - | - | >0.9999 | - | |
| | | | | | Vehicle Sal vs. Vehicle (2*R*,6*R*)-HNK (0.025) | - | - | >0.9999 | - | |
| | | | | | E2 Sal vs. E2 K (10) | - | - | >0.9999 | - | |
| | | | | | E2 Sal vs. E2 (2*R*,6*R*)-HNK (0.025) | - | - | >0.9999 | - | |
| | | | | | E2/P4 Sal vs. E2/P4 K (10) | - | - | >0.9999 | - | |
| | | | | | E2/P4 Sal vs. E2/P4 (2*R*,6*R*)-HNK (0.025) | - | - | >0.9999 | - | |
| | | | | | Vehicle Sal vs. E2 Sal | - | - | >0.9999 | - | |
| | | | | | Vehicle Sal vs. E2/P4 Sal | - | - | >0.9999 | - | |
| | | | | | Vehicle K (10) vs. E2 K (10) | - | - | >0.9999 | - | |
| | | | | | Vehicle K (10) vs. E2/P4 K (10) | - | - | >0.9999 | - | |
| | | | | | Vehicle (2*R*,6*R*)-HNK (0.025) vs. E2 (2*R*,6*R*)-HNK | - | - | >0.9999 | - | |
| | | | | | Vehicle (2*R*,6*R*)-HNK (0.025) vs. E2/P4 (2*R*,6*R*)- | - | - | >0.9999 | - | |
| | | | Freezing (min 4) | Tukey | Vehicle Sal vs. Vehicle K (10) | - | - | >0.9999 | - | |
| | | | | | Vehicle Sal vs. Vehicle (2*R*,6*R*)-HNK (0.025) | - | - | >0.9999 | - | |
| | | | | | E2 Sal vs. E2 K (10) | - | - | 0.9902 | - | |
| | | | | | E2 Sal vs. E2 (2*R*,6*R*)-HNK (0.025) | - | - | 0.8009 | - | |
| | | | | | E2/P4 Sal vs. E2/P4 K (10) | - | - | 0.9997 | - | |
| | | | | | E2/P4 Sal vs. E2/P4 (2*R*,6*R*)-HNK (0.025) | - | - | >0.9999 | - | |
| | | | | | Vehicle Sal vs. E2 Sal | - | - | >0.9999 | - | |
| | | | | | Vehicle Sal vs. E2/P4 Sal | - | - | 0.0052 | ** | |
| | | | | | Vehicle K (10) vs. E2 K (10) | - | - | 0.9942 | - | |
| | | | | | Vehicle K (10) vs. E2/P4 K (10) | - | - | 0.0846 | - | |
| | | | | | Vehicle (2*R*,6*R*)-HNK (0.025) vs. E2 (2*R*,6*R*)-HNK | - | - | 0.2439 | - | |
| | | | | | Vehicle (2*R*,6*R*)-HNK (0.025) vs. E2/P4 (2*R*,6*R*)- | - | - | 0.0008 | ** | |
| | | | Freezing (min 5) | Tukey | Vehicle Sal vs. Vehicle K (10) | - | - | 0.6260 | - | |
| | | | | | Vehicle Sal vs. Vehicle (2*R*,6*R*)-HNK (0.025) | - | - | 0.1992 | - | |
| | | | | | E2 Sal vs. E2 K (10) | - | - | 0.9967 | - | |
| | | | | | E2 Sal vs. E2 (2*R*,6*R*)-HNK (0.025) | - | - | 0.8820 | - | |
| | | | | | E2/P4 Sal vs. E2/P4 K (10) | - | - | 0.2576 | - | |
| | | | | | E2/P4 Sal vs. E2/P4 (2*R*,6*R*)-HNK (0.025) | - | - | 0.9130 | - | |
| | | | | | Vehicle Sal vs. E2 Sal | - | - | 0.3177 | - | |
| | | | | | Vehicle Sal vs. E2/P4 Sal | - | - | 0.0022 | ** | |
| | | | | | Vehicle K (10) vs. E2 K (10) | - | - | >0.9999 | - | |
| | | | | | Vehicle K (10) vs. E2/P4 K (10) | - | - | 0.0001 | ** | |
| | | | | | Vehicle (2*R*,6*R*)-HNK (0.025) vs. E2 (2*R*,6*R*)-HNK | - | - | 0.3684 | - | |
| | | | | | Vehicle (2*R*,6*R*)-HNK (0.025) vs. E2/P4 (2*R*,6*R*)- | - | - | 0.0388 | * | |
| | Contextual Fear Conditioning Re-exposure | CFC Re-exposure | Freezing (min 1-5) | RMANOVA | Hormone | 2.254 | 2,6 | 0.1112 | - | 6C |
| | | | | | Drug | 1.205 | 2,86 | 0.3048 | - | |
| | | | | | Hormone x Drug | 0.892 | 4,86 | 0.4721 | - | |
| | | | | | Time | 16.746 | 4,344 | <0.0001 | *** | |
| | | | | | Hormone x Time | 1.038 | 8,344 | 0.4071 | - | |
| | | | | | Drug x Time | 0.490 | 8,344 | 0.8631 | - | |
| | | | | | Hormone x Drug x Time | 0.351 | 16,344 | 0.9912 | - | |
| | | | Freezing Average (min 1-5) | ANOVA | Hormone | 2.254 | 2,86 | 0.1112 | - | 6D |
| | | | | | Drug | 1.205 | 2,86 | 0.3048 | - | |

| | | | | | | | | | | |
|---|---|---|---|---|---|---|---|---|---|---|
| | | | (min 1-5) | | Hormone x Drug | 0.8925 | 4,86 | 0.4721 | - | |
| | | | Immobility Time (min 1-6) | RMANOVA | Hormone | 25.983 | 2,85 | <0.0001 | *** | |
| | | | | | Drug | 2.644 | 2,85 | 0.0769 | - | |
| | | | | | Hormone x Drug | 0.798 | 4,85 | 0.5301 | - | |
| | | | | | Time | 54.540 | 5,425 | <0.0001 | *** | |
| | | | | | Hormone x Time | 2.633 | 10,425 | 0.0040 | ** | |
| | | | | | Drug x Time | 2.493 | 10,425 | 0.0065 | ** | |
| | | | | | Hormone x Drug x Time | 1.897 | 20,425 | 0.0114 | * | |
| | | | | Tukey | Vehicle Sal vs. Vehicle K (10) | - | - | 0.9420 | - | |
| | | | | | Vehicle Sal vs. Vehicle (2R,6R)-HNK (0.025) | - | - | >0.9999 | - | |
| | | | | | E2 Sal vs. E2 K (10) | - | - | 0.9994 | - | |
| | | | | | E2 Sal vs. E2 (2R,6R)-HNK (0.025) | - | - | 0.5359 | - | |
| | | | | | E2/P4 Sal vs. E2/P4 K (10) | - | - | 0.9888 | - | |
| | | | | | E2/P4 Sal vs. E2/P4 (2R,6R)-HNK (0.025) | - | - | 0.9683 | - | |
| | | | | | Vehicle Sal vs. E2 Sal | - | - | 0.2557 | - | |
| | | | | | Vehicle Sal vs. E2/P4 Sal | - | - | 0.6111 | - | |
| | | | | | Vehicle K (10) vs. E2 K (10) | - | - | 0.4394 | - | |
| | | | | | Vehicle K (10) vs. E2/P4 K (10) | - | - | 0.5003 | - | |
| | | | | | Vehicle (2R,6R)-HNK (0.025) vs. E2 (2R,6R)-HNK | - | - | 0.0005 | ** | |
| | | | | | Vehicle (2R,6R)-HNK (0.025) vs. E2/P4 (2R,6R)- | - | - | 0.9191 | - | |
| | | | Immobility Time (min 1) | Tukey | Vehicle Sal vs. Vehicle K (10) | - | - | 0.8342 | - | |
| | | | | | Vehicle Sal vs. Vehicle (2R,6R)-HNK (0.025) | - | - | 0.7769 | - | |
| | | | | | E2 Sal vs. E2 K (10) | - | - | 0.9604 | - | |
| | | | | | E2 Sal vs. E2 (2R,6R)-HNK (0.025) | - | - | 0.8964 | - | |
| | | | | | E2/P4 Sal vs. E2/P4 K (10) | - | - | 0.9331 | - | |
| | | | | | E2/P4 Sal vs. E2/P4 (2R,6R)-HNK (0.025) | - | - | 0.9989 | - | |
| | | | | | Vehicle Sal vs. E2 Sal | - | - | 0.0061 | ** | |
| | | | | | Vehicle Sal vs. E2/P4 Sal | - | - | 0.8647 | - | |
| | | | | | Vehicle K (10) vs. E2 K (10) | - | - | 0.0057 | ** | |
| | | | | | Vehicle K (10) vs. E2/P4 K (10) | - | - | 0.8780 | - | |
| | | | | | Vehicle (2R,6R)-HNK (0.025) vs. E2 (2R,6R)-HNK | - | - | 0.0008 | ** | |
| | | | | | Vehicle (2R,6R)-HNK (0.025) vs. E2/P4 (2R,6R)- | - | - | 0.2246 | - | |
| | | | Immobility Time (min 2) | Tukey | Vehicle Sal vs. Vehicle K (10) | - | - | >0.9999 | - | |
| | | | | | Vehicle Sal vs. Vehicle (2R,6R)-HNK (0.025) | - | - | 0.9820 | - | |
| | | | | | E2 Sal vs. E2 K (10) | - | - | 0.9932 | - | |
| | | | | | E2 Sal vs. E2 (2R,6R)-HNK (0.025) | - | - | 0.9952 | - | |
| | | | | | E2/P4 Sal vs. E2/P4 K (10) | - | - | >0.9999 | - | |
| | | | | | E2/P4 Sal vs. E2/P4 (2R,6R)-HNK (0.025) | - | - | >0.9999 | - | |
| | | | | | Vehicle Sal vs. E2 Sal | - | - | 0.3685 | - | |
| | | | | | Vehicle Sal vs. E2/P4 Sal | - | - | 0.6557 | - | |
| | | | | | Vehicle K (10) vs. E2 K (10) | - | - | 0.9914 | - | |
| | | | | | Vehicle K (10) vs. E2/P4 K (10) | - | - | 0.5792 | - | |
| | | | | | Vehicle (2R,6R)-HNK (0.025) vs. E2 (2R,6R)-HNK | - | - | 0.0028 | ** | |
| | | | | | Vehicle (2R,6R)-HNK (0.025) vs. E2/P4 (2R,6R)- | - | - | 0.9995 | - | |
| | | | Immobility Time (min 3) | Tukey | Vehicle Sal vs. Vehicle K (10) | - | - | >0.9999 | - | |
| | | | | | Vehicle Sal vs. Vehicle (2R,6R)-HNK (0.025) | - | - | 0.7718 | - | |
| | | | | | E2 Sal vs. E2 K (10) | - | - | >0.9999 | - | 6E |
| | | | | | E2 Sal vs. E2 (2R,6R)-HNK (0.025) | - | - | 0.9982 | - | |
| | Forced Swim Test Day 1 | FST Day 1 | | | E2/P4 Sal vs. E2/P4 K (10) | - | - | >0.9999 | - | |
| | | | | | E2/P4 Sal vs. E2/P4 (2R,6R)-HNK (0.025) | - | - | 0.9995 | - | |
| | | | | | Vehicle Sal vs. E2 Sal | - | - | 0.5811 | - | |
| | | | | | Vehicle Sal vs. E2/P4 Sal | - | - | 0.9753 | - | |
| | | | | | Vehicle K (10) vs. E2 K (10) | - | - | 0.6047 | - | |
| | | | | | Vehicle K (10) vs. E2/P4 K (10) | - | - | 0.9478 | - | |

| | | | | | | | | | | |
|---|---|---|---|---|---|---|---|---|---|---|
| Ovariectomized, hormone replacement, 1 week prophylactic drug | | | | | Vehicle (2R,6R)-HNK (0.025) vs. E2 (2R,6R)-HNK | - | - | 0.7544 | - | |
| | | | | | Vehicle (2R,6R)-HNK (0.025) vs. E2/P4 (2R,6R)- | - | - | 0.4500 | - | |
| | | | Immobility Time (min 4) | Tukey | Vehicle Sal vs. Vehicle K (10) | - | - | 0.8928 | - | |
| | | | | | Vehicle Sal vs. Vehicle (2R,6R)-HNK (0.025) | - | - | >0.9999 | - | |
| | | | | | E2 Sal vs. E2 K (10) | - | - | 0.2274 | - | |
| | | | | | E2 Sal vs. E2 (2R,6R)-HNK (0.025) | - | - | 0.0077 | ** | |
| | | | | | E2/P4 Sal vs. E2/P4 K (10) | - | - | 0.9995 | - | |
| | | | | | E2/P4 Sal vs. E2/P4 (2R,6R)-HNK (0.025) | - | - | 0.9988 | - | |
| | | | | | Vehicle Sal vs. E2 Sal | - | - | >0.9999 | - | |
| | | | | | Vehicle Sal vs. E2/P4 Sal | - | - | 0.9725 | - | |
| | | | | | Vehicle K (10) vs. E2 K (10) | - | - | 0.5006 | - | |
| | | | | | Vehicle K (10) vs. E2/P4 K (10) | - | - | 0.7255 | - | |
| | | | | | Vehicle (2R,6R)-HNK (0.025) vs. E2 (2R,6R)-HNK | - | - | 0.0005 | ** | |
| | | | | | Vehicle (2R,6R)-HNK (0.025) vs. E2/P4 (2R,6R)- | - | - | 0.9992 | - | |
| | | | Immobility Time (min 5) | Tukey | Vehicle Sal vs. Vehicle K (10) | - | - | 0.9763 | - | |
| | | | | | Vehicle Sal vs. Vehicle (2R,6R)-HNK (0.025) | - | - | >0.9999 | - | |
| | | | | | E2 Sal vs. E2 K (10) | - | - | >0.9999 | - | |
| | | | | | E2 Sal vs. E2 (2R,6R)-HNK (0.025) | - | - | 0.1485 | - | |
| | | | | | E2/P4 Sal vs. E2/P4 K (10) | - | - | 0.9914 | - | |
| | | | | | E2/P4 Sal vs. E2/P4 (2R,6R)-HNK (0.025) | - | - | 0.9286 | - | |
| | | | | | Vehicle Sal vs. E2 Sal | - | - | 0.7706 | - | |
| | | | | | Vehicle Sal vs. E2/P4 Sal | - | - | 0.7935 | - | |
| | | | | | Vehicle K (10) vs. E2 K (10) | - | - | 0.9999 | - | |
| | | | | | Vehicle K (10) vs. E2/P4 K (10) | - | - | 0.7831 | - | |
| | | | | | Vehicle (2R,6R)-HNK (0.025) vs. E2 (2R,6R)-HNK | - | - | <0.0001 | *** | |
| | | | | | Vehicle (2R,6R)-HNK (0.025) vs. E2/P4 (2R,6R)- | - | - | >0.9999 | - | |
| | | | Immobility Time (min 6) | Tukey | Vehicle Sal vs. Vehicle K (10) | - | - | 0.9046 | - | |
| | | | | | Vehicle Sal vs. Vehicle (2R,6R)-HNK (0.025) | - | - | >0.9999 | - | |
| | | | | | E2 Sal vs. E2 K (10) | - | - | >0.9999 | - | |
| | | | | | E2 Sal vs. E2 (2R,6R)-HNK (0.025) | - | - | >0.9999 | - | |
| | | | | | E2/P4 Sal vs. E2/P4 K (10) | - | - | 0.9421 | - | |
| | | | | | E2/P4 Sal vs. E2/P4 (2R,6R)-HNK (0.025) | - | - | 0.8585 | - | |
| | | | | | Vehicle Sal vs. E2 Sal | - | - | 0.4649 | - | |
| | | | | | Vehicle Sal vs. E2/P4 Sal | - | - | 0.6493 | - | |
| | | | | | Vehicle K (10) vs. E2 K (10) | - | - | 0.9837 | - | |
| | | | | | Vehicle K (10) vs. E2/P4 K (10) | - | - | 0.7130 | - | |
| | | | | | Vehicle (2R,6R)-HNK (0.025) vs. E2 (2R,6R)-HNK | - | - | 0.2164 | - | |
| | | | | | Vehicle (2R,6R)-HNK (0.025) vs. E2/P4 (2R,6R)- | - | - | >0.9999 | - | |
| | | | Immobility Time Average (min 3-6) | ANOVA | Hormone | 19.450 | 2,84 | <0.0001 | *** | data not shown |
| | | | | | Drug | 2.152 | 2,84 | 0.1226 | - | |
| | | | | | Hormone x Drug | 0.727 | 4,84 | 0.5760 | - | |
| | | | Immobility Time (min 1-6) | RMANOVA | Hormone | 9.468 | 2,82 | 0.0002 | ** | |
| | | | | | Drug | 6.087 | 2,82 | 0.0034 | ** | |
| | | | | | Hormone x Drug | 3.535 | 4,82 | 0.0103 | * | |
| | | | | | Time | 34.035 | 5,410 | <0.0001 | *** | |
| | | | | | Hormone x Time | 1.497 | 10,410 | 0.0138 | * | |
| | | | | | Drug x Time | 1.160 | 10,410 | 0.3162 | - | |
| | | | | | Hormone x Drug x Time | 0.603 | 20,410 | 0.9113 | - | |
| | | | | | Vehicle Sal vs. Vehicle K (10) | - | - | >0.9999 | - | |
| | | | | | Vehicle Sal vs. Vehicle (2R,6R)-HNK (0.025) | - | - | >0.9999 | - | |
| | | | | | E2 Sal vs. E2 K (10) | - | - | >0.9999 | - | |
| | | | | | E2 Sal vs. E2 (2R,6R)-HNK (0.025) | - | - | 0.1366 | - | |
| | | | | | E2/P4 Sal vs. E2/P4 K (10) | - | - | 0.0771 | - | |
| | | | | | E2/P4 Sal vs. E2/P4 (2R,6R)-HNK (0.025) | - | - | 0.2023 | - | |

| | | | | | | | | | | |
|---|---|---|---|---|---|---|---|---|---|---|
| | Forced Swim Test Day 2 | FST Day 2 | | Tukey | Vehicle Sal vs. E2 Sal | - | - | >0.9999 | - | 6F |
| | | | | | Vehicle Sal vs. E2/P4 Sal | - | - | >0.9999 | - | |
| | | | | | Vehicle K (10) vs. E2 K (10) | - | - | >0.9999 | - | |
| | | | | | Vehicle K (10) vs. E2/P4 K (10) | - | - | 0.0422 | * | |
| | | | | | Vehicle (2*R*,6*R*)-HNK (0.025) vs. E2 (2*R*,6*R*)-HNK | - | - | 0.0082 | ** | |
| | | | | | Vehicle (2*R*,6*R*)-HNK (0.025) vs. E2/P4 (2*R*,6*R*)-HNK (0.025) | - | - | 0.0025 | ** | |
| | | | Immobility Time (min 1) | Tukey | Vehicle Sal vs. Vehicle K (10) | - | - | >0.9999 | - | |
| | | | | | Vehicle Sal vs. Vehicle (2*R*,6*R*)-HNK (0.025) | - | - | >0.9999 | - | |
| | | | | | E2 Sal vs. E2 K (10) | - | - | 0.9992 | - | |
| | | | | | E2 Sal vs. E2 (2*R*,6*R*)-HNK (0.025) | - | - | 0.0658 | - | |
| | | | | | E2/P4 Sal vs. E2/P4 K (10) | - | - | 0.9958 | - | |
| | | | | | E2/P4 Sal vs. E2/P4 (2*R*,6*R*)-HNK (0.025) | - | - | 0.9687 | - | |
| | | | | | Vehicle Sal vs. E2 Sal | - | - | 0.9325 | - | |
| | | | | | Vehicle Sal vs. E2/P4 Sal | - | - | 0.9966 | - | |
| | | | | | Vehicle K (10) vs. E2 K (10) | - | - | 0.9996 | - | |
| | | | | | Vehicle K (10) vs. E2/P4 K (10) | - | - | 0.8224 | - | |
| | | | | | Vehicle (2*R*,6*R*)-HNK (0.025) vs. E2 (2*R*,6*R*)-HNK | - | - | 0.2213 | - | |
| | | | | | Vehicle (2*R*,6*R*)-HNK (0.025) vs. E2/P4 (2*R*,6*R*)-HNK (0.025) | - | - | 0.2130 | - | |
| | | | Immobility Time (min 2) | Tukey | Vehicle Sal vs. Vehicle K (10) | - | - | 0.9961 | - | |
| | | | | | Vehicle Sal vs. Vehicle (2*R*,6*R*)-HNK (0.025) | - | - | 0.998 | - | |
| | | | | | E2 Sal vs. E2 K (10) | - | - | 0.922 | - | |
| | | | | | E2 Sal vs. E2 (2*R*,6*R*)-HNK (0.025) | - | - | 0.3509 | - | |
| | | | | | E2/P4 Sal vs. E2/P4 K (10) | - | - | 0.1450 | - | |
| | | | | | E2/P4 Sal vs. E2/P4 (2*R*,6*R*)-HNK (0.025) | - | - | 0.7033 | - | |
| | | | | | Vehicle Sal vs. E2 Sal | - | - | >0.9999 | - | |
| | | | | | Vehicle Sal vs. E2/P4 Sal | - | - | >0.9999 | - | |
| | | | | | Vehicle K (10) vs. E2 K (10) | - | - | 0.9943 | - | |
| | | | | | Vehicle K (10) vs. E2/P4 K (10) | - | - | 0.4152 | - | |
| | | | | | Vehicle (2*R*,6*R*)-HNK (0.025) vs. E2 (2*R*,6*R*)-HNK | - | - | 0.0508 | - | |
| | | | | | Vehicle (2*R*,6*R*)-HNK (0.025) vs. E2/P4 (2*R*,6*R*)-HNK (0.025) | - | - | 0.2076 | - | |
| | | | Immobility Time (min 3) | Tukey | Vehicle Sal vs. Vehicle K (10) | - | - | 0.9901 | - | |
| | | | | | Vehicle Sal vs. Vehicle (2*R*,6*R*)-HNK (0.025) | - | - | >0.9999 | - | |
| | | | | | E2 Sal vs. E2 K (10) | - | - | >0.9999 | - | |
| | | | | | E2 Sal vs. E2 (2*R*,6*R*)-HNK (0.025) | - | - | 0.6804 | - | |
| | | | | | E2/P4 Sal vs. E2/P4 K (10) | - | - | 0.6408 | - | |
| | | | | | E2/P4 Sal vs. E2/P4 (2*R*,6*R*)-HNK (0.025) | - | - | 0.8529 | - | |
| | | | | | Vehicle Sal vs. E2 Sal | - | - | 0.9324 | - | |
| | | | | | Vehicle Sal vs. E2/P4 Sal | - | - | >0.9999 | - | |
| | | | | | Vehicle K (10) vs. E2 K (10) | - | - | 0.9984 | - | |
| | | | | | Vehicle K (10) vs. E2/P4 K (10) | - | - | 0.6213 | - | |
| | | | | | Vehicle (2*R*,6*R*)-HNK (0.025) vs. E2 (2*R*,6*R*)-HNK | - | - | 0.0131 | * | |
| | | | | | Vehicle (2*R*,6*R*)-HNK (0.025) vs. E2/P4 (2*R*,6*R*)-HNK (0.025) | - | - | 0.2279 | - | |
| | | | Immobility Time (min 4) | Tukey | Vehicle Sal vs. Vehicle K (10) | - | - | 0.9991 | - | |
| | | | | | Vehicle Sal vs. Vehicle (2*R*,6*R*)-HNK (0.025) | - | - | >0.9999 | - | |
| | | | | | E2 Sal vs. E2 K (10) | - | - | 0.9695 | - | |
| | | | | | E2 Sal vs. E2 (2*R*,6*R*)-HNK (0.025) | - | - | 0.7858 | - | |
| | | | | | E2/P4 Sal vs. E2/P4 K (10) | - | - | 0.0373 | * | |
| | | | | | E2/P4 Sal vs. E2/P4 (2*R*,6*R*)-HNK (0.025) | - | - | 0.8027 | - | |
| | | | | | Vehicle Sal vs. E2 Sal | - | - | 0.9990 | - | |

| | | | | | | | | | | |
|---|---|---|---|---|---|---|---|---|---|---|
| | | | 4) | | Vehicle Sal vs. E2/P4 Sal | - | - | >0.9999 | - | |
| | | | | | Vehicle K (10) vs. E2 K (10) | - | - | >0.9999 | - | |
| | | | | | Vehicle K (10) vs. E2/P4 K (10) | - | - | 0.012 | * | |
| | | | | | Vehicle (2R,6R)-HNK (0.025) vs. E2 (2R,6R)-HNK | - | - | 0.9189 | - | |
| | | | | | Vehicle (2R,6R)-HNK (0.025) vs. E2/P4 (2R,6R)-HNK (0.025) | - | - | 0.3153 | - | |
| | | | Immobility Time (min 5) | Tukey | Vehicle Sal vs. Vehicle K (10) | - | - | >0.9999 | - | |
| | | | | | Vehicle Sal vs. Vehicle (2R,6R)-HNK (0.025) | - | - | 0.9128 | - | |
| | | | | | E2 Sal vs. E2 K (10) | - | - | >0.9999 | - | |
| | | | | | E2 Sal vs. E2 (2R,6R)-HNK (0.025) | - | - | 0.8508 | - | |
| | | | | | E2/P4 Sal vs. E2/P4 K (10) | - | - | 0.2821 | - | |
| | | | | | E2/P4 Sal vs. E2/P4 (2R,6R)-HNK (0.025) | - | - | 0.0794 | - | |
| | | | | | Vehicle Sal vs. E2 Sal | - | - | >0.9999 | - | |
| | | | | | Vehicle Sal vs. E2/P4 Sal | - | - | 0.9926 | - | |
| | | | | | Vehicle K (10) vs. E2 K (10) | - | - | 0.9996 | - | |
| | | | | | Vehicle K (10) vs. E2/P4 K (10) | - | - | 0.5970 | - | |
| | | | | | Vehicle (2R,6R)-HNK (0.025) vs. E2 (2R,6R)-HNK | - | - | 0.1024 | - | |
| | | | | | Vehicle (2R,6R)-HNK (0.025) vs. E2/P4 (2R,6R)-HNK (0.025) | - | - | 0.0035 | ** | |
| | | | Immobility Time (min 6) | Tukey | Vehicle Sal vs. Vehicle K (10) | - | - | >0.9999 | - | |
| | | | | | Vehicle Sal vs. Vehicle (2R,6R)-HNK (0.025) | - | - | >0.9999 | - | |
| | | | | | E2 Sal vs. E2 K (10) | - | - | >0.9999 | - | |
| | | | | | E2 Sal vs. E2 (2R,6R)-HNK (0.025) | - | - | 0.2923 | - | |
| | | | | | E2/P4 Sal vs. E2/P4 K (10) | - | - | 0.2463 | - | |
| | | | | | E2/P4 Sal vs. E2/P4 (2R,6R)-HNK (0.025) | - | - | 0.0618 | - | |
| | | | | | Vehicle Sal vs. E2 Sal | - | - | >0.9999 | - | |
| | | | | | Vehicle Sal vs. E2/P4 Sal | - | - | >0.9999 | - | |
| | | | | | Vehicle K (10) vs. E2 K (10) | - | - | 0.9993 | - | |
| | | | | | Vehicle K (10) vs. E2/P4 K (10) | - | - | 0.0356 | * | |
| | | | | | Vehicle (2R,6R)-HNK (0.025) vs. E2 (2R,6R)-HNK | - | - | 0.0493 | * | |
| | | | | | Vehicle (2R,6R)-HNK (0.025) vs. E2/P4 (2R,6R)-HNK (0.025) | - | - | 0.0017 | ** | |
| | | | Immobility Time Average (min 3-6) | ANOVA | Hormone | 10.270 | 2,85 | 0.0001 | ** | 6G |
| | | | | | Drug | 5.784 | 2,85 | 0.0044 | ** | |
| | | | | | Hormone x Drug | 3.317 | 4,85 | 0.0142 | * | |
| | | | | Sidak | Vehicle Sal vs. Vehicle K (10) | - | - | 0.8674 | - | |
| | | | | | Vehicle Sal vs. Vehicle (2R,6R)-HNK (0.025) | - | - | 0.8309 | - | |
| | | | | | E2 Sal vs. E2 K (10) | - | - | 0.7351 | - | |
| | | | | | E2 Sal vs. E2 (2R,6R)-HNK (0.025) | - | - | 0.0300 | * | |
| | | | | | E2/P4 Sal vs. E2/P4 K (10) | - | - | 0.0023 | ** | |
| | | | | | E2/P4 Sal vs. E2/P4 (2R,6R)-HNK (0.025) | - | - | 0.0056 | ** | |
| | | | | | Vehicle Sal vs. E2 Sal | - | - | 0.9552 | - | |
| | | | | | Vehicle Sal vs. E2/P4 Sal | - | - | 0.9994 | - | |
| | | | | | Vehicle K (10) vs. E2 K (10) | - | - | 0.6898 | - | |
| | | | | | Vehicle K (10) vs. E2/P4 K (10) | - | - | 0.0011 | ** | |
| | | | | | Vehicle (2R,6R)-HNK (0.025) vs. E2 (2R,6R)-HNK | - | - | 0.0008 | ** | |
| | | | | | Vehicle (2R,6R)-HNK (0.025) vs. E2/P4 (2R,6R)-HNK (0.025) | - | - | <0.0001 | *** | |
| | | | Freezing (min 1-5) | RMANOVA | Drug | 1.096 | 1,22 | 0.3066 | - | S01A |
| | | | | | Time | 39.95 | 4,88 | <0.0001 | *** | |
| | | | | | Drug x Time | 1.239 | 4,88 | 0.3001 | - | |
| | | | | | Drug | 1.846 | 7,72 | 0.0915 | - | |

| | | | | | | | | | | |
|---|---|---|---|---|---|---|---|---|---|---|
| | | | Freezing (min 1-5) | RMANOVA | Time | 222 | 4,288 | <0.0001 | *** | S01B |
| | | | | | Drug x Time | 1.493 | 28,288 | 0.0565 | - | |
| | | | Freezing (min 1-5) | RMANOVA | Drug | 3.046 | 7,72 | 0.0073 | ** | S01C |
| | | | | | Time | 144.1 | 4,288 | <0.0001 | *** | |
| | | | | | Drug x Time | 2.185 | 28,288 | 0.0007 | ** | |
| | | | | Dunnett | Sal vs. (2*S*,6*S*)-HNK (0.025) | - | - | 0.0093 | ** | |
| | | | | | Sal vs. (2*S*,6*S*)-HNK (0.075) | - | - | 0.4587 | - | |
| | | | | | Sal vs. (2*S*,6*S*)-HNK (0.1) | - | - | 0.0033 | ** | |
| | | | | | Sal vs. (2*S*,6*S*)-HNK (0.3) | - | - | 0.0463 | * | |
| | | | | | Sal vs. (2*S*,6*S*)-HNK (2.5) | - | - | 0.1658 | - | |
| | | | | | Sal vs. (2*S*,6*S*)-HNK (10) | - | - | 0.0066 | ** | |
| | | | | | Sal vs. (2*S*,6*S*)-HNK (30) | - | - | 0.0473 | * | |
| | | | Freezing (min 1) | Dunnett | Sal vs. (2*S*,6*S*)-HNK (0.025) | - | - | >0.9999 | - | |
| | | | | | Sal vs. (2*S*,6*S*)-HNK (0.075) | - | - | >0.9999 | - | |
| | | | | | Sal vs. (2*S*,6*S*)-HNK (0.1) | - | - | >0.9999 | - | |
| | | | | | Sal vs. (2*S*,6*S*)-HNK (0.3) | - | - | >0.9999 | - | |
| | | | | | Sal vs. (2*S*,6*S*)-HNK (2.5) | - | - | >0.9999 | - | |
| | | | | | Sal vs. (2*S*,6*S*)-HNK (10) | - | - | >0.9999 | - | |
| | | | | | Sal vs. (2*S*,6*S*)-HNK (30) | - | - | >0.9999 | - | |
| | Contextual Fear Conditioning Training | CFC Training | Freezing (min 2) | Dunnett | Sal vs. (2*S*,6*S*)-HNK (0.025) | - | - | >0.9999 | - | |
| | | | | | Sal vs. (2*S*,6*S*)-HNK (0.075) | - | - | >0.9999 | - | |
| | | | | | Sal vs. (2*S*,6*S*)-HNK (0.1) | - | - | 0.9996 | - | |
| | | | | | Sal vs. (2*S*,6*S*)-HNK (0.3) | - | - | >0.9999 | - | |
| | | | | | Sal vs. (2*S*,6*S*)-HNK (2.5) | - | - | >0.9999 | - | |
| | | | | | Sal vs. (2*S*,6*S*)-HNK (10) | - | - | >0.9999 | - | |
| | | | | | Sal vs. (2*S*,6*S*)-HNK (30) | - | - | >0.9999 | - | |
| | | | Freezing (min 3) | Dunnett | Sal vs. (2*S*,6*S*)-HNK (0.025) | - | - | >0.9999 | - | |
| | | | | | Sal vs. (2*S*,6*S*)-HNK (0.075) | - | - | 0.9997 | - | |
| | | | | | Sal vs. (2*S*,6*S*)-HNK (0.1) | - | - | 0.9972 | - | |
| | | | | | Sal vs. (2*S*,6*S*)-HNK (0.3) | - | - | 0.9997 | - | |
| | | | | | Sal vs. (2*S*,6*S*)-HNK (2.5) | - | - | 0.9999 | - | |
| | | | | | Sal vs. (2*S*,6*S*)-HNK (10) | - | - | 0.9997 | - | |
| | | | | | Sal vs. (2*S*,6*S*)-HNK (30) | - | - | 0.9997 | - | |
| | | | Freezing (min 4) | Dunnett | Sal vs. (2*S*,6*S*)-HNK (0.025) | - | - | 0.0003 | ** | |
| | | | | | Sal vs. (2*S*,6*S*)-HNK (0.075) | - | - | 0.0468 | * | |
| | | | | | Sal vs. (2*S*,6*S*)-HNK (0.1) | - | - | <0.0001 | *** | |
| | | | | | Sal vs. (2*S*,6*S*)-HNK (0.3) | - | - | 0.0176 | * | |
| | | | | | Sal vs. (2*S*,6*S*)-HNK (2.5) | - | - | 0.2049 | - | |
| | | | | | Sal vs. (2*S*,6*S*)-HNK (10) | - | - | 0.0002 | ** | |
| | | | | | Sal vs. (2*S*,6*S*)-HNK (30) | - | - | 0.001 | ** | |
| | | | Freezing (min 5) | Dunnett | Sal vs. (2*S*,6*S*)-HNK (0.025) | - | - | <0.0001 | *** | |
| | | | | | Sal vs. (2*S*,6*S*)-HNK (0.075) | - | - | 0.6548 | - | |
| | | | | | Sal vs. (2*S*,6*S*)-HNK (0.1) | - | - | <0.0001 | *** | |
| | | | | | Sal vs. (2*S*,6*S*)-HNK (0.3) | - | - | 0.0007 | ** | |
| | | | | | Sal vs. (2*S*,6*S*)-HNK (2.5) | - | - | 0.0026 | ** | |
| | | | | | Sal vs. (2*S*,6*S*)-HNK (10) | - | - | <0.0001 | *** | |
| | | | | | Sal vs. (2*S*,6*S*)-HNK (30) | | | 0.0147 | * | |
| | | | Freezing (min 1-5) | RMANOVA | Drug | 2.052 | 1,22 | 0.1661 | - | |
| | | | | | Time | 14.23 | 4,88 | <0.0001 | *** | |
| | | | | | Drug x Time | 0.0471 | 4,88 | 0.9957 | - | |
| | | | Freezing (min 1-5) | RMANOVA | Drug | 0.6765 | 7,72 | 0.6913 | - | |
| | | | | | Time | 48.16 | 4,288 | <0.0001 | *** | |
| | | | | | Drug x Time | 1.944 | 28,288 | 0.0038 | ** | |
| | | | | | Sal vs. (2*R*,6*R*)-HNK (0.025) | - | - | >0.9999 | - | |

| | | | | | | | | | | |
|---|---|---|---|---|---|---|---|---|---|---|
| | | | Freezing (min 1) | Dunnett | Sal vs. (2R,6R)-HNK (0.075) | - | - | 0.2625 | - | |
| | | | | | Sal vs. (2R,6R)-HNK (0.1) | - | - | 0.3965 | - | |
| | | | | | Sal vs. (2R,6R)-HNK (0.3) | - | - | 0.2186 | - | |
| | | | | | Sal vs. (2R,6R)-HNK (2.5) | - | - | 0.0044 | ** | |
| | | | | | Sal vs. (2R,6R)-HNK (10) | - | - | 0.15 | - | |
| | | | | | Sal vs. (2R,6R)-HNK (30) | - | - | 0.9819 | - | |
| | | | Freezing (min 2) | Dunnett | Sal vs. (2R,6R)-HNK (0.025) | - | - | 0.94 | - | |
| | | | | | Sal vs. (2R,6R)-HNK (0.075) | - | - | 0.5234 | - | |
| | | | | | Sal vs. (2R,6R)-HNK (0.1) | - | - | 0.7885 | - | |
| | | | | | Sal vs. (2R,6R)-HNK (0.3) | - | - | 0.4929 | - | |
| | | | | | Sal vs. (2R,6R)-HNK (2.5) | - | - | 0.723 | - | |
| | | | | | Sal vs. (2R,6R)-HNK (10) | - | - | 0.782 | - | |
| | | | | | Sal vs. (2R,6R)-HNK (30) | - | - | 0.584 | - | |
| | | | Freezing (min 3) | Dunnett | Sal vs. (2R,6R)-HNK (0.025) | - | - | 0.9207 | - | |
| | | | | | Sal vs. (2R,6R)-HNK (0.075) | - | - | 0.4346 | - | |
| | | | | | Sal vs. (2R,6R)-HNK (0.1) | - | - | 0.945 | - | |
| | | | | | Sal vs. (2R,6R)-HNK (0.3) | - | - | 0.481 | - | |
| | | | | | Sal vs. (2R,6R)-HNK (2.5) | - | - | 0.3394 | - | |
| | | | | | Sal vs. (2R,6R)-HNK (10) | - | - | 0.8739 | - | |
| | | | | | Sal vs. (2R,6R)-HNK (30) | - | - | 0.8846 | - | |
| | | | Freezing (min 4) | Dunnett | Sal vs. (2R,6R)-HNK (0.025) | - | - | 0.9995 | - | |
| | | | | | Sal vs. (2R,6R)-HNK (0.075) | - | - | 0.9946 | - | |
| | | | | | Sal vs. (2R,6R)-HNK (0.1) | - | - | 0.9996 | - | |
| | | | | | Sal vs. (2R,6R)-HNK (0.3) | - | - | 0.9922 | - | |
| | | | | | Sal vs. (2R,6R)-HNK (2.5) | - | - | 0.995 | - | |
| | | | | | Sal vs. (2R,6R)-HNK (10) | - | - | 0.9999 | - | |
| | | | | | Sal vs. (2R,6R)-HNK (30) | - | - | 0.8891 | - | |
| | | | Freezing (min 5) | Dunnett | Sal vs. (2R,6R)-HNK (0.025) | - | - | >0.9999 | - | |
| | | | | | Sal vs. (2R,6R)-HNK (0.075) | - | - | 0.9912 | - | |
| | | | | | Sal vs. (2R,6R)-HNK (0.1) | - | - | 0.5591 | - | |
| | | | | | Sal vs. (2R,6R)-HNK (0.3) | - | - | 0.8736 | - | |
| | | | | | Sal vs. (2R,6R)-HNK (2.5) | - | - | 0.9719 | - | |
| | | | | | Sal vs. (2R,6R)-HNK (10) | - | - | 0.9995 | - | |
| | | | | | Sal vs. (2R,6R)-HNK (30) | - | - | 0.5605 | - | |
| | Contextual Fear Conditioning Re-exposure | CFC Re-exposure | Freezing (min 1-5) | RMANOVA | Drug | 5.66 | 7,72 | <0.0001 | *** | |
| | | | | | Time | 51.64 | 4,288 | <0.0001 | *** | data not shown |
| | | | | | Drug x Time | 1.821 | 28,288 | 0.0083 | ** | |
| | | | | Dunnett | Sal vs. (2S,6S)-HNK (0.025) | - | - | 0.0016 | ** | |
| | | | | | Sal vs. (2S,6S)-HNK (0.075) | - | - | 0.0129 | * | |
| | | | | | Sal vs. (2S,6S)-HNK (0.1) | - | - | <0.0001 | *** | |
| | | | | | Sal vs. (2S,6S)-HNK (0.3) | - | - | <0.0001 | *** | |
| | | | | | Sal vs. (2S,6S)-HNK (2.5) | - | - | 0.1475 | - | |
| | | | | | Sal vs. (2S,6S)-HNK (10) | - | - | 0.0486 | * | |
| | | | | | Sal vs. (2S,6S)-HNK (30) | - | - | 0.012 | * | |
| | | | Freezing (min 1) | Dunnett | Sal vs. (2S,6S)-HNK (0.025) | - | - | <0.0001 | *** | |
| | | | | | Sal vs. (2S,6S)-HNK (0.075) | - | - | 0.0002 | ** | |
| | | | | | Sal vs. (2S,6S)-HNK (0.1) | - | - | 0.0004 | ** | |
| | | | | | Sal vs. (2S,6S)-HNK (0.3) | - | - | 0.0003 | ** | |
| | | | | | Sal vs. (2S,6S)-HNK (2.5) | - | - | 0.006 | ** | |
| | | | | | Sal vs. (2S,6S)-HNK (10) | - | - | 0.0005 | ** | |
| | | | | | Sal vs. (2S,6S)-HNK (30) | - | - | 0.0003 | ** | |
| | | | | | Sal vs. (2S,6S)-HNK (0.025) | - | - | 0.0006 | ** | |
| | | | | | Sal vs. (2S,6S)-HNK (0.075) | - | - | 0.0152 | * | |
| | | | | | Sal vs. (2S,6S)-HNK (0.1) | - | - | <0.0001 | *** | |

| | | | | | | | | | | |
|---|---|---|---|---|---|---|---|---|---|---|
| | | | Freezing (min 2) | Dunnett | Sal vs. (2*S*,6*S*)-HNK (0.3) | - | - | 0.0003 | ** | |
| | | | | | Sal vs. (2*S*,6*S*)-HNK (2.5) | - | - | 0.3308 | - | |
| | | | | | Sal vs. (2*S*,6*S*)-HNK (10) | - | - | 0.0095 | ** | |
| | | | | | Sal vs. (2*S*,6*S*)-HNK (30) | - | - | 0.0489 | * | |
| | | | Freezing (min 3) | Dunnett | Sal vs. (2*S*,6*S*)-HNK (0.025) | - | - | 0.0151 | * | |
| | | | | | Sal vs. (2*S*,6*S*)-HNK (0.075) | - | - | 0.2889 | - | |
| | | | | | Sal vs. (2*S*,6*S*)-HNK (0.1) | - | - | 0.0001 | ** | |
| | | | | | Sal vs. (2*S*,6*S*)-HNK (0.3) | - | - | 0.0001 | ** | |
| | | | | | Sal vs. (2*S*,6*S*)-HNK (2.5) | - | - | 0.2169 | - | |
| | | | | | Sal vs. (2*S*,6*S*)-HNK (10) | - | - | 0.2648 | - | |
| | | | | | Sal vs. (2*S*,6*S*)-HNK (30) | - | - | 0.0845 | - | |
| | | | Freezing (min 4) | Dunnett | Sal vs. (2*S*,6*S*)-HNK (0.025) | - | - | 0.0990 | - | |
| | | | | | Sal vs. (2*S*,6*S*)-HNK (0.075) | - | - | 0.3296 | - | |
| | | | | | Sal vs. (2*S*,6*S*)-HNK (0.1) | - | - | 0.0016 | ** | |
| | | | | | Sal vs. (2*S*,6*S*)-HNK (0.3) | - | - | <0.0001 | *** | |
| | | | | | Sal vs. (2*S*,6*S*)-HNK (2.5) | - | - | 0.9702 | - | |
| | | | | | Sal vs. (2*S*,6*S*)-HNK (10) | - | - | 0.9994 | - | |
| | | | | | Sal vs. (2*S*,6*S*)-HNK (30) | - | - | 0.2042 | - | |
| | | | Freezing (min 5) | Dunnett | Sal vs. (2*S*,6*S*)-HNK (0.025) | - | - | 0.8935 | - | |
| | | | | | Sal vs. (2*S*,6*S*)-HNK (0.075) | - | - | 0.6411 | - | |
| 1 week prophylactic drug, male | | | | | Sal vs. (2*S*,6*S*)-HNK (0.1) | - | - | 0.0032 | ** | |
| | | | | | Sal vs. (2*S*,6*S*)-HNK (0.3) | - | - | 0.1629 | - | |
| | | | | | Sal vs. (2*S*,6*S*)-HNK (2.5) | - | - | 0.9704 | - | |
| | | | | | Sal vs. (2*S*,6*S*)-HNK (10) | - | - | 0.9026 | - | |
| | | | | | Sal vs. (2*S*,6*S*)-HNK (30) | - | - | 0.7476 | - | |
| | | | Immobility Time (min 1-6) | RMANOVA | Drug | 0.02342 | 1,22 | 0.8798 | - | |
| | | | | | Time | 20.07 | 5,110 | <0.0001 | *** | |
| | | | | | Drug x Time | 3.92 | 5,110 | 0.0026 | ** | |
| | | | Immobility Time (min 1) | Sidak | Sal vs. K (30) | - | - | 0.0743 | - | |
| | | | Immobility Time (min 2) | Sidak | Sal vs. K (30) | - | - | >0.9999 | - | |
| | | | Immobility Time (min 3) | Sidak | Sal vs. K (30) | - | - | >0.9999 | - | S01D |
| | | | Immobility Time (min 4) | Sidak | Sal vs. K (30) | - | - | 0.5600 | - | |
| | | | Immobility Time (min 5) | Sidak | Sal vs. K (30) | - | - | >0.9999 | - | |
| | | | Immobility Time (min 6) | Sidak | Sal vs. K (30) | - | - | >0.9999 | - | |
| | | | Immobility Time (min 1-6) | RMANOVA | Drug | 2.211 | 7,72 | 0.0429 | * | |
| | | | | | Time | 59.59 | 5,365 | <0.0001 | *** | |
| | | | | | Drug x Time | 2.002 | 35,365 | 0.0009 | ** | |
| | | | | Dunnett | Sal vs. (2*R*,6*R*)-HNK (0.025) | - | - | 0.7506 | - | |
| | | | | | Sal vs. (2*R*,6*R*)-HNK (0.075) | - | - | 0.6504 | - | |
| | | | | | Sal vs. (2*R*,6*R*)-HNK (0.1) | - | - | 0.9941 | - | |
| | | | | | Sal vs. (2*R*,6*R*)-HNK (0.3) | - | - | 0.9891 | - | |
| | | | | | Sal vs. (2*R*,6*R*)-HNK (2.5) | - | - | 0.9997 | - | |
| | | | | | Sal vs. (2*R*,6*R*)-HNK (10) | - | - | 0.0111 | * | |
| | | | | | Sal vs. (2*R*,6*R*)-HNK (30) | - | - | 0.9874 | - | |
| | | | Immobility Time (min | | Sal vs. (2*R*,6*R*)-HNK (0.025) | - | - | 0.9668 | - | |
| | | | | | Sal vs. (2*R*,6*R*)-HNK (0.075) | - | - | 0.0345 | * | |
| | | | | | Sal vs. (2*R*,6*R*)-HNK (0.1) | - | - | 0.8090 | - | |

| | | | | | | | | | | |
|---|---|---|---|---|---|---|---|---|---|---|
| | Forced Swim Test Day 1 | FST Day 1 | Immobility Time (min 1) | Dunnett | Sal vs. (2*R*,6*R*)-HNK (0.3) | - | - | 0.9976 | - | S01E |
| | | | | | Sal vs. (2*R*,6*R*)-HNK (2.5) | - | - | 0.7894 | - | |
| | | | | | Sal vs. (2*R*,6*R*)-HNK (10) | - | - | 0.0123 | * | |
| | | | | | Sal vs. (2*R*,6*R*)-HNK (30) | - | - | 0.9996 | - | |
| | | | Immobility Time (min 2) | Dunnett | Sal vs. (2*R*,6*R*)-HNK (0.025) | - | - | 0.9998 | - | |
| | | | | | Sal vs. (2*R*,6*R*)-HNK (0.075) | - | - | 0.9996 | - | |
| | | | | | Sal vs. (2*R*,6*R*)-HNK (0.1) | - | - | 0.9997 | - | |
| | | | | | Sal vs. (2*R*,6*R*)-HNK (0.3) | - | - | 0.9998 | - | |
| | | | | | Sal vs. (2*R*,6*R*)-HNK (2.5) | - | - | 0.9995 | - | |
| | | | | | Sal vs. (2*R*,6*R*)-HNK (10) | - | - | 0.0084 | ** | |
| | | | | | Sal vs. (2*R*,6*R*)-HNK (30) | - | - | 0.7386 | - | |
| | | | Immobility Time (min 3) | Dunnett | Sal vs. (2*R*,6*R*)-HNK (0.025) | - | - | 0.9863 | - | |
| | | | | | Sal vs. (2*R*,6*R*)-HNK (0.075) | - | - | 0.9971 | - | |
| | | | | | Sal vs. (2*R*,6*R*)-HNK (0.1) | - | - | 0.9933 | - | |
| | | | | | Sal vs. (2*R*,6*R*)-HNK (0.3) | - | - | 0.9103 | - | |
| | | | | | Sal vs. (2*R*,6*R*)-HNK (2.5) | - | - | 0.8887 | - | |
| | | | | | Sal vs. (2*R*,6*R*)-HNK (10) | - | - | 0.1164 | - | |
| | | | | | Sal vs. (2*R*,6*R*)-HNK (30) | - | - | 0.9994 | - | |
| | | | Immobility Time (min 4) | Dunnett | Sal vs. (2*R*,6*R*)-HNK (0.025) | - | - | 0.2657 | - | |
| | | | | | Sal vs. (2*R*,6*R*)-HNK (0.075) | - | - | 0.5759 | - | |
| | | | | | Sal vs. (2*R*,6*R*)-HNK (0.1) | - | - | 0.7809 | - | |
| | | | | | Sal vs. (2*R*,6*R*)-HNK (0.3) | - | - | 0.4524 | - | |
| | | | | | Sal vs. (2*R*,6*R*)-HNK (2.5) | - | - | 0.9941 | - | |
| | | | | | Sal vs. (2*R*,6*R*)-HNK (10) | - | - | 0.2398 | - | |
| | | | | | Sal vs. (2*R*,6*R*)-HNK (30) | - | - | 0.9995 | - | |
| | | | Immobility Time (min 5) | Dunnett | Sal vs. (2*R*,6*R*)-HNK (0.025) | - | - | 0.7169 | - | |
| | | | | | Sal vs. (2*R*,6*R*)-HNK (0.075) | - | - | 0.9999 | - | |
| | | | | | Sal vs. (2*R*,6*R*)-HNK (0.1) | - | - | 0.6508 | - | |
| | | | | | Sal vs. (2*R*,6*R*)-HNK (0.3) | - | - | 0.9732 | - | |
| | | | | | Sal vs. (2*R*,6*R*)-HNK (2.5) | - | - | 0.9868 | - | |
| | | | | | Sal vs. (2*R*,6*R*)-HNK (10) | - | - | 0.9895 | - | |
| | | | | | Sal vs. (2*R*,6*R*)-HNK (30) | - | - | 0.3435 | - | |
| | | | Immobility Time (min 6) | Dunnett | Sal vs. (2*R*,6*R*)-HNK (0.025) | - | - | 0.622 | - | |
| | | | | | Sal vs. (2*R*,6*R*)-HNK (0.075) | - | - | 0.9998 | - | |
| | | | | | Sal vs. (2*R*,6*R*)-HNK (0.1) | - | - | 0.9997 | - | |
| | | | | | Sal vs. (2*R*,6*R*)-HNK (0.3) | - | - | 0.9971 | - | |
| | | | | | Sal vs. (2*R*,6*R*)-HNK (2.5) | - | - | 0.8678 | - | |
| | | | | | Sal vs. (2*R*,6*R*)-HNK (10) | - | - | 0.3872 | - | |
| | | | | | Sal vs. (2*R*,6*R*)-HNK (30) | - | - | 0.7246 | - | |
| | | | Immobility Time (min 1-6) | RMANOVA | Drug | 5.981 | 7,72 | <0.0001 | *** | S01F |
| | | | | | Time | 121.7 | 5,365 | <0.0001 | *** | |
| | | | | | Drug x Time | 1.437 | 35,365 | 0.0559 | - | |
| | | | | Dunnett | Sal vs. (2*S*,6*S*)-HNK (0.025) | - | - | 0.1158 | - | |
| | | | | | Sal vs. (2*S*,6*S*)-HNK (0.075) | - | - | <0.0001 | *** | |
| | | | | | Sal vs. (2*S*,6*S*)-HNK (0.1) | - | - | <0.0001 | *** | |
| | | | | | Sal vs. (2*S*,6*S*)-HNK (0.3) | - | - | <0.0001 | *** | |
| | | | | | Sal vs. (2*S*,6*S*)-HNK (2.5) | - | - | 0.1281 | - | |
| | | | | | Sal vs. (2*S*,6*S*)-HNK (10) | - | - | 0.0112 | * | |
| | | | | | Sal vs. (2*S*,6*S*)-HNK (30) | - | - | 0.0003 | ** | |
| | | | Immobility Time (min 1-6) | RMANOVA | Drug | 31.12 | 1,22 | <0.0001 | *** | |
| | | | | | Time | 5.341 | 5,110 | 0.0002 | ** | |
| | | | | | Drug x Time | 0.2604 | 5,110 | 0.9337 | - | |
| | | | | *t*-test | Sal vs. K (30) | - | - | <0.0001 | *** | |
| | | | Immobility Time (min | | Drug | 1.316 | 7,72 | 0.2554 | - | |

| | | | | | | | | | | |
|---|---|---|---|---|---|---|---|---|---|---|
| | Forced Swim Test Day 2 | FST Day 2 | Immobility Time (min 1-6) | RMANOVA | Time | 20.09 | 5,360 | <0.0001 | *** | data not shown |
| | | | | | Drug x Time | 2.101 | 35,360 | 0.0004 | ** | |
| | | | Immobility Time (min 1) | Dunnett | Sal vs. (2*R*,6*R*)-HNK (0.025) | - | - | 0.0325 | * | |
| | | | | | Sal vs. (2*R*,6*R*)-HNK (0.075) | - | - | 0.8373 | - | |
| | | | | | Sal vs. (2*R*,6*R*)-HNK (0.1) | - | - | 0.9996 | - | |
| | | | | | Sal vs. (2*R*,6*R*)-HNK (0.3) | - | - | 0.9997 | - | |
| | | | | | Sal vs. (2*R*,6*R*)-HNK (2.5) | - | - | 0.3965 | - | |
| | | | | | Sal vs. (2*R*,6*R*)-HNK (10) | - | - | 0.6903 | - | |
| | | | | | Sal vs. (2*R*,6*R*)-HNK (30) | - | - | 0.9896 | - | |
| | | | Immobility Time (min 2) | Dunnett | Sal vs. (2*R*,6*R*)-HNK (0.025) | - | - | 0.9372 | - | |
| | | | | | Sal vs. (2*R*,6*R*)-HNK (0.075) | - | - | 0.9596 | - | |
| | | | | | Sal vs. (2*R*,6*R*)-HNK (0.1) | - | - | 0.7724 | - | |
| | | | | | Sal vs. (2*R*,6*R*)-HNK (0.3) | - | - | 0.9908 | - | |
| | | | | | Sal vs. (2*R*,6*R*)-HNK (2.5) | - | - | 0.3738 | - | |
| | | | | | Sal vs. (2*R*,6*R*)-HNK (10) | - | - | 0.9968 | - | |
| | | | | | Sal vs. (2*R*,6*R*)-HNK (30) | - | - | 0.9995 | - | |
| | | | Immobility Time (min 3) | Dunnett | Sal vs. (2*R*,6*R*)-HNK (0.025) | - | - | 0.9999 | - | |
| | | | | | Sal vs. (2*R*,6*R*)-HNK (0.075) | - | - | 0.1151 | - | |
| | | | | | Sal vs. (2*R*,6*R*)-HNK (0.1) | - | - | 0.7064 | - | |
| | | | | | Sal vs. (2*R*,6*R*)-HNK (0.3) | - | - | 0.9995 | - | |
| | | | | | Sal vs. (2*R*,6*R*)-HNK (2.5) | - | - | 0.4141 | - | |
| | | | | | Sal vs. (2*R*,6*R*)-HNK (10) | - | - | 0.9997 | - | |
| | | | | | Sal vs. (2*R*,6*R*)-HNK (30) | - | - | 0.9996 | - | |
| | | | Immobility Time (min 4) | Dunnett | Sal vs. (2*R*,6*R*)-HNK (0.025) | - | - | 0.9943 | - | |
| | | | | | Sal vs. (2*R*,6*R*)-HNK (0.075) | - | - | 0.9957 | - | |
| | | | | | Sal vs. (2*R*,6*R*)-HNK (0.1) | - | - | 0.5356 | - | |
| | | | | | Sal vs. (2*R*,6*R*)-HNK (0.3) | - | - | 0.9152 | - | |
| | | | | | Sal vs. (2*R*,6*R*)-HNK (2.5) | - | - | 0.9493 | - | |
| | | | | | Sal vs. (2*R*,6*R*)-HNK (10) | - | - | 0.8793 | - | |
| | | | | | Sal vs. (2*R*,6*R*)-HNK (30) | - | - | 0.8575 | - | |
| | | | Immobility Time (min 5) | Dunnett | Sal vs. (2*R*,6*R*)-HNK (0.025) | - | - | 0.9844 | - | |
| | | | | | Sal vs. (2*R*,6*R*)-HNK (0.075) | - | - | 0.1978 | - | |
| | | | | | Sal vs. (2*R*,6*R*)-HNK (0.1) | - | - | 0.8044 | - | |
| | | | | | Sal vs. (2*R*,6*R*)-HNK (0.3) | - | - | 0.8774 | - | |
| | | | | | Sal vs. (2*R*,6*R*)-HNK (2.5) | - | - | 0.9996 | - | |
| | | | | | Sal vs. (2*R*,6*R*)-HNK (10) | - | - | 0.9997 | - | |
| | | | | | Sal vs. (2*R*,6*R*)-HNK (30) | - | - | 0.798 | - | |
| | | | Immobility Time (min 6) | Dunnett | Sal vs. (2*R*,6*R*)-HNK (0.025) | - | - | 0.1771 | - | |
| | | | | | Sal vs. (2*R*,6*R*)-HNK (0.075) | - | - | 0.3349 | - | |
| | | | | | Sal vs. (2*R*,6*R*)-HNK (0.1) | - | - | 0.848 | - | |
| | | | | | Sal vs. (2*R*,6*R*)-HNK (0.3) | - | - | 0.1618 | - | |
| | | | | | Sal vs. (2*R*,6*R*)-HNK (2.5) | - | - | 0.9974 | - | |
| | | | | | Sal vs. (2*R*,6*R*)-HNK (10) | - | - | >0.9999 | - | |
| | | | | | Sal vs. (2*R*,6*R*)-HNK (30) | - | - | 0.9998 | - | |
| | | | Immobility Time (min 1-6) | RMANOVA | Drug | 0.8367 | 7,72 | 0.5311 | - | |
| | | | | | Time | 38.9 | 5,360 | <0.0001 | *** | |
| | | | | | Drug x Time | 1.137 | 35,360 | 0.2789 | - | |
| | | | Freezing (min 1-5) | RMANOVA | Drug | 0.847 | 3,69 | 0.4729 | - | S02A |
| | | | | | Time | 174.800 | 4,276 | <0.0001 | *** | |
| | | | | | Drug x Time | 1.126 | 12,276 | 0.3384 | - | |
| | | | | RMANOVA | Drug | 2.451 | 6,67 | 0.0334 | * | |
| | | | | | Time | 166.500 | 4,268 | <0.0001 | *** | |
| | | | | | Drug x Time | 1.784 | 24,268 | 0.0154 | * | |

| | | | | | | | | | | |
|---|---|---|---|---|---|---|---|---|---|---|
| | Contextual Fear Conditioning Training | CFC Training | Freezing (min 1-5) | Dunnett | Sal vs. (2*R*,6*R*)-HNK (0.025) | - | - | 0.5300 | - | S02B |
| | | | | | Sal vs. (2*R*,6*R*)-HNK (0.075) | - | - | 0.9327 | - | |
| | | | | | Sal vs. (2*R*,6*R*)-HNK (0.1) | - | - | 0.4764 | - | |
| | | | | | Sal vs. (2*R*,6*R*)-HNK (0.3) | - | - | 0.9880 | - | |
| | | | | | Sal vs. (2*R*,6*R*)-HNK (2.5) | - | - | 0.9935 | - | |
| | | | | | Sal vs. (2*R*,6*R*)-HNK (10) | - | - | 0.1229 | - | |
| | | | Freezing (min 1) | Dunnett | Sal vs. (2*R*,6*R*)-HNK (0.025) | - | - | 0.9998 | - | |
| | | | | | Sal vs. (2*R*,6*R*)-HNK (0.075) | - | - | >0.9999 | - | |
| | | | | | Sal vs. (2*R*,6*R*)-HNK (0.1) | - | - | 0.9998 | - | |
| | | | | | Sal vs. (2*R*,6*R*)-HNK (0.3) | - | - | >0.9999 | - | |
| | | | | | Sal vs. (2*R*,6*R*)-HNK (2.5) | - | - | 0.9998 | - | |
| | | | | | Sal vs. (2*R*,6*R*)-HNK (10) | - | - | >0.9999 | - | |
| | | | Freezing (min 2) | Dunnett | Sal vs. (2*R*,6*R*)-HNK (0.025) | - | - | 0.9997 | - | |
| | | | | | Sal vs. (2*R*,6*R*)-HNK (0.075) | - | - | >0.9999 | - | |
| | | | | | Sal vs. (2*R*,6*R*)-HNK (0.1) | - | - | 0.9997 | - | |
| | | | | | Sal vs. (2*R*,6*R*)-HNK (0.3) | - | - | 0.9999 | - | |
| | | | | | Sal vs. (2*R*,6*R*)-HNK (2.5) | - | - | >0.9999 | - | |
| | | | | | Sal vs. (2*R*,6*R*)-HNK (10) | - | - | 0.9999 | - | |
| | | | Freezing (min 3) | Dunnett | Sal vs. (2*R*,6*R*)-HNK (0.025) | - | - | 0.9928 | - | |
| | | | | | Sal vs. (2*R*,6*R*)-HNK (0.075) | - | - | 0.9997 | - | |
| | | | | | Sal vs. (2*R*,6*R*)-HNK (0.1) | - | - | >0.9999 | - | |
| | | | | | Sal vs. (2*R*,6*R*)-HNK (0.3) | - | - | >0.9999 | - | |
| | | | | | Sal vs. (2*R*,6*R*)-HNK (2.5) | - | - | 0.9996 | - | |
| | | | | | Sal vs. (2*R*,6*R*)-HNK (10) | - | - | 0.9997 | - | |
| | | | Freezing (min 4) | Dunnett | Sal vs. (2*R*,6*R*)-HNK (0.025) | - | - | 0.6477 | - | |
| | | | | | Sal vs. (2*R*,6*R*)-HNK (0.075) | - | - | 0.8288 | - | |
| | | | | | Sal vs. (2*R*,6*R*)-HNK (0.1) | - | - | >0.9999 | - | |
| | | | | | Sal vs. (2*R*,6*R*)-HNK (0.3) | - | - | 0.5981 | - | |
| | | | | | Sal vs. (2*R*,6*R*)-HNK (2.5) | - | - | 0.9447 | - | |
| | | | | | Sal vs. (2*R*,6*R*)-HNK (10) | - | - | 0.1153 | - | |
| | | | Freezing (min 5) | Dunnett | Sal vs. (2*R*,6*R*)-HNK (0.025) | - | - | 0.2481 | - | |
| | | | | | Sal vs. (2*R*,6*R*)-HNK (0.075) | - | - | 0.8924 | - | |
| | | | | | Sal vs. (2*R*,6*R*)-HNK (0.1) | - | - | 0.0006 | ** | |
| | | | | | Sal vs. (2*R*,6*R*)-HNK (0.3) | - | - | 0.9999 | - | |
| | | | | | Sal vs. (2*R*,6*R*)-HNK (2.5) | - | - | 0.9982 | - | |
| | | | | | Sal vs. (2*R*,6*R*)-HNK (10) | - | - | 0.0016 | ** | |
| | | | Freezing (min 1-5) | RMANOVA | Drug | 4.888 | 6,65 | 0.0004 | ** | |
| | | | | | Time | 119.700 | 4,260 | <0.0001 | *** | |
| | | | | | Drug x Time | 2.496 | 24,260 | 0.0002 | ** | |
| | | | | Dunnett | Sal vs. (2*S*,6*S*)-HNK (0.025) | - | - | 0.9486 | - | |
| | | | | | Sal vs. (2*S*,6*S*)-HNK (0.075) | - | - | 0.9996 | - | |
| | | | | | Sal vs. (2*S*,6*S*)-HNK (0.1) | - | - | 0.1596 | - | |
| | | | | | Sal vs. (2*S*,6*S*)-HNK (0.3) | - | - | 0.0067 | ** | |
| | | | | | Sal vs. (2*S*,6*S*)-HNK (2.5) | - | - | 0.4710 | - | |
| | | | | | Sal vs. (2*S*,6*S*)-HNK (10) | - | - | 0.3388 | - | |
| | | | Freezing (min 1) | Dunnett | Sal vs. (2*S*,6*S*)-HNK (0.025) | - | - | >0.9999 | - | |
| | | | | | Sal vs. (2*S*,6*S*)-HNK (0.075) | - | - | >0.9999 | - | |
| | | | | | Sal vs. (2*S*,6*S*)-HNK (0.1) | - | - | 0.9997 | - | |
| | | | | | Sal vs. (2*S*,6*S*)-HNK (0.3) | - | - | 0.9955 | - | |
| | | | | | Sal vs. (2*S*,6*S*)-HNK (2.5) | - | - | 0.9999 | - | |
| | | | | | Sal vs. (2*S*,6*S*)-HNK (10) | - | - | 0.9999 | - | |
| | | | Freezing (min 2) | Dunnett | Sal vs. (2*S*,6*S*)-HNK (0.025) | - | - | >0.9999 | - | |
| | | | | | Sal vs. (2*S*,6*S*)-HNK (0.075) | - | - | >0.9999 | - | |
| | | | | | Sal vs. (2*S*,6*S*)-HNK (0.1) | - | - | 0.9999 | - | |

| | | | | | | | | | | |
|---|---|---|---|---|---|---|---|---|---|---|
| | | | Freezing (min 2) | Dunnett | Sal vs. (2*S*,6*S*)-HNK (0.3) | - | - | >0.9999 | - | S02C |
| | | | | | Sal vs. (2*S*,6*S*)-HNK (2.5) | - | - | 0.9998 | - | |
| | | | | | Sal vs. (2*S*,6*S*)-HNK (10) | - | - | >0.9999 | - | |
| | | | Freezing (min 3) | Dunnett | Sal vs. (2*S*,6*S*)-HNK (0.025) | - | - | 0.9997 | - | |
| | | | | | Sal vs. (2*S*,6*S*)-HNK (0.075) | - | - | >0.9999 | - | |
| | | | | | Sal vs. (2*S*,6*S*)-HNK (0.1) | - | - | 0.9996 | - | |
| | | | | | Sal vs. (2*S*,6*S*)-HNK (0.3) | - | - | 0.9919 | - | |
| | | | | | Sal vs. (2*S*,6*S*)-HNK (2.5) | - | - | 0.9997 | - | |
| | | | | | Sal vs. (2*S*,6*S*)-HNK (10) | - | - | >0.9999 | - | |
| | | | Freezing (min 4) | Dunnett | Sal vs. (2*S*,6*S*)-HNK (0.025) | - | - | 0.3943 | - | |
| | | | | | Sal vs. (2*S*,6*S*)-HNK (0.075) | - | - | 0.9697 | - | |
| | | | | | Sal vs. (2*S*,6*S*)-HNK (0.1) | - | - | 0.0803 | - | |
| | | | | | Sal vs. (2*S*,6*S*)-HNK (0.3) | - | - | 0.0084 | ** | |
| | | | | | Sal vs. (2*S*,6*S*)-HNK (2.5) | - | - | 0.6274 | - | |
| | | | | | Sal vs. (2*S*,6*S*)-HNK (10) | - | - | 0.3784 | - | |
| | | | Freezing (min 5) | Dunnett | Sal vs. (2*S*,6*S*)-HNK (0.025) | - | - | >0.9999 | - | |
| | | | | | Sal vs. (2*S*,6*S*)-HNK (0.075) | - | - | >0.9999 | - | |
| | | | | | Sal vs. (2*S*,6*S*)-HNK (0.1) | - | - | 0.0120 | * | |
| | | | | | Sal vs. (2*S*,6*S*)-HNK (0.3) | - | - | <0.0001 | *** | |
| | | | | | Sal vs. (2*S*,6*S*)-HNK (2.5) | - | - | 0.0564 | - | |
| | | | | | Sal vs. (2*S*,6*S*)-HNK (10) | - | - | 0.0199 | * | |
| | Contextual Fear Conditioning Re-exposure | CFC Re-exposure | (*R,S*)-ketamine Freezing (min 1-5) | RMANOVA | Drug | 0.767 | 3,69 | 0.5166 | - | data not shown |
| | | | | | Time | 16.130 | 4,276 | <0.0001 | *** | |
| | | | | | Drug x Time | 1.891 | 12,276 | 0.0353 | * | |
| | | | (*R,S*)-ketamine Freezing (min 1) | Dunnett | Sal vs. K (2.5) | - | - | >0.9999 | - | |
| | | | | | Sal vs. K (10) | - | - | 0.9991 | - | |
| | | | | | Sal vs. K (30) | - | - | 0.3196 | - | |
| | | | (*R,S*)-ketamine Freezing (min 2) | Dunnett | Sal vs. K (2.5) | - | - | 0.9850 | - | |
| | | | | | Sal vs. K (10) | - | - | 0.6360 | - | |
| | | | | | Sal vs. K (30) | - | - | 0.3682 | - | |
| | | | (*R,S*)-ketamine Freezing (min 3) | Dunnett | Sal vs. K (2.5) | - | - | 0.4097 | - | |
| | | | | | Sal vs. K (10) | - | - | 0.7900 | - | |
| | | | | | Sal vs. K (30) | - | - | 0.7423 | - | |
| | | | (*R,S*)-ketamine Freezing (min 4) | Dunnett | Sal vs. K (2.5) | - | - | 0.1104 | - | |
| | | | | | Sal vs. K (10) | - | - | 0.3964 | - | |
| | | | | | Sal vs. K (30) | - | - | 0.7727 | - | |
| | | | (*R,S*)-ketamine Freezing (min 5) | Dunnett | Sal vs. K (2.5) | - | - | 0.1912 | - | |
| | | | | | Sal vs. K (10) | - | - | 0.2901 | - | |
| | | | | | Sal vs. K (30) | - | - | 0.9875 | - | |
| | | | (2*R*,6*R*)-HNK Freezing (min 1-5) | RMANOVA | Drug | 1.815 | 6,67 | 0.1093 | - | |
| | | | | | Time | 30.210 | 4,268 | <0.0001 | *** | |
| | | | | | Drug x Time | 1.359 | 24,268 | 0.1266 | - | |
| | | | (2*S*,6*S*)-HNK Freezing (min 1-5) | RMANOVA | Drug | 2.939 | 6,65 | 0.0134 | * | |
| | | | | | Time | 42.050 | 4,260 | <0.0001 | *** | |
| | | | | | Drug x Time | 0.793 | 24,260 | 0.7369 | - | |
| | | | | Dunnett | Sal vs. (2*S*,6*S*)-HNK (0.025) | - | - | 0.7553 | - | |
| | | | | | Sal vs. (2*S*,6*S*)-HNK (0.075) | - | - | 0.1069 | - | |
| | | | | | Sal vs. (2*S*,6*S*)-HNK (0.1) | - | - | 0.9998 | - | |
| | | | | | Sal vs. (2*S*,6*S*)-HNK (0.3) | - | - | 0.0221 | * | |
| | | | | | Sal vs. (2*S*,6*S*)-HNK (2.5) | - | - | 0.9955 | - | |
| | | | | | Sal vs. (2*S*,6*S*)-HNK (10) | - | - | 0.7497 | - | |
| | | | (*R,S*)-ketamine Immobility Time (min 1-6) | RMANOVA | Drug | 1.691 | 3,69 | 0.1771 | - | S02D |
| | | | | | Time | 117.400 | 5,345 | <0.0001 | *** | |
| | | | | | Drug x Time | 1.267 | 15,345 | 0.2208 | - | |

| | | | | | | | | | | |
|---|---|---|---|---|---|---|---|---|---|---|
| 1 week prophylactic drug, female | Forced Swim Test Day 1 | FST Day 1 | (2*R*,6*R*)-HNK Immobility Time (min 1-6) | RMANOVA | Drug | 1.727 | 6,65 | 0.1288 | - | S02E |
| | | | | | Time | 5325.000 | 70 | <0.0001 | *** | |
| | | | | | Drug x Time | 3.305 | 30,325 | <0.0001 | *** | |
| | | | (2*R*,6*R*)-HNK Immobility Time (min 1) | Dunnett | Sal vs. (2*R*,6*R*)-HNK (0.025) | - | - | 0.9995 | - | |
| | | | | | Sal vs. (2*R*,6*R*)-HNK (0.075) | - | - | 0.0002 | ** | |
| | | | | | Sal vs. (2*R*,6*R*)-HNK (0.1) | - | - | 0.9996 | - | |
| | | | | | Sal vs. (2*R*,6*R*)-HNK (0.3) | - | - | 0.0133 | * | |
| | | | | | Sal vs. (2*R*,6*R*)-HNK (2.5) | - | - | 0.0003 | ** | |
| | | | | | Sal vs. (2*R*,6*R*)-HNK (10) | - | - | 0.0117 | * | |
| | | | (2*R*,6*R*)-HNK Immobility Time (min 2) | Dunnett | Sal vs. (2*R*,6*R*)-HNK (0.025) | - | - | 0.8126 | - | |
| | | | | | Sal vs. (2*R*,6*R*)-HNK (0.075) | - | - | 0.0836 | - | |
| | | | | | Sal vs. (2*R*,6*R*)-HNK (0.1) | - | - | 0.9996 | - | |
| | | | | | Sal vs. (2*R*,6*R*)-HNK (0.3) | - | - | 0.9996 | - | |
| | | | | | Sal vs. (2*R*,6*R*)-HNK (2.5) | - | - | 0.6335 | - | |
| | | | | | Sal vs. (2*R*,6*R*)-HNK (10) | - | - | >0.9999 | - | |
| | | | (2*R*,6*R*)-HNK Immobility Time (min 3) | Dunnett | Sal vs. (2*R*,6*R*)-HNK (0.025) | - | - | 0.9001 | - | |
| | | | | | Sal vs. (2*R*,6*R*)-HNK (0.075) | - | - | 0.8511 | - | |
| | | | | | Sal vs. (2*R*,6*R*)-HNK (0.1) | - | - | 0.4844 | - | |
| | | | | | Sal vs. (2*R*,6*R*)-HNK (0.3) | - | - | >0.9999 | - | |
| | | | | | Sal vs. (2*R*,6*R*)-HNK (2.5) | - | - | 0.9998 | - | |
| | | | | | Sal vs. (2*R*,6*R*)-HNK (10) | - | - | 0.6174 | - | |
| | | | (2*R*,6*R*)-HNK Immobility Time (min 4) | Dunnett | Sal vs. (2*R*,6*R*)-HNK (0.025) | - | - | 0.7094 | - | |
| | | | | | Sal vs. (2*R*,6*R*)-HNK (0.075) | - | - | 0.9025 | - | |
| | | | | | Sal vs. (2*R*,6*R*)-HNK (0.1) | - | - | 0.3133 | - | |
| | | | | | Sal vs. (2*R*,6*R*)-HNK (0.3) | - | - | 0.6490 | - | |
| | | | | | Sal vs. (2*R*,6*R*)-HNK (2.5) | - | - | 0.9999 | - | |
| | | | | | Sal vs. (2*R*,6*R*)-HNK (10) | - | - | 0.5278 | - | |
| | | | (2*R*,6*R*)-HNK Immobility Time (min 5) | Dunnett | Sal vs. (2*R*,6*R*)-HNK (0.025) | - | - | 0.8150 | - | |
| | | | | | Sal vs. (2*R*,6*R*)-HNK (0.075) | - | - | 0.8135 | - | |
| | | | | | Sal vs. (2*R*,6*R*)-HNK (0.1) | - | - | 0.3201 | - | |
| | | | | | Sal vs. (2*R*,6*R*)-HNK (0.3) | - | - | 0.3838 | - | |
| | | | | | Sal vs. (2*R*,6*R*)-HNK (2.5) | - | - | 0.9404 | - | |
| | | | | | Sal vs. (2*R*,6*R*)-HNK (10) | - | - | 0.6322 | - | |
| | | | (2*R*,6*R*)-HNK Immobility Time (min 6) | Dunnett | Sal vs. (2*R*,6*R*)-HNK (0.025) | - | - | 0.5398 | - | |
| | | | | | Sal vs. (2*R*,6*R*)-HNK (0.075) | - | - | 0.9982 | - | |
| | | | | | Sal vs. (2*R*,6*R*)-HNK (0.1) | - | - | 0.8091 | - | |
| | | | | | Sal vs. (2*R*,6*R*)-HNK (0.3) | - | - | 0.9996 | - | |
| | | | | | Sal vs. (2*R*,6*R*)-HNK (2.5) | - | - | 0.9978 | - | |
| | | | | | Sal vs. (2*R*,6*R*)-HNK (10) | - | - | 0.1556 | - | |
| | | | (2*S*,6*S*)-HNK Immobility Time (min 1-6) | RMANOVA | Drug | 2.255 | 6,65 | 0.0487 | * | |
| | | | | | Time | 66.080 | 5,325 | <0.0001 | *** | |
| | | | | | Drug x Time | 4.627 | 30,325 | <0.0001 | *** | |
| | | | | Dunnett | Sal vs. (2*S*,6*S*)-HNK (0.025) | - | - | 0.5788 | - | |
| | | | | | Sal vs. (2*S*,6*S*)-HNK (0.075) | - | - | 0.2040 | - | |
| | | | | | Sal vs. (2*S*,6*S*)-HNK (0.1) | - | - | 0.0819 | - | |
| | | | | | Sal vs. (2*S*,6*S*)-HNK (0.3) | - | - | 0.0292 | * | |
| | | | | | Sal vs. (2*S*,6*S*)-HNK (2.5) | - | - | 0.9167 | - | |
| | | | | | Sal vs. (2*S*,6*S*)-HNK (10) | - | - | 0.9999 | - | |
| | | | (2*S*,6*S*)-HNK Immobility Time (min 1) | Dunnett | Sal vs. (2*S*,6*S*)-HNK (0.025) | - | - | <0.0001 | *** | |
| | | | | | Sal vs. (2*S*,6*S*)-HNK (0.075) | - | - | 0.0033 | ** | |
| | | | | | Sal vs. (2*S*,6*S*)-HNK (0.1) | - | - | <0.0001 | *** | |
| | | | | | Sal vs. (2*S*,6*S*)-HNK (0.3) | - | - | 0.0005 | ** | |

| | | | | | | | | | | |
|---|---|---|---|---|---|---|---|---|---|---|
| | | | | | Sal vs. (2*S*,6*S*)-HNK (2.5) | - | - | 0.0002 | ** | |
| | | | | | Sal vs. (2*S*,6*S*)-HNK (10) | - | - | <0.0001 | *** | |
| | | | (2*S*,6*S*)-HNK Immobility Time (min 2) | Dunnett | Sal vs. (2*S*,6*S*)-HNK (0.025) | - | - | 0.1668 | - | |
| | | | | | Sal vs. (2*S*,6*S*)-HNK (0.075) | - | - | 0.8193 | - | |
| | | | | | Sal vs. (2*S*,6*S*)-HNK (0.1) | - | - | 0.0545 | - | |
| | | | | | Sal vs. (2*S*,6*S*)-HNK (0.3) | - | - | 0.0048 | ** | |
| | | | | | Sal vs. (2*S*,6*S*)-HNK (2.5) | - | - | 0.2325 | - | |
| | | | | | Sal vs. (2*S*,6*S*)-HNK (10) | - | - | 0.9798 | - | |
| | | | (2*S*,6*S*)-HNK Immobility Time (min 3) | Dunnett | Sal vs. (2*S*,6*S*)-HNK (0.025) | - | - | 0.9996 | - | S02F |
| | | | | | Sal vs. (2*S*,6*S*)-HNK (0.075) | - | - | 0.7739 | - | |
| | | | | | Sal vs. (2*S*,6*S*)-HNK (0.1) | - | - | 0.9946 | - | |
| | | | | | Sal vs. (2*S*,6*S*)-HNK (0.3) | - | - | 0.2804 | - | |
| | | | | | Sal vs. (2*S*,6*S*)-HNK (2.5) | - | - | 0.9982 | - | |
| | | | | | Sal vs. (2*S*,6*S*)-HNK (10) | - | - | 0.7396 | - | |
| | | | (2*S*,6*S*)-HNK Immobility Time (min 4) | Dunnett | Sal vs. (2*S*,6*S*)-HNK (0.025) | - | - | 0.9999 | - | |
| | | | | | Sal vs. (2*S*,6*S*)-HNK (0.075) | - | - | 0.7425 | - | |
| | | | | | Sal vs. (2*S*,6*S*)-HNK (0.1) | - | - | 0.9907 | - | |
| | | | | | Sal vs. (2*S*,6*S*)-HNK (0.3) | - | - | 0.5474 | - | |
| | | | | | Sal vs. (2*S*,6*S*)-HNK (2.5) | - | - | 0.9474 | - | |
| | | | | | Sal vs. (2*S*,6*S*)-HNK (10) | - | - | 0.7460 | - | |
| | | | (2*S*,6S)-HNK Immobility Time (min 5) | Dunnett | Sal vs. (2*S*,6*S*)-HNK (0.025) | - | - | >0.9999 | - | |
| | | | | | Sal vs. (2*S*,6*S*)-HNK (0.075) | - | - | 0.7651 | - | |
| | | | | | Sal vs. (2*S*,6*S*)-HNK (0.1) | - | - | 0.8777 | - | |
| | | | | | Sal vs. (2*S*,6*S*)-HNK (0.3) | - | - | 0.7740 | - | |
| | | | | | Sal vs. (2*S*,6*S*)-HNK (2.5) | - | - | 0.9579 | - | |
| | | | | | Sal vs. (2*S*,6*S*)-HNK (10) | - | - | 0.9404 | - | |
| | | | (2*S*,6*S*)-HNK Immobility Time (min 6) | Dunnett | Sal vs. (2*S*,6*S*)-HNK (0.025) | - | - | 0.9911 | - | |
| | | | | | Sal vs. (2*S*,6*S*)-HNK (0.075) | - | - | 0.4048 | - | |
| | | | | | Sal vs. (2*S*,6*S*)-HNK (0.1) | - | - | >0.9999 | - | |
| | | | | | Sal vs. (2*S*,6*S*)-HNK (0.3) | - | - | 0.3391 | - | |
| | | | | | Sal vs. (2*S*,6*S*)-HNK (2.5) | - | - | 0.9825 | - | |
| | | | | | Sal vs. (2*S*,6*S*)-HNK (10) | - | - | 0.2796 | - | |
| | | | (*R,S*)-ketamine Immobility Time (min 1-6) | RMANOVA | Drug | 4.689 | 3,69 | 0.0049 | ** | |
| | | | | | Time | 25.580 | 5,345 | < 0.0001 | *** | |
| | | | | | Drug x Time | 0.807 | 15,345 | 0.6695 | - | |
| | | | | Dunnett | Sal vs. K (2.5) | - | - | 0.8140 | - | |
| | | | | | Sal vs. K (10) | - | - | 0.0089 | ** | |
| | | | | | Sal vs. K (30) | - | - | 0.2291 | - | |
| | | | (2*R*,6*R*)-HNK Immobility Time (min 1-6) | RMANOVA | Drug | 2.995 | 6,67 | 0.0119 | * | |
| | | | | | Time | 30.680 | 5,335 | < 0.0001 | *** | |
| | | | | | Drug x Time | 1.633 | 30,335 | 0.0217 | * | |
| | | | | Dunnett | Sal vs. (2*R*,6*R*)-HNK (0.025) | - | - | 0.0132 | * | |
| | | | | | Sal vs. (2*R*,6*R*)-HNK (0.075) | - | - | 0.7700 | - | |
| | | | | | Sal vs. (2*R*,6*R*)-HNK (0.1) | - | - | 0.9798 | - | |
| | | | | | Sal vs. (2*R*,6*R*)-HNK (0.3) | - | - | 0.9997 | - | |
| | | | | | Sal vs. (2*R*,6*R*)-HNK (2.5) | - | - | 0.8498 | - | |
| | | | | | Sal vs. (2*R*,6*R*)-HNK (10) | - | - | 0.9933 | - | |
| | | | (2*R*,6*R*)-HNK Immobility Time (min 1) | Dunnett | Sal vs. (2*R*,6*R*)-HNK (0.025) | - | - | 0.0083 | ** | |
| | | | | | Sal vs. (2*R*,6*R*)-HNK (0.075) | - | - | 0.1143 | - | |
| | | | | | Sal vs. (2*R*,6*R*)-HNK (0.1) | - | - | 0.9951 | - | |
| | | | | | Sal vs. (2*R*,6*R*)-HNK (0.3) | - | - | 0.9997 | - | |

| | | | | | | | | | | |
|---|---|---|---|---|---|---|---|---|---|---|
| | Forced Swim Test Day 2 | FST Day 2 | [illegible] | | Sal vs. (2*R*,6*R*)-HNK (2.5) | - | - | 0.1419 | - | data not shown |
| | | | | | Sal vs. (2*R*,6*R*)-HNK (10) | - | - | 0.9996 | - | |
| | | | (2*R*,6*R*)-HNK Immobility Time (min 2) | Dunnett | Sal vs. (2*R*,6*R*)-HNK (0.025) | - | - | 0.3280 | - | |
| | | | | | Sal vs. (2*R*,6*R*)-HNK (0.075) | - | - | 0.6998 | - | |
| | | | | | Sal vs. (2*R*,6*R*)-HNK (0.1) | - | - | 0.1110 | - | |
| | | | | | Sal vs. (2*R*,6*R*)-HNK (0.3) | - | - | 0.9999 | - | |
| | | | | | Sal vs. (2*R*,6*R*)-HNK (2.5) | - | - | 0.6782 | - | |
| | | | | | Sal vs. (2*R*,6*R*)-HNK (10) | - | - | 0.9922 | - | |
| | | | (2*R*,6*R*)-HNK Immobility Time (min 3) | Dunnett | Sal vs. (2*R*,6*R*)-HNK (0.025) | - | - | 0.2657 | - | |
| | | | | | Sal vs. (2*R*,6*R*)-HNK (0.075) | - | - | 0.9998 | - | |
| | | | | | Sal vs. (2*R*,6*R*)-HNK (0.1) | - | - | 0.9684 | - | |
| | | | | | Sal vs. (2*R*,6*R*)-HNK (0.3) | - | - | 0.9997 | - | |
| | | | | | Sal vs. (2*R*,6*R*)-HNK (2.5) | - | - | 0.9996 | - | |
| | | | | | Sal vs. (2*R*,6*R*)-HNK (10) | - | - | 0.9981 | - | |
| | | | (2*R*,6*R*)-HNK Immobility Time (min 4) | Dunnett | Sal vs. (2*R*,6*R*)-HNK (0.025) | - | - | 0.0534 | - | |
| | | | | | Sal vs. (2*R*,6*R*)-HNK (0.075) | - | - | 0.9951 | - | |
| | | | | | Sal vs. (2*R*,6*R*)-HNK (0.1) | - | - | 0.9979 | - | |
| | | | | | Sal vs. (2*R*,6*R*)-HNK (0.3) | - | - | 0.9353 | - | |
| | | | | | Sal vs. (2*R*,6*R*)-HNK (2.5) | - | - | 0.9916 | - | |
| | | | | | Sal vs. (2*R*,6*R*)-HNK (10) | - | - | 0.9723 | - | |
| | | | (2*R*,6*R*)-HNK Immobility Time (min 5) | Dunnett | Sal vs. (2*R*,6*R*)-HNK (0.025) | - | - | 0.0049 | ** | |
| | | | | | Sal vs. (2*R*,6*R*)-HNK (0.075) | - | - | 0.9999 | - | |
| | | | | | Sal vs. (2*R*,6*R*)-HNK (0.1) | - | - | 0.4487 | - | |
| | | | | | Sal vs. (2*R*,6*R*)-HNK (0.3) | - | - | 0.9646 | - | |
| | | | | | Sal vs. (2*R*,6*R*)-HNK (2.5) | - | - | 0.9999 | - | |
| | | | | | Sal vs. (2*R*,6*R*)-HNK (10) | - | - | 0.9686 | - | |
| | | | (2*R*,6*R*)-HNK Immobility Time (min 6) | Dunnett | Sal vs. (2*R*,6*R*)-HNK (0.025) | - | - | 0.0498 | * | |
| | | | | | Sal vs. (2*R*,6*R*)-HNK (0.075) | - | - | 0.7180 | - | |
| | | | | | Sal vs. (2*R*,6*R*)-HNK (0.1) | - | - | 0.4146 | - | |
| | | | | | Sal vs. (2*R*,6*R*)-HNK (0.3) | - | - | 0.9492 | - | |
| | | | | | Sal vs. (2*R*,6*R*)-HNK (2.5) | - | - | 0.9966 | - | |
| | | | | | Sal vs. (2*R*,6*R*)-HNK (10) | - | - | 0.9998 | - | |
| | | | (2*S*,6*S*)-HNK Immobility Time (min 1-6) | RMANOVA | Drug | 2.256 | 6,65 | 0.0487 | * | |
| | | | | | Time | 8.509 | 5,325 | < 0.0001 | *** | |
| | | | | | Drug x Time | 2.843 | 30,325 | < 0.0001 | *** | |
| | | | | Dunnett | Sal vs. (2*S*,6*S*)-HNK (0.025) | - | - | 0.9997 | - | |
| | | | | | Sal vs. (2*S*,6*S*)-HNK (0.075) | - | - | 0.0526 | - | |
| | | | | | Sal vs. (2*S*,6*S*)-HNK (0.1) | - | - | 0.5496 | - | |
| | | | | | Sal vs. (2*S*,6*S*)-HNK (0.3) | - | - | 0.9995 | - | |
| | | | | | Sal vs. (2*S*,6*S*)-HNK (2.5) | - | - | 0.7831 | - | |
| | | | | | Sal vs. (2*S*,6*S*)-HNK (10) | - | - | >0.9999 | - | |
| | | | (2*S*,6*S*)-HNK Immobility Time (min 1) | Dunnett | Sal vs. (2*S*,6*S*)-HNK (0.025) | - | - | 0.9129 | - | |
| | | | | | Sal vs. (2*S*,6*S*)-HNK (0.075) | - | - | 0.9850 | - | |
| | | | | | Sal vs. (2*S*,6*S*)-HNK (0.1) | - | - | 0.0043 | ** | |
| | | | | | Sal vs. (2*S*,6*S*)-HNK (0.3) | - | - | 0.0841 | - | |
| | | | | | Sal vs. (2*S*,6*S*)-HNK (2.5) | - | - | 0.9997 | - | |
| | | | | | Sal vs. (2*S*,6*S*)-HNK (10) | - | - | 0.3257 | - | |
| | | | (2*S*,6*S*)-HNK Immobility Time (min 2) | Dunnett | Sal vs. (2*S*,6*S*)-HNK (0.025) | - | - | 0.9949 | - | |
| | | | | | Sal vs. (2*S*,6*S*)-HNK (0.075) | - | - | 0.9997 | - | |
| | | | | | Sal vs. (2*S*,6*S*)-HNK (0.1) | - | - | 0.3196 | - | |
| | | | | | Sal vs. (2*S*,6*S*)-HNK (0.3) | - | - | 0.5483 | - | |

| | | | | | | | | | | |
|---|---|---|---|---|---|---|---|---|---|---|
| | | | 2) | | Sal vs. (2*S*,6*S*)-HNK (2.5) | - | - | >0.9999 | - | |
| | | | | | Sal vs. (2*S*,6*S*)-HNK (10) | - | - | 0.9824 | - | |
| | | | (2*S*,6*S*)-HNK Immobility Time (min 3) | Dunnett | Sal vs. (2*S*,6*S*)-HNK (0.025) | - | - | 0.9996 | - | |
| | | | | | Sal vs. (2*S*,6*S*)-HNK (0.075) | - | - | 0.9469 | - | |
| | | | | | Sal vs. (2*S*,6*S*)-HNK (0.1) | - | - | 0.9815 | - | |
| | | | | | Sal vs. (2*S*,6*S*)-HNK (0.3) | - | - | 0.9979 | - | |
| | | | | | Sal vs. (2*S*,6*S*)-HNK (2.5) | - | - | 0.6411 | - | |
| | | | | | Sal vs. (2*S*,6*S*)-HNK (10) | - | - | 0.9636 | - | |
| | | | (2*S*,6*S*)-HNK Immobility Time (min 4) | Dunnett | Sal vs. (2*S*,6*S*)-HNK (0.025) | - | - | 0.8990 | - | |
| | | | | | Sal vs. (2*S*,6*S*)-HNK (0.075) | - | - | 0.3811 | - | |
| | | | | | Sal vs. (2*S*,6*S*)-HNK (0.1) | - | - | 0.9996 | - | |
| | | | | | Sal vs. (2*S*,6*S*)-HNK (0.3) | - | - | 0.9997 | - | |
| | | | | | Sal vs. (2*S*,6*S*)-HNK (2.5) | - | - | 0.8244 | - | |
| | | | | | Sal vs. (2*S*,6*S*)-HNK (10) | - | - | 0.9999 | - | |
| | | | (2*S*,6*S*)-HNK Immobility Time (min 5) | Dunnett | Sal vs. (2*S*,6*S*)-HNK (0.025) | - | - | 0.8687 | - | |
| | | | | | Sal vs. (2*S*,6*S*)-HNK (0.075) | - | - | <0.0001 | *** | |
| | | | | | Sal vs. (2*S*,6*S*)-HNK (0.1) | - | - | 0.9659 | - | |
| | | | | | Sal vs. (2*S*,6*S*)-HNK (0.3) | - | - | 0.8648 | - | |
| | | | | | Sal vs. (2*S*,6*S*)-HNK (2.5) | - | - | 0.3928 | - | |
| | | | | | Sal vs. (2*S*,6*S*)-HNK (10) | - | - | 0.8600 | - | |
| | | | (2*S*,6*S*)-HNK Immobility Time (min 6) | Dunnett | Sal vs. (2*S*,6*S*)-HNK (0.025) | - | - | >0.9999 | - | |
| | | | | | Sal vs. (2*S*,6*S*)-HNK (0.075) | - | - | 0.0001 | ** | |
| | | | | | Sal vs. (2*S*,6*S*)-HNK (0.1) | - | - | 0.9105 | - | |
| | | | | | Sal vs. (2*S*,6*S*)-HNK (0.3) | - | - | 0.1993 | - | |
| | | | | | Sal vs. (2*S*,6*S*)-HNK (2.5) | - | - | 0.9788 | - | |
| | | | | | Sal vs. (2*S*,6*S*)-HNK (10) | - | - | 0.9927 | - | |
| Drug levels, brain and plasma | Liquid chromatography mass spectrometry | LC-MS | Brain concentration, (*R,S*)-ketamine | *t*-test | (2*R*,6*R*)-HNK vs. (2*S*,6*S*)-HNK | - | - | 0.823 | - | S03B |
| | | | Plasma concentration, (*R,S*)-ketamine | *t*-test | (2*R*,6*R*)-HNK vs. (2*S*,6*S*)-HNK | - | - | 0.8854 | - | S03C |
| 1 week prophylactic drug, no stress | Contextual Fear Conditioning Training | CFC Training | Freezing (min 1-5) | RMANOVA | Drug | 0.6917 | 2,12 | 0.5196 | - | S04B |
| | | | | | Time | 1.165 | 4,48 | 0.3380 | - | |
| | | | | | Drug x Time | 0.8011 | 8,48 | 0.6046 | - | |
| | Contextual Fear Conditioning Re-exposure | CFC Re-exposure | Freezing (min 1-5) | RMANOVA | Drug | 0.6055 | 2,12 | 0.5616 | - | S04C |
| | | | | | Time | 3.355 | 4,48 | 0.0168 | * | |
| | | | | | Drug x Time | 0.6445 | 8,48 | 0.7364 | - | |
| | | | Freezing Average | ANOVA | Drug | 0.6055 | 2,12 | 0.5616 | - | S04D |
| | Forced Swim Test Day 1 | FST Day 1 | Immobility Time (min 1-6) | RMANOVA | Drug | 2.106 | 2,12 | 0.1645 | - | S04E |
| | | | | | Time | 10.61 | 5,60 | <0.0001 | *** | |
| | | | | | Drug x Time | 1.854 | 10,60 | 0.0702 | - | |
| | | | Immobility Time | ANOVA | Drug | 2.656 | 2,12 | 0.1109 | - | |
| | Forced Swim Test Day 2 | FST Day 2 | Immobility Time (min 1-6) | RMANOVA | Drug | 0.6938 | 2,12 | 0.5187 | - | S04F |
| | | | | | Time | 4.298 | 5,60 | 0.0021 | ** | |
| | | | | | Drug x Time | 0.317 | 10,60 | 0.9737 | - | |
| | | | Immobility Time | ANOVA | Drug | 0.2816 | 2,12 | 0.7594 | - | S04G |
| | | | Time in Open Arms (min 1-6) | RMANOVA | Drug | 5.438 | 2,60 | 0.0208 | * | S05A |
| | | | | | Time | 1.840 | 5,60 | 0.1186 | - | |
| | | | | | Drug x Time | 0.661 | 10,60 | 0.7552 | - | |
| | | | | Dunnett | Sal vs. K (10) | - | - | 0.0269 | * | |

| | | | | | | | | | | |
|---|---|---|---|---|---|---|---|---|---|---|
| 1 week prophylactic drug, learned helplessness stress | Elevated Plus Maze | EPM | | Dunnett | Sal vs. (2*R*,*R* )-HNK (0.025) | - | - | 0.9997 | - | |
| | | | Time in Closed Arms (min 1-6) | RMANOVA | Drug | 3.777 | 2,60 | 0.0534 | - | S05B |
| | | | | | Time | 2.235 | 5,60 | 0.0623 | - | |
| | | | | | Drug x Time | 0.547 | 10,60 | 0.8498 | - | |
| | | | Time in Center (min 1-6) | RMANOVA | Drug | 0.826 | 2,60 | 0.4612 | - | S05C |
| | | | | | Time | 0.783 | 5,60 | 0.5657 | - | |
| | | | | | Drug x Time | 0.645 | 10,60 | 0.7696 | - | |
| | | | Distance Travelled in Open Arms (min 1-6) | RMANOVA | Drug | 2.322 | 2,60 | 0.1404 | - | data not shown |
| | | | | | Distance | 2.373 | 5,60 | 0.0496 | * | |
| | | | | | Drug x Distance | 1.873 | 10,60 | 0.0671 | - | |
| | | | Distance Travelled in Open Arms Average (min 1-6) | ANOVA | Drug | 2.322 | 2,12 | 0.1404 | - | |
| | | | Distance Travelled in Closed Arms (min 1-6) | RMANOVA | Drug | 1.284 | 2,60 | 0.3125 | - | |
| | | | | | Distance | 2.866 | 5,60 | 0.0219 | * | |
| | | | | | Drug x Distance | 1.438 | 10,60 | 0.1860 | - | |
| | | | Distance Travelled in Closed Arms Average (min 1-6) | ANOVA | Drug | 1.284 | 2,12 | 0.3125 | - | |
| | | | Distance Travelled in Center (min 1-6) | RMANOVA | Drug | 0.689 | 2,60 | 0.5207 | - | |
| | | | | | Distance | 1.778 | 5,60 | 0.1310 | - | |
| | | | | | Drug x Distance | 0.658 | 10,60 | 0.7582 | - | |
| | | | Distance Travelled in Center Average (min 1-6) | ANOVA | Drug | 0.689 | 2,12 | 0.5207 | - | |
| | | | Entries into Open Arms Average (min 1-6) | ANOVA | Drug | 3.352 | 2,12 | 0.0697 | - | |
| | | | Entries into Closed Arms Average (min 1-6) | ANOVA | Drug | 0.079 | 2,12 | 0.9249 | - | |
| | | | Entries into Center Average (min 1-6) | ANOVA | Drug | 0.389 | 2,12 | 0.6860 | - | |
| | Contextual Fear Conditioning Training | CFC Training | Freezing (min 1-5) | RMANOVA | Drug | 4.539 | 4,100 | 0.0068 | ** | S06B |
| | | | | | Time | 113.383 | 4,100 | <0.0001 | *** | |
| | | | | | Drug x Time | 3.012 | 16,100 | 0.0004 | ** | |
| | | | | Dunnett | Sal vs. K (2.5) | - | - | 0.0670 | - | |
| | | | | | Sal vs. K (10) | - | - | 0.0359 | * | |
| | | | | | Sal vs. K (30) | - | - | 0.1758 | - | |
| | | | | | Sal vs. (2*R* ,6*R* )-HNK (0.025) | - | - | >0.9999 | - | |
| | | | Freezing (min 1) | Dunnett | Sal vs. K (2.5) | - | - | >0.9999 | - | |
| | | | | | Sal vs. K (10) | - | - | >0.9999 | - | |
| | | | | | Sal vs. K (30) | - | - | >0.9999 | - | |
| | | | | | Sal vs. (2*R* ,6*R* )-HNK (0.025) | - | - | >0.9999 | - | |
| | | | Freezing (min 2) | Dunnett | Sal vs. K (2.5) | - | - | >0.9999 | - | |
| | | | | | Sal vs. K (10) | - | - | 0.9998 | - | |
| | | | | | Sal vs. K (30) | - | - | 0.9910 | - | |
| | | | | | Sal vs. (2*R* ,6*R* )-HNK (0.025) | - | - | 0.9999 | - | |
| | | | Freezing (min 3) | Dunnett | Sal vs. K (2.5) | - | - | 0.9972 | - | |
| | | | | | Sal vs. K (10) | - | - | 0.9999 | - | |
| | | | | | Sal vs. K (30) | - | - | 0.9983 | - | |

| | | | | | | | | | | |
|---|---|---|---|---|---|---|---|---|---|---|
| 3 day prophylactic drug | | | | | Sal vs. (2*R*,6*R*)-HNK (0.025) | - | - | 0.9928 | - | |
| | | | Freezing (min 4) | Dunnett | Sal vs. K (2.5) | - | - | 0.0125 | * | |
| | | | | | Sal vs. K (10) | - | - | 0.0619 | - | |
| | | | | | Sal vs. K (30) | - | - | 0.3038 | - | |
| | | | | | Sal vs. (2*R*,6*R*)-HNK (0.025) | - | - | 0.9997 | - | |
| | | | Freezing (min 5) | Dunnett | Sal vs. K (2.5) | - | - | <0.0001 | *** | |
| | | | | | Sal vs. K (10) | - | - | 0.0002 | ** | |
| | | | | | Sal vs. K (30) | - | - | 0.0068 | ** | |
| | | | | | Sal vs. (2*R*,6*R*)-HNK (0.025) | - | - | >0.9999 | - | |
| | Contextual Fear Conditioning Re-exposure | CFC Re-exposure | Freezing (min 1-5) | RMANOVA | Drug | 2.041 | 4,100 | 0.1193 | - | S06C |
| | | | | | Time | 11.504 | 4,100 | <0.0001 | *** | |
| | | | | | Drug x Time | 1.163 | 16,100 | 0.3110 | - | |
| | | | Freezing Average | ANOVA | Drug | 2.041 | 4,25 | 0.1193 | - | S06D |
| | Forced Swim Test Day 1 | FST Day 1 | Immobility Time (min 1-6) | RMANOVA | Drug | 11.903 | 4,125 | <0.0001 | *** | S06E |
| | | | | | Time | 15.387 | 5,125 | <0.0001 | *** | |
| | | | | | Drug x Time | 6.709 | 20,125 | <0.0001 | *** | |
| | | | | Dunnett | Sal vs. K (2.5) | - | - | 0.9916 | - | |
| | | | | | Sal vs. K (10) | - | - | 0.9999 | - | |
| | | | | | Sal vs. K (30) | - | - | >0.9999 | - | |
| | | | | | Sal vs. (2*R*,6*R*)-HNK (0.025) | - | - | 0.0003 | ** | |
| | | | Immobility Time (min 1) | Dunnett | Sal vs. K (2.5) | - | - | 0.9998 | - | |
| | | | | | Sal vs. K (10) | - | - | 0.9998 | - | |
| | | | | | Sal vs. K (30) | - | - | 0.8877 | - | |
| | | | | | Sal vs. (2*R*,6*R*)-HNK (0.025) | - | - | <0.0001 | *** | |
| | | | Immobility Time (min 2) | Dunnett | Sal vs. K (2.5) | - | - | 0.9397 | - | |
| | | | | | Sal vs. K (10) | - | - | 0.9294 | - | |
| | | | | | Sal vs. K (30) | - | - | 0.9999 | - | |
| | | | | | Sal vs. (2*R*,6*R*)-HNK (0.025) | - | - | <0.0001 | ** | |
| | | | Immobility Time (min 3) | Dunnett | Sal vs. K (2.5) | - | - | 0.9557 | - | |
| | | | | | Sal vs. K (10) | - | - | 0.9968 | - | |
| | | | | | Sal vs. K (30) | - | - | 0.9979 | - | |
| | | | | | Sal vs. (2*R*,6*R*)-HNK (0.025) | - | - | 0.0658 | ** | |
| | | | Immobility Time (min 4) | Dunnett | Sal vs. K (2.5) | - | - | >0.9999 | - | |
| | | | | | Sal vs. K (10) | - | - | 0.9759 | - | |
| | | | | | Sal vs. K (30) | - | - | 0.9489 | - | |
| | | | | | Sal vs. (2*R*,6*R*)-HNK (0.025) | - | - | 0.0261 | ** | |
| | | | Immobility Time (min 5) | Dunnett | Sal vs. K (2.5) | - | - | 0.9918 | - | |
| | | | | | Sal vs. K (10) | - | - | >0.9999 | - | |
| | | | | | Sal vs. K (30) | - | - | 0.9999 | - | |
| | | | | | Sal vs. (2*R*,6*R*)-HNK (0.025) | - | - | 0.0100 | ** | |
| | | | Immobility Time (min 6) | Dunnett | Sal vs. K (2.5) | - | - | 0.9965 | - | |
| | | | | | Sal vs. K (10) | - | - | >0.9999 | - | |
| | | | | | Sal vs. K (30) | - | - | 0.9853 | - | |
| | | | | | Sal vs. (2*R*,6*R*)-HNK (0.025) | - | - | 0.0148 | ** | |
| | | | Immobility Time Average (min 3-6) | ANOVA | Drug | 7.524 | 4,5 | 0.0004 | ** | |
| | | | | Dunnett | Sal vs. K (2.5) | - | - | 0.9825 | - | |
| | | | | | Sal vs. K (10) | - | - | >0.9999 | - | |
| | | | | | Sal vs. K (30) | - | - | 0.9984 | - | |
| | | | | | Sal vs. (2*R*,6*R*)-HNK (0.025) | - | - | 0.0029 | ** | |
| | | | Immobility Time (min 1-6) | RMANOVA | Drug | 14.210 | 4,25 | <0.0001 | *** | S06F |
| | | | | | Time | 3.300 | 5,125 | 0.0078 | ** | |
| | | | | | Drug x Time | 0.512 | 20,125 | 0.0883 | - | |
| | | | | | Sal vs. K (2.5) | - | - | 0.7580 | - | |

| | | | | | | | | | | |
|---|---|---|---|---|---|---|---|---|---|---|
| | Forced Swim Test Day 2 | FST Day 2 | 1-6) | Dunnett | Sal vs. K (10) | - | - | 0.8071 | - | |
| | | | | | Sal vs. K (30) | - | - | 0.6191 | - | |
| | | | | | Sal vs. (2*R*,6*R*)-HNK (0.025) | - | - | 0.0007 | ** | |
| | | | Immobility Time Average (min 3-6) | ANOVA | Drug | 7.868 | 4,25 | 0.0003 | ** | S06G |
| | | | | Dunnett | Sal vs. K (2.5) | - | - | 0.8054 | - | |
| | | | | | Sal vs. K (10) | - | - | 0.8470 | - | |
| | | | | | Sal vs. K (30) | - | - | 0.6822 | - | |
| | | | | | Sal vs. (2*R*,6*R*)-HNK (0.025) | - | - | 0.0161 | * | |
| 24 hour prophylactic drug | Contextual Fear Conditioning | CFC Training | Freezing (min 1-5) | RMANOVA | Drug | 1.400 | 1,72 | 0.2522 | - | S07B |
| | | | | | Time | 80.926 | 4,72 | <0.0001 | *** | |
| | | | | | Drug x Time | 2.019 | 4,72 | 0.1008 | - | |
| | Contextual Fear Conditioning Re-exposure | CFC Re-exposure | Freezing (min 1-5) | RMANOVA | Drug | 1.527 | 1,68 | 0.2334 | - | S07C |
| | | | | | Time | 6.065 | 4,68 | 0.0003 | ** | |
| | | | | | Drug x Time | 0.216 | 4,68 | 0.9289 | - | |
| | | | Freezing Average | *t*-test | Sal vs. (2*R*,6*R*)-HNK (0.025) | - | - | 0.2334 | - | S07D |
| | Forced Swim Test Day 1 | FST Day 1 | Immobility Time (min 1-6) | RMANOVA | Drug | 0.349 | 1,85 | 0.5622 | - | S07E |
| | | | | | Time | 14.149 | 5,85 | <0.0001 | *** | |
| | | | | | Drug x Time | 0.394 | 5,85 | 0.8516 | - | |
| | | | Immobility Time | *t*-test | Sal vs. (2*R*,6*R*)-HNK (0.025) | - | - | 0.5779 | - | |
| | Forced Swim Test Day 2 | FST Day 2 | Immobility Time (min 1-6) | RMANOVA | Drug | 2.102 | 1,17 | 0.1653 | - | S07F |
| | | | | | Time | 6.780 | 5,85 | <0.0001 | *** | |
| | | | | | Drug x Time | 1.123 | 5,85 | 0.3544 | - | |
| | | | Immobility Time | *t*-test | Sal vs. (2*R*,6*R*)-HNK (0.025) | - | - | 0.1367 | - | S07G |
| No stress, antidepressant drug | Forced Swim Test Day 1 | FST Day 1 | Immobility Time (min 1-6) | RMANOVA | Drug | 0.095 | 2,12 | 0.9102 | - | S08B |
| | | | | | Time | 14.724 | 5,60 | <0.0001 | *** | |
| | | | | | Drug x Time | 1.257 | 10,60 | 0.2749 | - | |
| | | | Immobility Time Average (min 3-6) | ANOVA | Drug | 0.294 | 2,12 | 0.7503 | - | data not shown |
| | Forced Swim Test Day 2 | FST Day 2 | Immobility Time (min 1-6) | RMANOVA | Drug | 1.136 | 2,60 | 0.3532 | - | S08C |
| | | | | | Time | 20.037 | 5,60 | <0.0001 | *** | |
| | | | | | Drug x Time | 1.709 | 10,60 | 0.0996 | - | |
| | | | Immobility Time | ANOVA | Drug | 0.577 | 2,10 | 0.5763 | - | S08D |
| Stress, antidepressant drug | Forced Swim Test Day 1 | FST Day 1 | Immobility Time (min 1-6) | RMANOVA | Drug | 0.088 | 2,50 | 0.9166 | - | S09B |
| | | | | | Time | 16.293 | 5,50 | <.0001 | *** | |
| | | | | | Drug x Time | 0.548 | 10,50 | 0.8473 | - | |
| | | | Immobility Time Average (min 3-6) | ANOVA | Drug | 0.047 | 2,10 | 0.9540 | - | data not shown |
| | Forced Swim Test Day 2 | FST Day 2 | Immobility Time (min 1-6) | RMANOVA | Drug | 0.703 | 2,10 | 0.5182 | - | S09C |
| | | | | | Time | 4.107 | 5,50 | 0.0033 | ** | |
| | | | | | Drug x Time | 0.923 | 10,50 | 0.5207 | - | |
| | | | Immobility Time | ANOVA | Drug | 0.234 | 2,10 | 0.7957 | - | S09D |
| | Contextual Fear Conditioning Training | CFC Training | Freezing (min 1-5) | RMANOVA | Drug | 3.866 | 2,40 | 0.0570 | - | data not shown |
| | | | | | Time | 79.288 | 4,40 | <.0001 | *** | |
| | | | | | Drug x Time | 3.473 | 8,40 | 0.0039 | ** | |
| | | | Freezing (min 1) | Dunnett | Sal vs. K (10) | - | - | 0.8425 | - | |
| | | | | | Sal vs. (2*R*,6*R*)-HNK (0.025) | - | - | 0.8504 | - | |
| | | | Freezing (min 2) | Dunnett | Sal vs. K (10) | - | - | 0.7779 | - | |
| | | | | | Sal vs. (2*R*,6*R*)-HNK (0.025) | - | - | 0.8142 | - | |
| | | | Freezing (min 3) | Dunnett | Sal vs. K (10) | - | - | 0.7068 | - | |
| | | | | | Sal vs. (2*R*,6*R*)-HNK (0.025) | - | - | 0.9251 | - | |

| | | | | | | | | | | |
|---|---|---|---|---|---|---|---|---|---|---|
| | | | Freezing (min 4) | Dunnett | Sal vs. K (10) | - | - | 0.0196 | * | |
| | | | | | Sal vs. (2*R* ,6*R* )-HNK (0.025) | - | - | 0.1372 | - | |
| | | | Freezing (min 5) | Dunnett | Sal vs. K (10) | - | - | 0.1230 | - | |
| | | | | | Sal vs. (2*R* ,6*R* )-HNK (0.025) | - | - | 0.0106 | * | |
| | Contextual Fear Conditioning Re-exposure | CFC Re-exposure | Freezing (min 1-5) | RMANOVA | Drug | 0.129 | 2,40 | 0.8804 | - | S09E |
| | | | | | Time | 2.908 | 4,40 | 0.0334 | * | |
| | | | | | Drug x Time | 0.879 | 8,40 | 0.5420 | - | |
| | | | Freezing Average | ANOVA | Drug | 0.129 | 2,10 | 0.8804 | - | S09F |
